# Inferring gene regulatory networks by hypergraph variational autoencoder

**DOI:** 10.1101/2024.04.01.586509

**Authors:** Guangxin Su, Hanchen Wang, Ying Zhang, Adelle CF Coster, Marc R. Wilkins, Pablo F. Canete, Di Yu, Yang Yang, Wenjie Zhang

## Abstract

In constructing Gene Regulatory Networks (GRNs), it is crucial to consider cellular heterogeneity and differential gene regulatory modules. However, traditional methods have predominantly focused on cellular heterogeneity, approaching the subject from a relatively narrow scope. We present HyperG-VAE, a Bayesian deep generative model that utilizes a hypergraph to model single-cell RNA sequencing (scRNA-seq) data. HyperG-VAE employs a cell encoder with a Structural Equation Model to address cellular heterogeneity and build GRNs, alongside a gene encoder using hypergraph self-attention to identify gene modules. Encoders are synergistically optimized by a decoder, enabling HyperG-VAE to excel in GRN inference, single-cell clustering, and data visualization, evidenced by benchmarks. Additionally, HyperG-VAE effectively reveals gene regulation patterns and shows robustness in varied downstream analyses, demonstrated using B cell development data in bone marrow. The interplay of encoders by the overlapping genes between predicted GRNs and gene modules is further validated by gene set enrichment analysis, underscoring that the gene encoder boosts the GRN inference. HyperG-VAE proves efficient in scRNA-seq data analysis and GRN inference.

## 1 Introduction

Gene Regulatory Networks (GRNs) within single-cell RNA sequencing (scRNA-seq) datasets present a sophisticated interplay of transcription factors (TFs) and target genes, uniquely capturing the modulation of gene expression and thereby delineating the intricate cellular functions and responses within diverse cell populations [1]. GRNs illuminate core biological processes and underpin applications from disease modeling to therapeutic design [2, 3, 4], empowering researchers to interpret the mechanisms of gene interactions within cells and leverage this understanding for medical and biotechnological innovations [5, 6].

Numerous methodologies have emerged for inferring GRNs from single-cell transcriptomic data. The algorithms emphasize co-expression networks in a statistical way (e.g., PPCOR [7] and LocaTE [8]) or aim to decipher causal relationships between TFs and their target genes based on the analysis of the gene interactions among cells (e.g., DeepSEM [9] and PIDC [10]). Despite their achievements, these algorithms still have inherent limitations. Specifically, these approaches mainly focus on cellular heterogeneity and overlook the critical importance of simultaneously considering cellular heterogeneity and gene module information in the model design. Generally, from the view of underlying principles, we can divide the methodologies into deep learning methods and traditional statistical algorithms. Many deep learning (e.g., DeepTFni [11] and DeepSEM [9]) based methodologies primarily build upon foundational models [12, 13]. The frequent oversight in these models is the inherent relationships between cells and genes, as informed by domain expertise. This often leads to models that compromise on explainability and narrow their application scope. For the traditional statistical algorithms, such as Bayesian networks [14, 15] and ensemble methods [16, 17, 18], it can be computationally expensive, and it remains a challenge to extend these methodologies to encompass broader nonlinear paradigms.

Additionally, the scRNA-seq data is frequently marred by noise and incompleteness, attributable to phenomena such as amplification biases inherent to reverse transcription and PCR amplification processes [19, 20], as well as the issue of low quantities of nucleic acids in single cells [21]. To get a more robust GRN, several methodologies [22, 23] leverage multi-omic datasets, capturing kinds of cellular information to enrich the model’s comprehensiveness. However, integrating multi-omic datasets presents substantial challenges, particularly regarding harmonizing data from disparate sources and platforms and could also introduce additional noise [24].

To address the problems and construct a reliable GRN, we model scRNA-seq data as a hypergraph and present Hypergraph Variational Autoencoder (HyperG-VAE), a Bayesian deep generative model to process the hypergraph data. Distinct from current approaches, HyperG-VAE simultaneously captures cellular heterogeneity and gene modules ^1^ through its cell and gene encoders individually during the GRNs construction. Two encoders employ variational inference to learn stochastic representations of genes and cells, offering a more flexible and robust approach to manage real-world data complexities. This could be particularly effective in handling noise in scRNA-seq datasets, a capability that has been demonstrated in previous studies [25, 26]. Within a shared embedding space, the dual encoders of our model interact, boosting its cohesiveness. The joint optimization manner elucidates gene regulatory mechanisms within gene modules across various cell clusters, thereby augmenting the model’s ability to delineate complex gene regulatory interactions and significantly improving its explainability.

Our study evaluates the performance of HyperG-VAE in various scRNA-seq applications. These include i. GRN inference, ii. cell embedding, iii. gene embedding, and iv. gene regulation hypergraph construction. Through benchmark comparisons, encompassing tasks like GRNs inference, data visualization, and single-cell clustering, we establish that HyperG-VAE outperforms existing state-of-the-art methods. Additionally, HyperG-VAE demonstrates its utility in elucidating the regulatory patterns governing B cell development in bone marrow. Our model also excels in learning gene expression modules and cell clusters, which connect the gene encoder and cell encoder individually to boost gene regulatory hypergraph prediction. This integrated functionality of HyperG-VAE improves our comprehension of single-cell transcriptomic data, ultimately providing better insights into the realm of GRNs inference.

## 2 Results

### Framework overview

We introduce HyperG-VAE, a Bayesian deep generative model specifically designed to address the complex challenge of gene regulation network inference using scRNA-seq data, which is represented as a hypergraph (Fig.1 and Methods). Our HyperG-VAE takes into account the interplay between gene modules and cellular heterogeneity, allowing for a more accurate representation of cell-specific regulatory mechanisms. This interplay could be incorporated in a hypergraph to capture the nuanced interactions of genes across diverse cellular states. In the hypergraph, we conceptualize genes expressed within individual cells as nodes, interconnected through unique hyperedges (cells) (Fig.1b).

HyperG-VAE incorporates two encoders: a cell encoder and a gene encoder, enabling it to learn the hypergraph representation ***H***^***ν***^ (Fig.1c). The cell encoder generates cell representations ***H***^***ε***^ in the form of hypergraph duality, facilitating the embedding of high-order relations via structural equation layers. GRN construction (Fig.1d) is realized in this structural equation layers through a learnable causal interaction matrix. In addition, the cell encoder can adeptly capture the gene regulation process in a cell-specific manner, elucidating a clearer landscape of cellular heterogeneity (Fig.1e). The gene encoder is specifically designed to process observed gene representations, denoted as ***H***^***ν***^. Given that genes within a module generally manifest consistent expression profiles across cells, we employs a multi-head self-attention mechanism that is specifically designed for hypergraph in this work. This not only discerns varying gene expression levels but also assigns appropriate weights to the genes expressed in the same cell during the message-passing phase. Thus, the gene encoder enhances the model’s ability to understand and integrate the intricate interdependencies among genes, thereby aiding in the effective embedding of gene clusters (Fig.1f). Finally, a hypergraph decoder is utilized to reconstruct the original topology of the hypergraph (Fig.1g) using the learned latent embedding of genes and cells. Utilizing the reconstructed hypergraph and the learned inter-gene relationships, we can also infer a gene regulatory hypergraph (Fig.1g). This hypergraph encompasses gene regulatory modules that span across various cell stages.

**Fig. 1.**
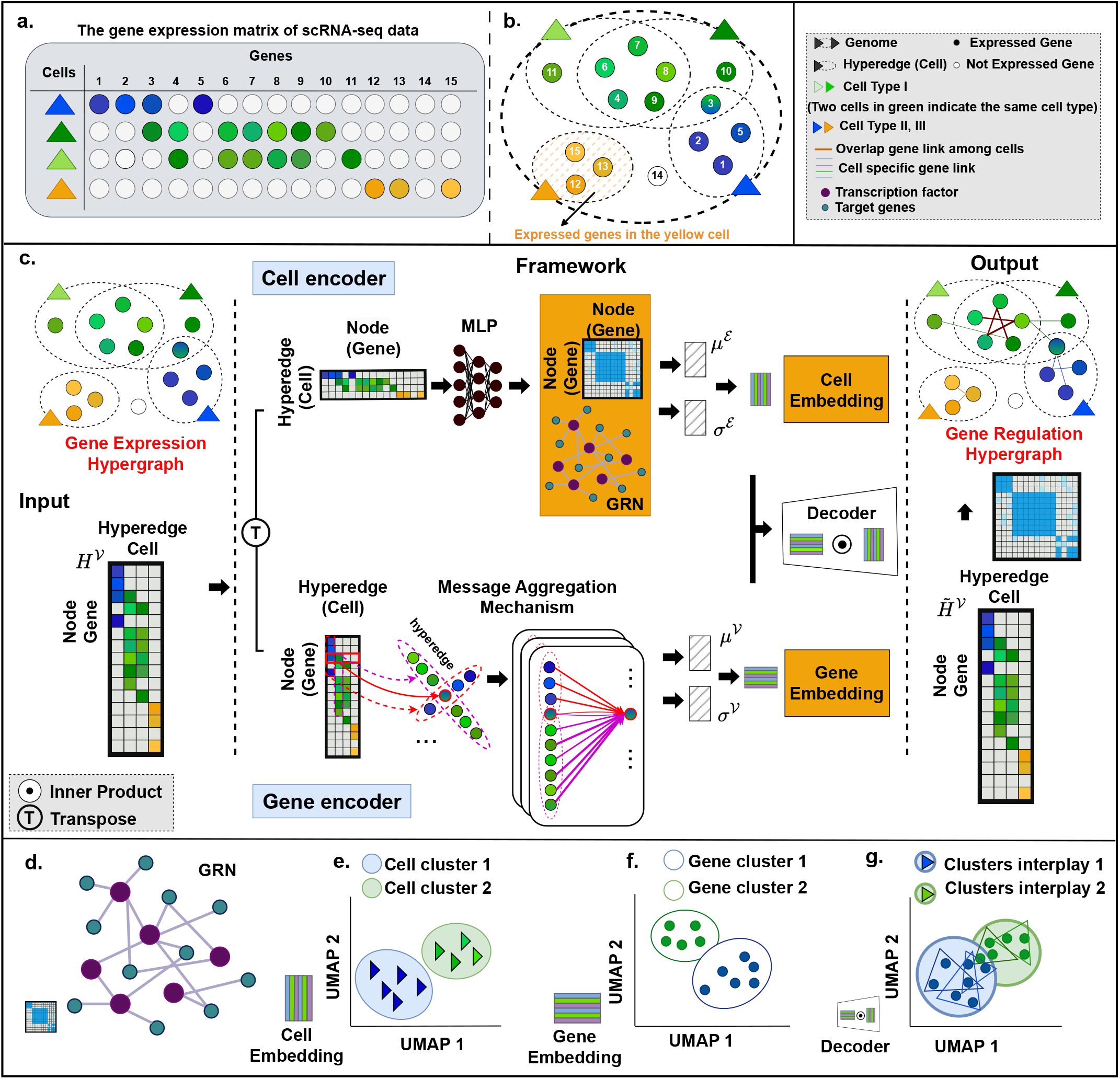
Overview of the HyperG-VAE. **a**, HyperG-VAE, which takes expression value matrix derived from single-cell RNA-seq data as input. In the provided table, four cells exhibit expression across fifteen genes, with color gradients indicating varying gene expression levels (white circles mean no expression). **b**, The colored circles with serial numbers denote distinct genes, expressed within specific cells, functioning as interlinked nodes. These nodes are interconnected by a singular hyperedge (small dashed ellipses) symbolizing the cell (triangle). Together, these nodes and hyperedges form a hypergraph structure. Node coloration reflects a composite of gene expression levels of the given gene across cells; for instance, gene 3 manifests a blend of green and blue hues. The largest dashed ellipse is the genome shared by all cells. **c**, The neural network architecture of HyperG-VAE, two encoders are designed to process the provided input matrix. The cell encoder uses the Structural Equation Model (SEM) to discern cellular heterogeneity and form the GRN, while the gene encoder, employing a hypergraph self-attention mechanism, focuses on gene module analysis. The decoder subsequently reconstructs the input matrix, leveraging the shared latent space of both gene and cell embeddings. Inferred gene regulation hypergraph integrates cellular and gene representations, drawing on relationships derived from the learned GRN. **d-g**, Downstream tasks that can be pursued by HyperG-VAE include GRN construction, clustering both cells and genes, and modeling the interplay between gene modules and cellular heterogeneity. Further details are provided in the legend, located in the upper right of the figure.

HyperG-VAE enhances the GRNs inference by incorporating the above two encoders to mutually augment each other’s embedding quality (Fig.1c) while preserving the high-order gene relations among cells, constrained by hypergraph variational evidence lower bound (Methods). Specifically, the cell encoder incorporates a structure equation model (SEM) on gene coexpression space to infer the GRNs; the learning of gene modules by the gene encoder aids in the inference of GRNs, since the gene module conceivably incorporates TF–target regulation patterns. By integrating the embedding of genes and cells through joint learning, we observe the substantial performance of downstream tasks (Fig.1d-g), including the inference of GRNs, cell clustering, gene clustering, and interplay characterization between gene modules and cellular heterogeneity, among others.

### HyperG-VAE achieves accurate prediction of GRNs

We evaluate the performance on GRNs inference of HyperG-VAE based on the setting of BEELINE framework [27]. Our evaluation encompassed seven scRNA-seq datasets. This includes two cell lines from human and five mouse cell lines (More details can be found in the Supplementary). Furthermore, the EPR and AUPRC are used to evaluate the GRNs performance based on four kinds of ground truth: STRING [28], Non-specific ChIP-seq [29, 30, 31], Cell-type-specific ChIP-seq [32, 33, 34], and loss-/gain-of-function (LOF/GOF) groundtruth network [34]. As recommended by Pratapa et al. [27], our analysis for each dataset prioritized the most variable transcription factors and the top *N* most-varying genes, where *N* is set to 500 and 1, 000. We selected seven state-of-the-art baseline algorithms based on the evaluation of BEELINE to compare with HyperG-VAE, they are DeepSEM [9], GENIE3 [17], PIDC [10], GRNBoost2 [18], SCODE [35], ppcor [7] and SINCERITIES [36]. Introduction and settings of the algorithms can be found in the Supplementary.

Overall, HyperG-VAE demonstrates a discernible enhancement in performance when compared with other baseline methods in terms of both AUPRC and EPR metrics (Fig.2 and Extended Data Fig.1). For scaled results of datasets composed of all significantly varying TFs and the 500 most-varying genes (as shown in Fig.2), HyperG-VAE surpasses the other seven benchmarked methods in 42 of the 44 (95%) evaluated conditions. Compared with the second-best method (DeepSEM), HyperG-VAE enhances results by at least 10% in 19 out of the 44 benchmarks. Furthermore, in comparison to other commendable approaches such as PIDC and GENIE3, our approach registered significant enhancements. For PIDC, 38 out of 44 instances showed improvements of over 10%, with 27 surpassing 30% and 22 going beyond 50%. Similarly, with GENIE3, 33 out of 44 instances marked at least a 10% enhancement, 26 surpassed 30%, and an impressive 20 recorded at least a 50% increase. For scaled results of datasets composed of all significantly varying TFs and the 1000 most-varying genes (Extended Data Fig.1), HyperG-VAE achieves the best prediction performance on 84% (37*/*44) of the benchmarks. In comparison to the runner-up method, DeepSEM, HyperG-VAE outperforms by a margin of at least 10% in 17 of the 44 evaluated benchmarks. Notably, the average enhancement in EPR stands at 11.35%, while that in AUPRC is 7.16%.

**Fig. 2.**
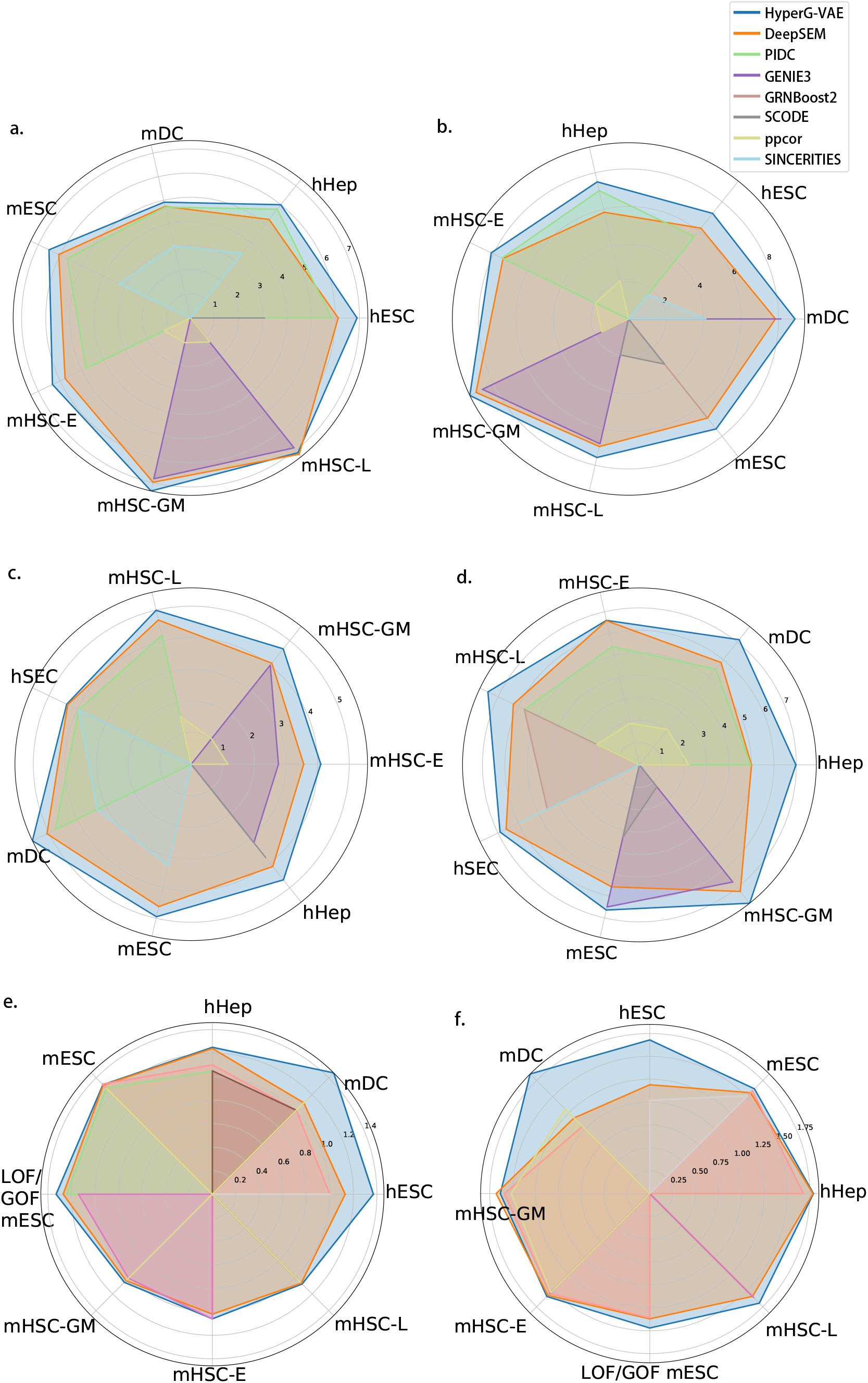
Benchmarks of different GRN inference methods on experimental single-cell RNA-seq datasets by EPR and AUPRC scores. We contrast the scaled performance of HyperG-VAE against seven alternative algorithms. The overall figure shows results for datasets composed of all significantly varying TFs and the 500 most-varying genes. These evaluations span seven datasets, delineated by four unique ground-truth benchmarks: a-b) STRING, c-d) Non-specific ChIP-seq, e-f) Cell-type-specific ChIP-seq, and LOF/GOF. For each figure pair, the left denotes the median AUPRC results, and the right represents the median EPR outcomes. Notably, results inferior to random predictions are omitted from these visualizations. EPR is defined as the odds ratio of the true positives among the top K predicted edges between the model and the random predictions where K denotes the number of edges in ground-truth GRN. AUPRC ratio is defined as the odds ratio of the area under the precision-recall curve (AUPRC) between the model and the random predictions.

With single-cell sequencing data, robustly inferring GRNs from limited cells is pivotal, especially for capturing rare cellular phenotypes and transient states [11, 9]. Here, we explore the fluctuations in EPR performance and the robustness of HyperG-VAE when confronted with limited training data (Extended Data Fig.2a). We constructed mESC datasets [37] composed of all significantly varying TFs and the 500 and 1000 most-varying genes respectively and evaluated the accuracy based on four unique groundtruth benchmark by randomly subsampling single cells following the BEELINE benchmark [27]. Upon adjusting the number of subsampled single cells to 400, 300, 200, 100, and 50, we registered average performance retentions of 94%, 92%, 91%, 80%, and 53%, respectively. Remarkably, when training with cell counts exceeding 100, a robust 79.17% (19*/*24) retained more than 90% of their performance, and for counts greater than 50, a compelling 87.50% (28*/*32) maintained above 80% efficacy. When utilizing cell-type-specific ChIP-seq as the benchmark, the performance remains notably stable, with an average performance retention of 93%. Furthermore, when assessed against the other three ground truths and the training cell count exceeds 50, there’s only a modest decline in efficacy, averaging 88% performance retention in comparison to the median value derived from all cells. Beyond performance evaluation, we also examined HyperG-VAE’s scalability with expansive datasets (Extended Data Fig.2b).

### HyperG-VAE reveals the gene regulation patterns of B cell development in bone marrow

To evaluate HyperG-VAE’s proficiency in elucidating GRNs and to assess the effectiveness of both cell clustering embedding and gene module embedding components within HyperG-VAE, we deployed HyperG-VAE on scRNA-seq data of B cell development in bone marrow [38] (More details of the data can be found in the Supplementary), as illustrated in Fig.3. The progression of B cell development from hematopoietic stem cells follows a sequential yet adaptable developmental pathway governed by interactions among environmental stimuli, signaling cascades, and transcriptional networks [39]. Throughout this developmental trajectory, transcription factors play a pivotal role in regulating cell cycle, differentiation, and advancement to subsequent developmental stages. These critical checkpoints encompass the initial commitment to lymphocytic progenitors, the specification of pre-B cells, progression through immature stages, entry into the peripheral B-cell pool, B cell maturation, and subsequent differentiation into plasma cells [40]. Each of these regulatory nodes is controlled by complex transcriptional networks, which along with sensing and signalling systems determine the final outcomes.

**Fig. 3.**
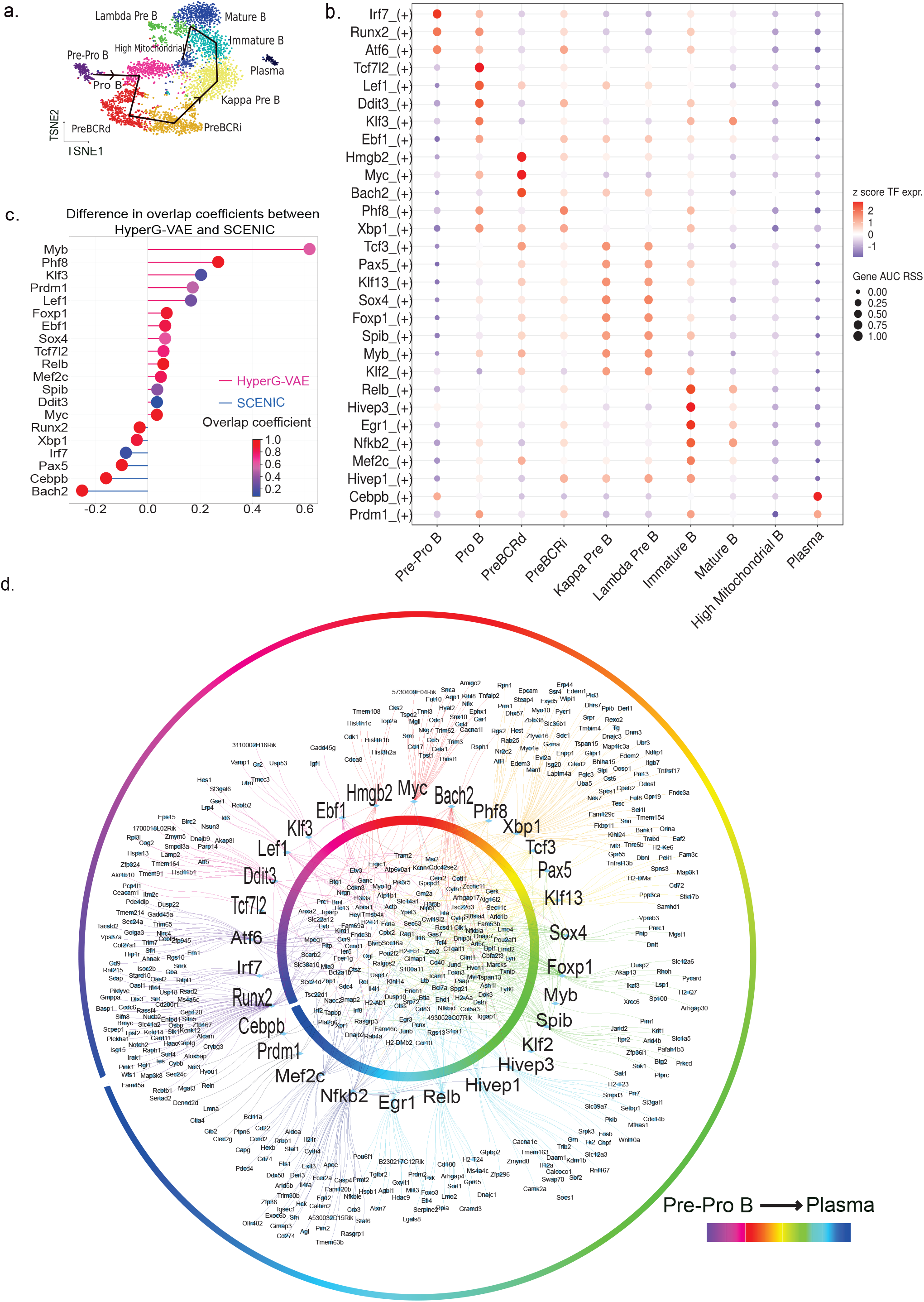
GRN prediction by HyperG-VAE across developmental B cell states in bone marrow. **a**, *t*-SNE visualization of cell embedding on the bone marrow B cell dataset, the embedding is learned by the cell encoder of HyperG-VAE. Black lines depict the trajectory from Pre-Pro B cells to Mature B cells. **b**, The accuracy of GRN prediction by cross-validation with publicly available ChIP-seq datasets. The overlap coefficient quantifies the concordance between sets of target genes for each transcription factor, as derived from GRN prediction and ChIP-seq database respectively. The x-axis represents the difference value of overlap coefficients between HyperG-VAE and SCENIC (default). Pink lines indicate superior performance by HyperG-VAE, while blue lines favor the default SCENIC. The dot-plot illustrates the overlap coefficient of the more effective approach for each regulon, depicted on a color gradient. **c**, Heat map/dot-plot showing TF expression of the regulon on a color scale and cell-type specificity (RSS) of the regulon on a size scale. Cell states are arranged in a sequence that reflects the progression of bone marrow B cell development stages. **d**, The GRN visualization for the bone marrow B cell dataset with ten states from Pre-Pro B state to plasma state, as delineated by HyperG-VAE; inner circle shows the co-binding of shared target genes while outer circle presents TF-focused target genes.

HyperG-VAE uncovers the cell embedding by dimensionality reduction and distinctly segregates the primary cell types across various stages of bone marrow B cell development (Fig.3a). Significantly, HyperG-VAE also effectively captures the linear progression of B cell development, spanning from early pro B, late pro B, large pre B, small pre B, immature B, to mature B cells. In our pursuit to unveil the gene regulation patterns in developmental B cells, our HyperG-VAE, in conjunction with SCENIC [41], successfully identifies established master regulators associated with different developmental stages (Fig.3b, Extended Data Fig.3), including pre-pro B (Runx2), pro B (Ebf1, Lef1), large pre B (Myc, Hmgb2), small pre B (Tcf3, Sox4), immature B (Relb, Egr1), mature B (Nfkb2), and plasma cells (Cebpb, Prdm1).

Furthermore, we conducted a benchmark assessment to compare the performance of HyperG-VAE against SCENIC using its default settings. By the reference of ChIP-seq database [33], the accuracy was evaluated based on the degree of overlap coefficient between the ChIP-seq coverage and the predicted target genes from both methods. Our HyperG-VAE, when combined with SCENIC, demonstrates superior performance compared to the standard SCENIC approach, exhibiting higher accuracy in detecting TF-target patterns for the key transcription factors (as illustrated in Fig.3c). The comprehensive gene regulation network spanning the developmental B cells in the bone marrow is depicted in Fig.3d. We find that the GRNs show TF-target regulation patterns in two ways: transcript factors co-binding to shared predicted enhancers (the inner circle in Fig.3d) and TF-specific target genes (the outer circle in Fig.3d). We also observe that the cooperativity between TFs is stronger within cell types along the development path, indicating that some TFs are involved in multiple stages of B cell development.

### Gene expression module learning enhances HyperG-VAE in GRN inference

Our HyperG-VAE model augments the GRNs prediction by integrating gene space learning, as depicted in Fig.1c. HyperG-VAE uncovers the gene expression modules visualized by Uniform Manifold Approximation and Projection (UMAP) [42] in Fig.4a. By associating these gene modules with the key transcription factors and corresponding target genes of pathways along B cell development, we annotate the gene modules with specific cell types, indicating that these gene clusters are activated in different stages of developmental B cells (Fig.4a,b).

**Fig. 4.**
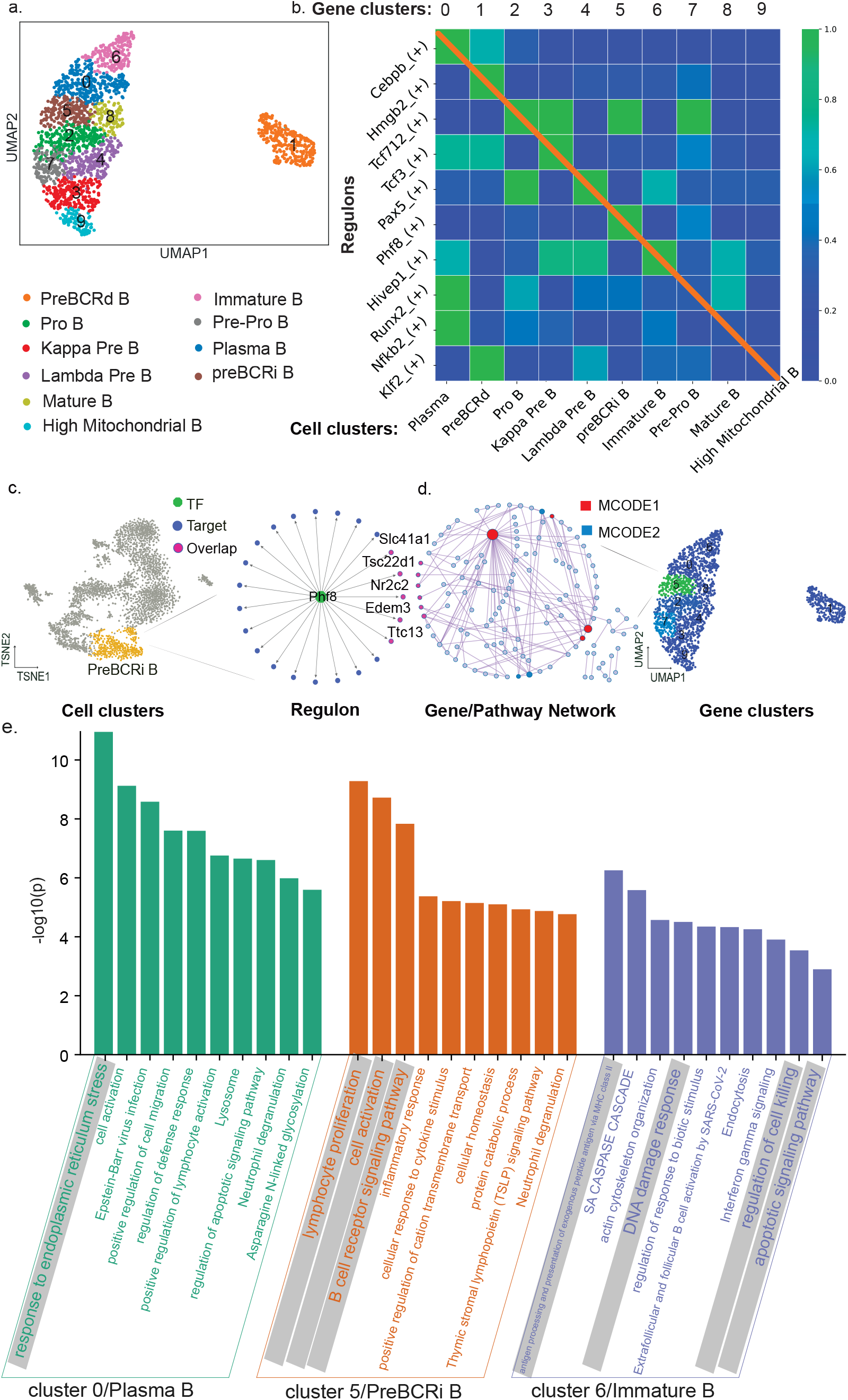
The interplay between gene embedding and cell clusters. **a**, Gene embedding by the gene encoder of HyperG-VAE on developmental B cell data. Gene cluster encoded by numbers are associated with different cell types by colors. **b**, The heatmap illustrates normalized overlap values between gene clusters and TF regulons from different B cell states. Lighter colors indicates larger overlap scores. **c**, *t*-SNE visualization of cellular embeddings with highlighted ‘PreBCRi B’ cell state, together with the associated regulon ‘Phf8 (+)’ and related target genes. **d**, Pathway enrichment analysis on gene cluster 5 with associated Molecular Complex Detection (MCODE) network components. **e**, Pathway enrichment analysis of different gene clusters. The pathways in shade show the dominant gene programs for each gene cluster.

We further apply gene set enrichment analysis (GSEA) [43] (Methods) to investigate the gene clusters (Fig.4e, Supplementary Fig.1-4). The pathways identified through GSEA validate the accuracy of our gene cluster annotations. For example, large pre B cells (cluster preBCRi B) is associated with signals initiating diverse processes which include proliferation and recombination of the light chain gene [44]; the GSEA results show the related pathways: lymphocyte proliferation, cell activation, and B cell receptor signaling pathway. Immature B cells exhibit B cell central tolerance, which is governed by mechanisms such as receptor editing and apoptosis [45]. The pathways identified in the corresponding gene clusters includes antigen processing and presentation of exogenous peptide antigen, DNA damage response, regulation of cell killing and apoptotic signalling pathway. Plasma cells are terminally differentiated B-lymphocytes that secrete immunoglobulins, also known as antibodies [46]. Considering the substantial demands placed on these cells for secretory biological processes, the pathways associated with the relevant gene cluster shed light on the cellular response to endoplasmic reticulum stress.

We show that the gene modules are associated with different biological pathways during B cell development in the bone marrow. These gene modules implicitly incorporate the gene regulation patterns leading to different cell types. On the other hand, distinct cell types of B cell clustering are engaged in various immunological environments [39, 47], resulting in different signalling pathways for B cell activation and fate decisions. We exemplify this joint relationship with an example involving B cells at the large pre-B stage, as shown in Fig.4c,d. This specific cell state (Fig.4c) is characterized by gene regulation patterns associated with cell proliferation, reflected by the regulon Phf8(+) [48]. The corresponding gene cluster (Fig.4d) is linked to a Molecular Complex Detection (MCODE) network which belongs to the lymphocyte proliferation pathway (4e) and shares target genes with the Phf8(+) regulon.

Therefore, our HyperG-VAE reciprocally integrates these two concepts: cell clustering and gene module detection, in the aim of revealing Gene Regulatory Networks (GRNs). Concretely, the cell embedding process groups together similar cells that share common pathways, while the gene modules aggregate genes exhibiting similar regulation patterns, thereby enhancing the accuracy of GRNs computations.

### HyperG-VAE constructs the cell-type-specific GRN on B cell development in bone marrow

We have demonstrated that gene modules associated with various biological pathways correspond to distinct cell types within bone marrow development in B cells. Essentially, these distinct gene regulation patterns influence cell fate commitment, leading to the development of diverse cell types with varying gene regulation profiles. Thus, we employ HyperG-VAE to investigate each individual state of developmental B cells and construct a more accurate GRN for B cell at specific developmental stages, as illustrated in Fig.5. B cell development in the bone marrow can be broadly categorized into four states: pro-B, large pre-B, small pre-B, and mature B [40]. These four stages are visualized using UMAP, as depicted in Fig.5b. For each of these states, we employed HyperG-VAE to compute GRNs and uncover the predominant regulatory patterns, as illustrated in Fig.5a-c. HyperG-VAE effectively reveals the key transcription factors and their associated target genes within each cell state. For example, in the pro-B state, Ebf1 [49] and Pax5 [50] play significant roles, while Myc [38] stands out in the large pre-B state, Bach2 [51, 38] is crucial in the small pre-B state, and Klf2 [52] and Ctcf [53] are notable in the mature state.

**Fig. 5.**
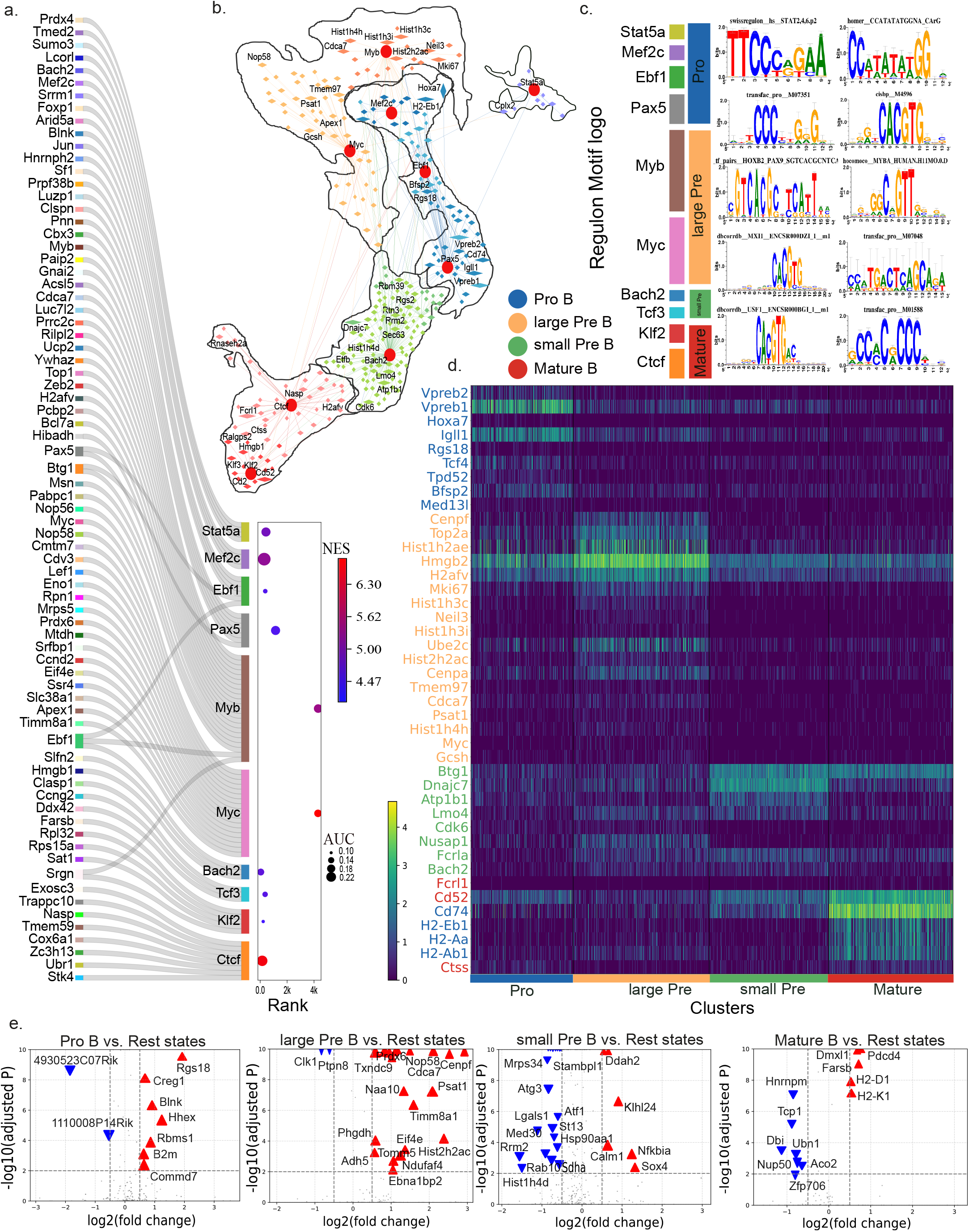
Cell-type-specific GRN analysis of development B cells in bone marrow. **a**, The sankey diagram shows significant regulons and corresponding target genes of different states along B cell development in bone barrow, with normalized enrichment score (NES) encoded by color shade and area under the curve (AUC) score by dot size. **b**, Gene regulatory hypergraph at the cell clustering level, illustrating the four principal B cell states as four hyperedges. Conserved transcription factors are highlighted with red dots, and target genes are depicted as diamonds, where size reflects the logFC in gene expression of a given state compared to others. Highly expressed genes are labeled in the figure. **c**, The motif of significant TFs along the four principal stages. **d**, Heatmap displays the expression of the top genes, ranked by logFC, across cells classified into four distinct cell states. The genes are selected by the overlap of top logFC genes and predicted target genes. The genes’ color corresponds to the cell states in which the regulation pattern is predicted. **e**. Volcano plot of differentially expressed genes of different states. The blue inverted triangles denote down-regulated genes, and the red triangles denote up-regulated genes.

The aforementioned transcription factors, along with their respective target genes, collectively constitute the regulons that characterize the four major cell states, allowing for the construction of a gene regulatory hypergraph at the cell clustering level (Fig.5a,b). For each major state, we overlap the top-predicted target genes by HyperG-VAE (Fig.5b) with the differentially expressed genes (DGEs, Fig.5e) and identify the principal marker genes (Fig.5d). Specifically, Ebf1 and Pax5 are essential in the pro-B state of bone marrow to maintain an early B cell phenotype characterized by the expression of B cell-specific genes such as Vpreb and Igll1 for surrogate light chain production [49, 50]. In the large pre-B stage, the enriched regulons encompass the transcription factor Myc [38] and other genes related to the cell cycle, such as Mki67, Cenpf, Cenpa, and Hmgb2. Additionally, nucleosome-related genes, such as Hist1h2ae and Hist1h3c, are also enriched in this state due to the high rate of cell proliferation. In the small pre-B stage, both Bach2 and Btg1 restrain cell proliferation [54, 55]. It is noteworthy that the mature state markers H2-Ab1, H2-Eb1, H2-Aa and Cd74 are assigned as target genes in the pro-B, suggesting that these genes may be actively repressed in the early B cell development stage.

### HyperG-VAE addresses cellular heterogeneity and learns the cell representation

Cellular heterogeneity is a hallmark of complex biological systems, manifesting as diverse cell types and states within scRNA-seq datasets [56]. We hypothesize that the latent space inferred by the cell encoder of HyperG-VAE captures this biological variability among cells. Leveraging domain expertise, we can map these clusters to known cell types or states, ensuring that the computational predictions align with manual inspection and annotation. To evaluate the performance, we applied HyperG-VAE to three biologically relevant scRNA-seq datasets, including an Alzheimer’s disease (AD) dataset [57], a colorectal cancer dataset [58], and the widely-used mouse brain dataset, known as the Zeisel dataset [59] (More details can be found in the Supplementary). To benchmark HyperG-VAE, we also compared its low-dimensional embeddings with those of six other algorithms: autoCell [60], DCA [61], scVI [62], DESC [63], SAUCIE [64], scVAE [65]. We followed the Louvain algorithm [66] to cluster all the single cells into an identical number of clusters for each method (Methods). To assess the precision of clustering against established reference labels, we employed four metrics: the adjusted Rand index (ARI), normalized mutual information (NMI), homogeneity (HOM), and completeness (COM). These metrics span a scale from 0, indicating random clustering, to 1, signifying perfect alignment with reference clusters, with superior values indicating enhanced accuracy.

Overall, the performance of HyperG-VAE surpasses that of its counterparts, as evidenced in Fig.6a. Specifically, for the Zeisel dataset, the clusters generated using HyperG-VAE align more closely with the existing cell-type annotations, registering an NMI of 83.1% and an ARI of 83.7%. In comparison, the next best-performing algorithm, autoCell, recorded an NMI of 78.0% and an ARI of 80.6%. Furthermore, we evaluated HyperG-VAE’s latent space to determine its ability to capture the biological diversity among individual cells in the Zeisel dataset, as illustrated in Fig.6b. We visualized the data embedding by UMAP. Compared to other algorithms, the distinct separation observed with HyperG-VAE across most clusters indicates effective clustering, suggesting that HyperG-VAE’s cell encoder adeptly distinguishes between various cell states or types. While algorithms such as autoCell, scVI, and scVAE have achieved results that are comparable, the differentiation between their clusters is not as pronounced as with HyperG-VAE. For the remaining algorithms, the substantial overlap among clusters hinders the classifier from producing optimal results. Specifically, compared to other methodologies founded on conventional single-layer Variational Autoencoders (VAEs), the enhanced visualization capabilities of HyperG-VAE underscore the potential benefits of incorporating gene modules in cell embedding processes.

**Fig. 6.**
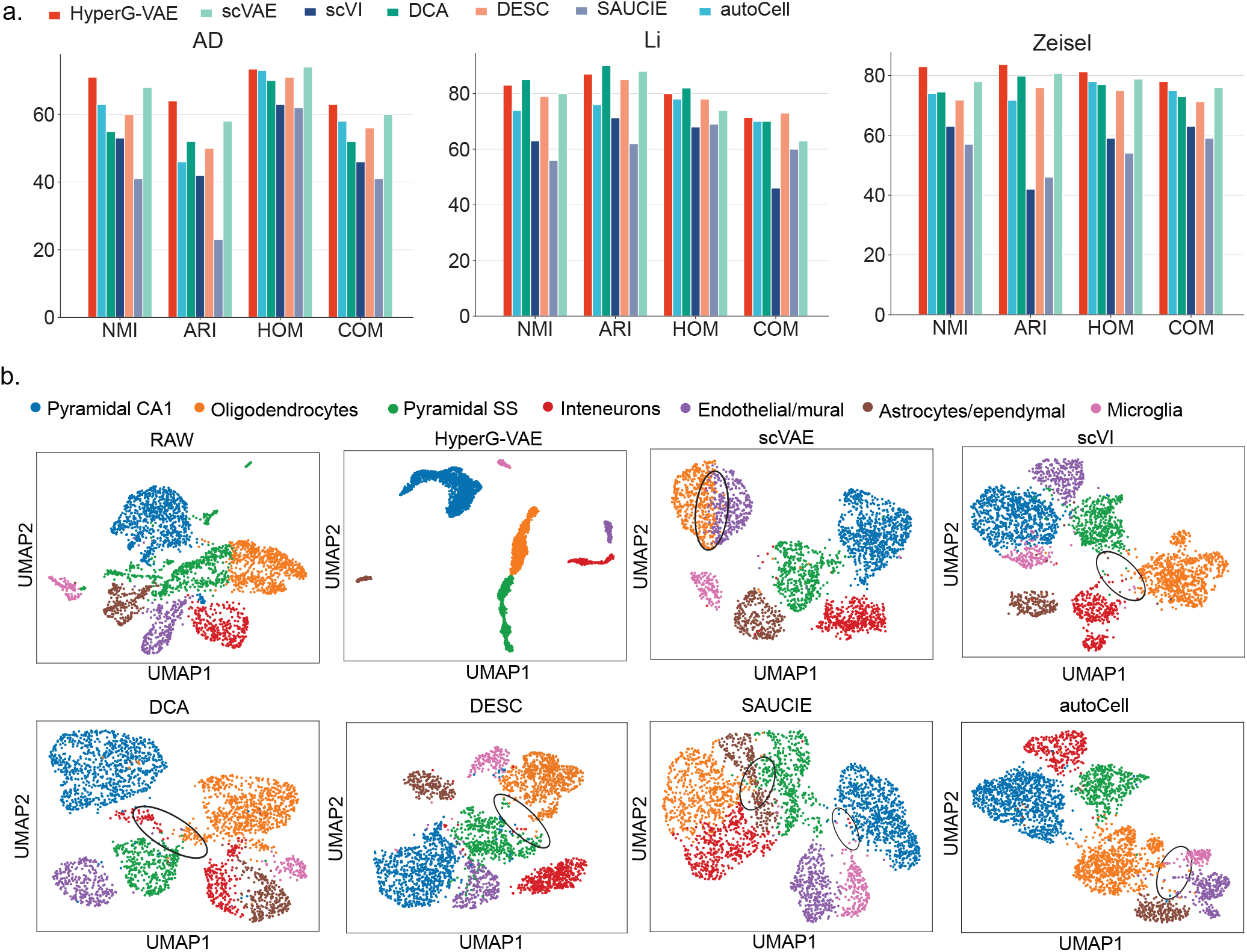
Benchmarks of single cell clustering and embedding. **a**, The cell clustering performance of HyperG-VAE on the single-cell datasets compared with six baseline methods on four key metrics: NMI, ARI, COM, and HOM. **b**, UMAP visualization of latent representations on the Zeisel dataset for different methods. NMI, normalized mutual information (the higher the value the better); ARI, adjusted rand index (the higher the value the better); COM, completeness (the higher the value the better); HOM, homogeneity (the higher the value the better).

## 3 Discussion

In this work, we introduce HyperG-VAE, a sophisticated model designed for the construction of Gene Regulatory Networks (GRNs). Uniquely, HyperG-VAE leverages a hypergraph framework, wherein genes expressed within individual cells are represented as nodes connected by distinct hyperedges, capturing the latent gene correlations among single cells. As a key algorithmic innovation of HyperG-VAE, the transformation of scRNA-seq data into a hypergraph offers unique advantages compared to existing GRNs inference methods. These advantages include improved modeling of cellular heterogeneity, enhanced analysis of gene modules, increased sensitivity to gene correlations among cells, and improved visualization and interpretation of GRNs. This direct use of hypergraph, as opposed to traditional pairwise methods like Star-Expansion (SE) and Clique-Expansion (CE) [67], captures complex multi-dimensional relationships more effectively, avoiding the increased complexity and information loss associated with SE and CE. By maintaining the hypergraph’s original form, HyperG-VAE preserves the data’s full complexity and integrity, enhancing analytical depth and reducing computational demands.

In addition to modelling scRNA-seq data into a hypergraph, HyperG-VAE effectively integrates gene modules and cellular heterogeneity, demonstrating superior performance compared to existing methods. On the one hand, our study reveals that HyperG-VAE outperforms related existing state-of-the-art algorithms in GRNs inference, cell-type classification, and visualization tasks respectively, as evidenced by its enhanced performance across several widely recognized benchmark datasets. On the other hand, we also utilize HyperG-VAE on scRNA-seq data of B cell development in bone marrow [38] to evaluate its performance in a biologically relevant context. Firstly, HyperG-VAE achieves accurate prediction of GRNs and successfully identifies key master regulators and target genes across different developmental stages. Meanwhile, we cross-validated our results with publicly available ChIP-seq datasets [33], further demonstrating HyperG-VAE’s robust performance in predicting regulons based on GRNs inference. Secondly, subsequent evaluations across various tasks further highlighted the effectiveness of HyperG-VAE’s carefully designed encoder components, with their synergistic interaction significantly bolstering the model’s overall performance. Specifically, the cell encoder within HyperG-VAE predicts the GRNs through a structural equation model while also pinpointing unique cell clusters and tracing the developmental lineage of B cells.; the gene encoder uncovers gene modules that implicitly encapsulate patterns of gene regulation, thereby enhancing the accuracy of GRNs predictions. To demonstrate this interaction, we highlight the shared genes between gene clusters and the predicted target genes within cell clusters. These shared genes are notably present in pathways identified by GSEA analysis, signifying the connections between gene modules identified by gene encoders and cell clusters delineated by cell encoders.

Our HyperG-VAE leveraging the self-attention mechanism has undeniably propelled models to achieve remarkable performance [68, 69, 70]. However, despite its prowess, self-attention-based models still have inherent limitations. Specifically, the self-attention’s quadratic complexity concerning sequence length presents challenges. For sequences of length *N*, it necessitates 𝒪(*N* ^2^) computations, rendering it computationally demanding and memory-inefficient, especially for longer sequences. Future efforts to address this limitation will explore to adapt the techniques of attention matrix sparse factorization and positive orthogonal random features, as demonstrated in studies [71, 72], to ease computational demands.

Our proposed model HyperG-VAE holds promise as a foundational framework, adaptable to a multitude of biological contexts in future research endeavors. While our study primarily emphasizes the interrelationships between genes and cells of RNA-seq data using a hypergraph constructed, there is the possibility of evolving into a heterogeneous hypergraph VAE by incorporating other omics data such as scATAC-seq datasets. Such an advancement would facilitate the seamless integration of multi-omics datasets, bolstering tasks such as data integration and GRNs construction. Additionally, while the present model does not explicitly use metadata for genes and cells, future enhancements that integrate this metadata into the hypergraph-centric framework could significantly improve the representations of nodes (genes) and hyperedges (cells). The weights assigned to these hyperedges can also be factored into the model’s learning phase, offering a more comprehensive analysis. In the generative phase of HyperG-VAE, gene-cell interactions proceed through a cohesive mechanism, facilitating the development of a robust GRN underscored by the interplay between gene modules and cell clusters. Moreover, advancing to a single-cell-level, fine-grained gene relation hypergraph application study could further enhance our understanding of single-cell datasets analysis. Furthermore, subsequent research could explore the dynamic construction of temporal GRNs on chronological single cell data, drawing upon the foundational principle of simultaneously considering cellular heterogeneity and gene modules, as demonstrated in this work.

Overall, HyperG-VAE provides a competitive solution for GRNs construction and related downstream works. The consideration of hypergraph helps in effectively capturing the intricate interconnections within complex scenarios. Its inherent versatility allows HyperG-VAE to be adaptable to a wide range of biological contexts, notably including the integration and GRN construction of multi-omics datasets.

## 4 Methods

### Preliminaries

#### Notation

Given a hypergraph ***𝒢*** = { ***𝒱, ℰ***}, where ***𝒱*** = {***v***_**1**_, …, ***v***_***m***_} denotes the set of nodes, and ***ℰ*** = ***e***_**1**_, …, ***e***_***n***_ is the set of hyperedges. Within the hypergraph framework, it is possible for numerous nodes to be interconnected by a solitary hyperedge. Aligning the hypergraph framework with the gene regulation networks (GRNs) paradigm, the expressed genes are mapped as nodes while individual cells stand in as the hyperedges, thus crafting a representation of the cellular architecture as a hypergraph. And, we hope to learn a causal interaction matrix ***Ã*** by HyperG-VAE to approximate the regulatory network ***A*** among genes in real world. Both ***Ã*** and ***A*** are square matrices, where the elements within these matrices signify the levels of regulatory interaction between pairs of genes. In the context of hypergraphs, let ***H***^***𝒱***^∈ *ℝ*^*m×n*^ represent the expression matrix of scRNA-seq dataset, where *m* represents the number of cells and *n* indicates the number of genes. and ***M*** signify the *m* × *n* incidence matrix. The matrix ***M*** is also of size *m* × *n*. If node *i* is linked to hyperedge *j* (gene *i* expressed in cell *j*), then 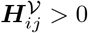 and ***M***_*ij*_ = 1. In the absence of such a link, both 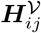 and ***M***_*ij*_ are set to 0. For the hypergraph ***𝒢***, its dual is defined as 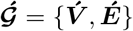. Here, ***𝒱*** = ***ℰ*** and 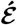 comprises sets ***é***_***i***_ where each ***é***_***i***_ corresponds to edges in *ε* that contain node ***v***_***i***_. As a direct consequence, the feature matrix of the dual, ***H***^***ε***^ ∈ ℛ^*n×m*^, is the transpose of the feature matrix ***H***^***𝒱***^ ∈ ℛ^*m×n*^ of 𝒢.

#### Structural Equation Model

Within the dual of scRNA-seq expression matrix ***H***^***ε***^, we employ the Structural Equation Model (SEM) [73], a statistical approach that integrates factor analysis and multiple regression, to model causal relationships and deduce the intricate dynamics present within gene regulatory networks (GRNs), considering both observed and latent gene interactions. Specifically, our approach is rooted in the Linear SEM:

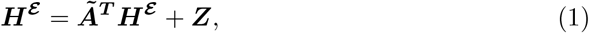

Here, ***Z*** ∈ ℛ^*m×d*^ is the intrinsic noise component following a Gaussian distribution denoted by 𝒩 (**0, *I***). The adjacent matrix ***Ã*** indicates the conditional dependencies among genes. This characteristic implies a mechanism to derive ***H***^***ε***^ from the noise matrix ***Z***, expressed as:

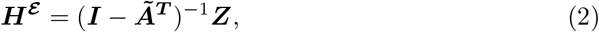

This expression elucidates the relationship between ***H***^***ε***^ and ***Z*** while highlighting the underlying network structure of the GRN as captured by the matrix ***Ã***.

### Hypergraph Variational Evidence Lower Bound

The input scRNA-seq expression matrix ***H***^***𝒱***^ is often noisy and incomplete due to factors like amplification biases during reverse transcription and PCR amplification [19, 74, 20], can compromise the efficacy of basic autoencoders. These autoencoders risk overfitting to training data by solely penalizing reconstruction error, which are influenced by suboptimal expression matrices [75]. To relief the problem, within HyperG-VAE, the hypergraph’s stochastic distribution is tailored to emphasize the latent spaces of nodes and hyperedges, rather than merely relying on observed inputs. Specifically, the node and hyperedge latent spaces are independently derived using distinct encoders and are subsequently refined according to equation (3): related proof could be found in Supplementary A.

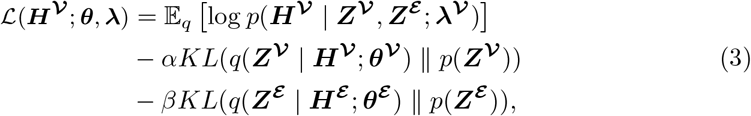

As a crucial loss function of HyperG-VAE, the Evidence Lower Bound (ELBO) is formulated with respect to the observed hypergraph node matrix ***H***^***𝒱***^ and the parameters ***θ*** and ***λ*** which need to be estimated. Specifically, the expectation term, E_*q*_, is the likelihood of the model’s reconstruction of the node matrix using the latent representations for nodes ***Z***^***𝒱***^ and hyperedges ***Z***^***ε***^ . Moreover, the Kullback-Leibler (KL) divergence assesses the deviation of the learned latent distribution, *q*(***Z***^*•*^| ***H***^*•*^ ), from a designated prior *p*(***Z***^*•*^). The coefficients *α* and *β* modulate the magnitude of this regularization.

### HyperG-VAE Node Encoder

For the expression matrix ***H***^***𝒱***^, each row ***h***_***i***_ delineates the expression profile of a gene across diverse cells. Concurrently, a particular gene might manifest across numerous cells and associate with other genes via distinct hyperedges ***e***_***k***_.

In the message-passing phase, row weights should account for expression coherence: genes within the same module typically exhibit consistent expression profiles across cells [76, 77], warranting higher weights than genes with more variable expressions.

Based on the basic idea of GAT [78], we have devised a novel attention computation mechanism tailored for hypergraph, which enables (implicitly) specifying different weights to different nodes share a common hyperedge ***e***_***k***_.

A scoring function *e*: ℝ^*n*^ × ℝ^*n*^ → ℝ computes a score for two genes share a common hyperedge (***h***_***i***_, ***h***_***j***_), which indicates the importance of the expression profiles of two genes ***v***_***i***_ and ***v***_***j***_, which belong to the same hyperedge ***e***_***k***_:

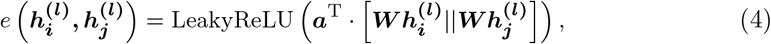

where ***a*** ∈ ℝ^2*n′*^, ***W*** ∈ ℝ^*n′×n*^ are trainable parameters, and || denotes vector concatenation. These attention scores are normalized across all hyperedges using softmax, and the attention function is defined as:

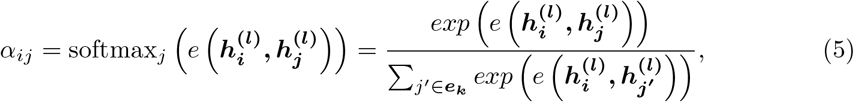

We denote the coefficient matrix, whose entries are *α*_*ij*_, if 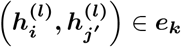, and 0

otherwise. Then, GAT computes a weighted average of the transformed features of the neighbor nodes followed by a non-linearity *σ* as the new representation of ***h***_***i***_, using the normalized attention coefficients:

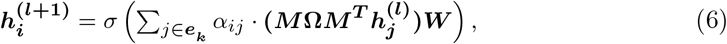

In layer (*l* + 1), the representation of ***h***_***i***_ is denoted by 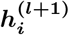. The hyperedge weight matrix **Ω** ∈ ℝ^*n×n*^, is set as the identity matrix, due to the lack of prior knowledge regarding cell relationships. In this paper, we refer to equation (4) to (6) as the computation of each layer in an *L*-layer HyperG-VAE node encoder. We also leveraged the *multi*-*head attention* mechanism, akin to the strategy used in Vaswani et al. [68] to stabilize the learning process of self-attention.

Through the message-passing layers, the input node features of ***H***^***𝒱***^ could be represented as 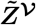, two individual fully connected layers are then employed to estimate the means ***μ***^***𝒱***^ and variances ***σ***^***𝒱***^ of *q*(***Z***^*𝒱*^ | ***H***^*𝒱*^ ; ***θ***^*𝒱*^):

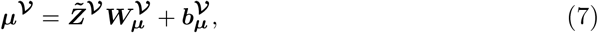

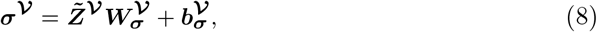

Where 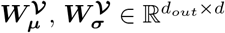, *d* is the dimensionality of the final node embedding ***Z***^***𝒱***^, which is sampled by the following process:

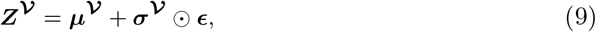

where **ε** ∼ 𝒩 (**0, *I***) and is scaled element-wise by ***σ***^***𝒱***^ . The collective set of parameters, encapsulated within ***θ***^***𝒱***^, offers the posterior estimates for *q*(***Z***^*𝒱*^ | ***H***^*𝒱*^ ; ***θ***^*𝒱*^).

### HyperG-VAE Hyperedge Encoder

Based on the equation (1) and nonlinear version of the SEM proposed by [79], the encoder part of the SEM variational autoencoder could be represented as:

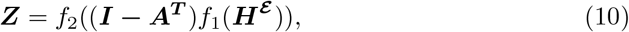

here, the functions *f*_1_ and *f*_2_, parameterized for potential non-linear transformations, adeptly act upon ***H***^***ε***^ and ***Z***, respectively.

Based on equation (10), to encode the high-order semantics and complex relations represented in the form of hyperedges, a hyperedge encoder first conducts a non-linear feature transformation from the observed embedding ***H***^***ε***^ into a common latent space 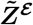, which is as follows:

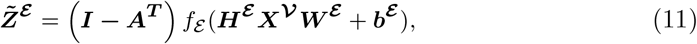

While the gene expression profile is given by ***h***_***i***_, ***X***^***𝒱***^ ∈ ***ℝ***^*m×f*^ denotes the initial *f* - dimensional gene features matrix. Due to the absence of this detailed feature information in our dataset, ***X***^***𝒱***^ is simplified as an identity matrix, ***I***. *f*_*ε*_ stands for multilayer neural network, ***W***^***ε***^ is the learnable weight matrices, and ***b***^***ε***^ is bias.

Given the fused hyperedge embedding 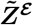, two individual fully connected layers are then employed to estimate the means ***μ***^***ε***^ and variances ***σ***^***ε***^ of *q*(***Z***^*ε*^ | ***H***^*𝒱*^ ; ***θ***^*ε*^ ):

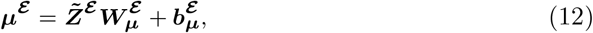

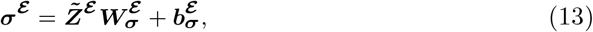

Where 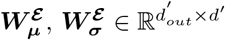, *d*^*′*^ is the dimensionality of the ***Z***^***ε***^, which is sampled by the following process:

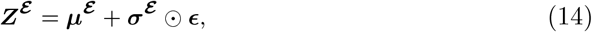

where **ε** ∼ 𝒩 (**0, *I***) and is scaled element-wise by ***σ***^***ε***^ . The collective set of parameters, encapsulated within ***θ***^***ε***^, offers the posterior estimates for *q*(***Z***^*ε*^ | ***H***^*𝒱*^ ; ***θ***^*ε*^ ).

### Generative Model

In the decoding phase, the hypergraph is reconstructed utilizing the latent space representations, ***Z***^***𝒱***^ and ***Z***^***ε***^, acquired from the node and hyperedge encoders, respectively.

To keep the nonlinear SEM of the hyperedge encoder, we first reconstruct the representation of ***H***^***ε***^, and we use the corresponding decoder of equation (10):

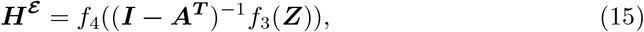

In this work, we can represent the inner content of *f*_4_ as:

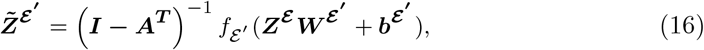

where ***W***^***ε′***^ is the learnable weight matrices, and ***b***^***ε′***^ is bias. Correspondingly, we can get the estimated means ***μ***^***ε′***^ and variances ***σ***^***ε′***^ based on ***Z***^***ε***^ :

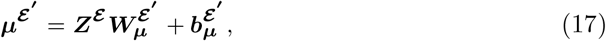

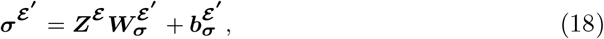

where 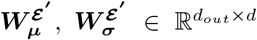, *d* is the dimensionality of the final hyperedge representation ***Z***^***ε′***^, which is sampled by the following process:

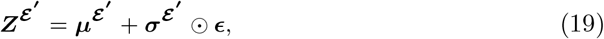

Where **ε** ∼ 𝒩 (**0, *I***) and is scaled element-wise by ***σ***^***ε***^ .

Finally, the estimated hypergraph based on distributions *p*(***H***^*𝒱*^ | ***Z***^*𝒱*^, ***Z***^***ε***^ ; ***λ***^*𝒱*^ ) is represented as:

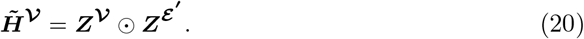

### GRN inference

Central to HyperG-VAE, the gene regulatory network is elucidated by the learned causal interaction matrix ***Ã***, as outlined in equation (11). Crucially, the absolute values within matrix ***Ã*** convey the potential links between genes, underscoring their probability of interrelations. Furthermore, leveraging the state-of-the-art efficacy of HyperG-VAE as reflected across diverse benchmarks, we amalgamate it with SCENIC [41], which is a method renowned for its robustness in GRNs analysis-complements our model by leveraging its ability to distill biologically meaningful gene regulations. This confluence, juxtaposing the precision inherent in HyperG-VAE with SCENIC’s profound insights, furnishes us with a deep learning-based GRNs imbued with biological interpretability.

### Experimental setup

HyperG-VAE was devised to infer gene regulatory networks from scRNA-seq data without relying on cell type annotations. Before feeding into the model, the scRNA-seq expression data underwent log-transformation followed by Z-normalization to ensure optimal data representation. For the initialization of the gene interaction matrix, denoted as ***Ã***, the matrix diagonal was set to zeros, while the other entries followed a Gaussian distribution 𝒩 (**1*/*(*m −* 1), *ϵ***^***ε***^). Here, *m* represents the number of genes, and ***ϵ*** is a small value introduced to prevent entrapment in local optima.

We chose a two-step alternative optimization approach. The RMSprop algorithm [80] was selected initially for tuning the weights within the HyperG-VAE layers over specific epochs. Then, in a separate phase, the weight matrix ***Ã***, which plays a critical role in our architecture, was fine-tuned over another set of epochs, employing a differential learning rate strategy. This bifurcated approach not only fortified the model’s robustness but also ensured granular weight updates in both the matrix and the neural layers. We utilized the kaiming uniform technique [81] to initialize MLPs, crucially defining the initial conditions of our model. The gene (node) encoder, taking the constructed hypergraph as input, employs the Xavier uniform initialization [82] for optimal training. During training, the model’s objective function was guided by a multi-faceted loss: a reconstruction component to maintain data fidelity, two KL divergences (sourced from both the node encoder and the hyperedge encoder) to ensure latent variable alignment with a priori distributions, and a penalty promoting sparsity in the adjacency matrix. This ensured both accuracy in reconstruction and interpretability in inferred gene interactions.

This holistic framework was crafted in Python and leaned heavily on the computational prowess of the PyTorch framework [83], complemented by scanpy [84] for preliminary data handling. Key hyperparameters are selected based on a grid search strategy, more details could be checked in Supplementary Table 1.

#### Datasets and data processing Datasets used for GRN evaluation

We evaluate the performance on GRN inference of HyperG-VAE based on the setting of BEELINE framework [27]. Our evaluation encompassed seven scRNA-seq datasets. This includes two cell lines from human, human embryonic stem cells (hESC) [85] and human mature hepatocytes (hHEP) [86]. Additionally, five mouse cell lines are studied here: mouse dendritic cells (mDC) [87], mouse embryonic stem cells (mESC) [37], mouse hematopoietic stem cells with erythroid-lineage (mHSC-E) [88], mouse hematopoietic stem cells with granulocyte-monocyte-lineage (mHSC-GM) [88] and mouse hematopoietic stem cells with lymphoid-lineage (mHSC-L) [88]. Furthermore, the EPR and AUPRC the GRN performance based on four kinds of groundstruth: STRING [28], Non-specific ChIP-seq [29, 30, 31], Cell-type-specific ChIP-seq [32, 33, 34], and loss-/gain-of-function (LOF/GOF) groundtruth network [34]. Following the guidelines outlined by Pratapa et al. [27], our dataset-specific analysis emphasized the most variable transcription factors and considered the top N most-varying genes, with N being 500 and 1,000. We meticulously adhered to the raw data preprocessing steps detailed in their work and, for evaluation, disregarded any edges that did not originate from TFs.

#### scRNA-seq datasets of Bone marrow developmental B cells

We assess the overarching capability of HyperG-VAE in modeling gene regulatory networks pivotal to B cell development and transformation based on previously published bone marrow developmental B cells datasets [38]. The raw sequencing data in this study were processed using the CellRanger pipeline (version 3.1.0, 10X Genomics), where the “mkfastq” function demultiplexed three Illumina libraries (mRNA transcript expression (RNA), mouse-specific hashtag oligos (HTO), and cell surface marker levels using antibody-derived tags (ADT)) and “count” aligned reads to the mouse genome (mm10) to generate count tables. Analysis was carried out in R using the Seurat package [89], involving filtering of the RNA dataset to include only GEMs expressing more than 300 genes and excluding those with high mitochondrial RNA levels. Normalization was performed using a centered-log ratio method. Doublets were identified in GEMs using both DoubletFinder and HTODemux methods; however, due to discrepancies in classification and challenges with DoubletFinder in identifying similar doublets, subsequent analyses relied solely on HTODemux classifications. GEMs identified as multiplets or negative were removed, leaving a refined dataset of wildtype (WT) singlets, which expressed a median of 1409 genes with 3548 counts. These WT singlets then underwent a transformation process using Seurat’s “SCTransform” function, factoring in the percentage of mitochondrial expression, to prepare a high-quality, normalized dataset for further study.

#### Datasets used for cellular heterogeneity study

We assessed the efficacy of HyperG-VAE by applying it to three pertinent scRNA-seq datasets: an Alzheimer’s disease (AD) study [57], a colorectal cancer investigation [58], and the renowned mouse brain dataset, often referred to as the Zeisel dataset [59]. HyperG-VAE processes raw scRNA-seq gene expression profiles directly. The initial phase of data preprocessing involves rigorous data filtering and quality control. Considering the significant dropout rates characteristic of scRNA-seq expression datasets, only genes with non-zero expression in over 1% of cells and cells with non-zero expression in more than 1% of genes are retained. Subsequently, genes are ranked based on their standard deviation, and the top 2,000 genes in terms of variance are selected for further analysis.

### SCENIC and Chip-Atlas setting

In our approach to further filter reliable gene regulatory networks (GRN) from single-cell RNA-sequencing data, we integrated HyperG-VAE with SCENIC, focusing on discerning crucial gene co-expression modules. Specifically, only the top 0.5% of gene pairs predicted by HyperVAE, based on their co-expression significance, are channeled into SCENIC for rigorous regulon analysis. Using the *MusMusculus* genome reference, our model evaluates regulatory regions defined as 500 bp upstream, 5-kb, and 10-kb centered around each gene’s transcription start site (TSS), collectively referred to as gene-motif rankings. The analysis adopts criteria for GRN derivation of SCENIC: a feature AUC (default: 0.05), gene rank threshold (default: 5,000), and a normalized enrichment score (NES) threshold (default: 3.0).

To validate the predicted regulons, we cross-verified our computational results with publicly available ChIP-seq datasets [33]. Following the foundational settings of SCENIC, we specifically tailored the study to the M. musculus (mm9) genome. Furthermore, in our evaluation approach, we incorporated multiple transcription start sites (TSS) ranges, including 1k, 5k, and 10k, to ensure a comprehensive understanding of gene expression.

### Latent representation visualization and clustering

In both HyperG-VAE and the comparative methodologies, if the size of hidden embeddings exceeded 10, we commenced by extracting the foremost 10 principal components (PCs) via principal component analysis. Subsequently, a cell neighborhood graph was computed, setting the “n neighbors” parameter to 30. Visualization of dataset results was then performed in a two-dimensional space using the default parameters of the UMAP algorithm. For cell clustering, the Louvain algorithm was employed, and the “resolution” parameter was fine-tuned using a binary search to yield a cluster count consistent with cell-type annotations.

### Gene set enrichment analysis (GSEA)

For the analysis of gene clusters, we employed the default settings of Metascape [43]. Specifically, enrichment analysis for given gene lists encompassed pathway and process assessments using GO Biological Processes, GO Cellular Components, GO Molecular Functions, and DisGeNET ontologies. The entire genome served as the background for enrichment. Terms meeting stringent criteria: p-value *<* 0.01, minimum count of 3, and enrichment factor *>* 1.5 (ratio of observed to expected counts)were selected. Statistical rigor was maintained by employing cumulative hypergeometric distribution for p-value calculation, Banjamini-Hochberg procedure for q-value adjustment, and Kappa scores for hierarchical clustering. Clusters, defined by sub-trees with a similarity exceeding 0.3, were identified based on membership similarities. Each cluster is represented by its most statistically significant term. This comprehensive approach ensures robust and reliable insights into gene function and pathway associations.

### Metric used in this paper

#### EPR

EPR is defined as the odds ratio of the true positives among the top K predicted edges between the model and the random predictions where K denotes the number of edges in ground-truth GRN.

#### AUPRC

AUPRC ratio is defined as the odds ratio of the area under the precision-recall curve (AUPRC) between the model and the random predictions.

#### Overlap coefficient

The Overlap coefficient is a similarity measure related to the Jaccard Similarity, but whereas the Jaccard Similarity considers both the intersection and union of two sets, the Overlap Coefficient only considers the intersection relative to the smaller set. It’s used to quantify the overlap between two sets. Given two sets, *A* and *B*, the Overlap Coefficient *O* is defined as:

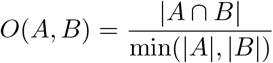

The value of the Overlap Coefficient lies between 0 and 1: A value of 1 indicates that the sets are identical, and 0 indicates that the sets have no elements in common.

#### NES

The Normalized Enrichment Score (NES) quantifies the enrichment of a given motif at the top of a ranking compared to motifs generated by chance. Mathematically, NES is defined as:

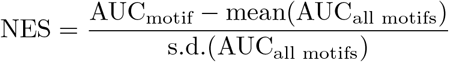

where AUC_motif_ represents the Area Under the Curve for the top 0.5% of the ranked motifs for the gene of interest, and the mean and standard deviation are calculated across the AUCs of all motifs in the dataset. A higher NES indicates a more significant enrichment of the motif in the given context.

#### NMI

NMI (Normalized Mutual Information) quantifies the mutual dependence between two clustering assignments, offering a value between 0 (completely independent assignments) and 1 (identical assignments).

#### ARI

ARI (Adjusted Rand Index) is an adjusted variant of the Rand Index that gauges clustering similarity while accounting for random agreement. Its values range from -1 (perfect disagreement) to 1 (perfect agreement), with 0 indicating random agreement.

#### HOM

HOM (Homogeneity) evaluates whether each cluster comprises solely members of a single class. It ranges from 0 (poor homogeneity) to 1 (perfect homogeneity).

#### COM

COM (Completeness) assesses if all members of a given class are confined to the same cluster, with scores spanning from 0 (low completeness) to 1 (perfect completeness).

## 5 Data availability

We provide all datasets used and analyzed in this study. The gene experimental scRNA-seq datasets were downloaded from Gene Expression Omnibus with the accession numbers GSE81252 (hHEP dataset [86]), GSE75748 (hESC dataset [85]), [GSE98664](http://www.example.com/) (mESC dataset [37]), GSE48968 (mDC dataset [87]), GSE81682 (mHSC dataset ), dataset [85]), [GSE98664](http://www.example.com/) (mESC dataset [37]), GSE48968 (mDC dataset [87]), GSE81682 (mHSC dataset ), GSE168158 (bone marrow developmental B cells dataset [38]), GSE138852(Alzheimer’s disease (AD) dataset [57]), GSE81861( colorectal cancer dataset [58]), and GSE60361 (Zeisel dataset [59]).

The full TF list used in the parts related to SCENIC can be found on the GitHub of pySCENIC https://github.com/aertslab/pySCENIC/tree/master/resources. The ChIP-seq datasets can be accessible through link https://chip-atlas.org/. The motif logo for the regulon used in this paper is available at https://motifcollections.aertslab.org/.

## 6 Code availability

The codes generated in the study are available in GitHub (https://github.com/guangxinsuuu/HyperG-VAE.)

---

**Extended Data Fig. 1.**
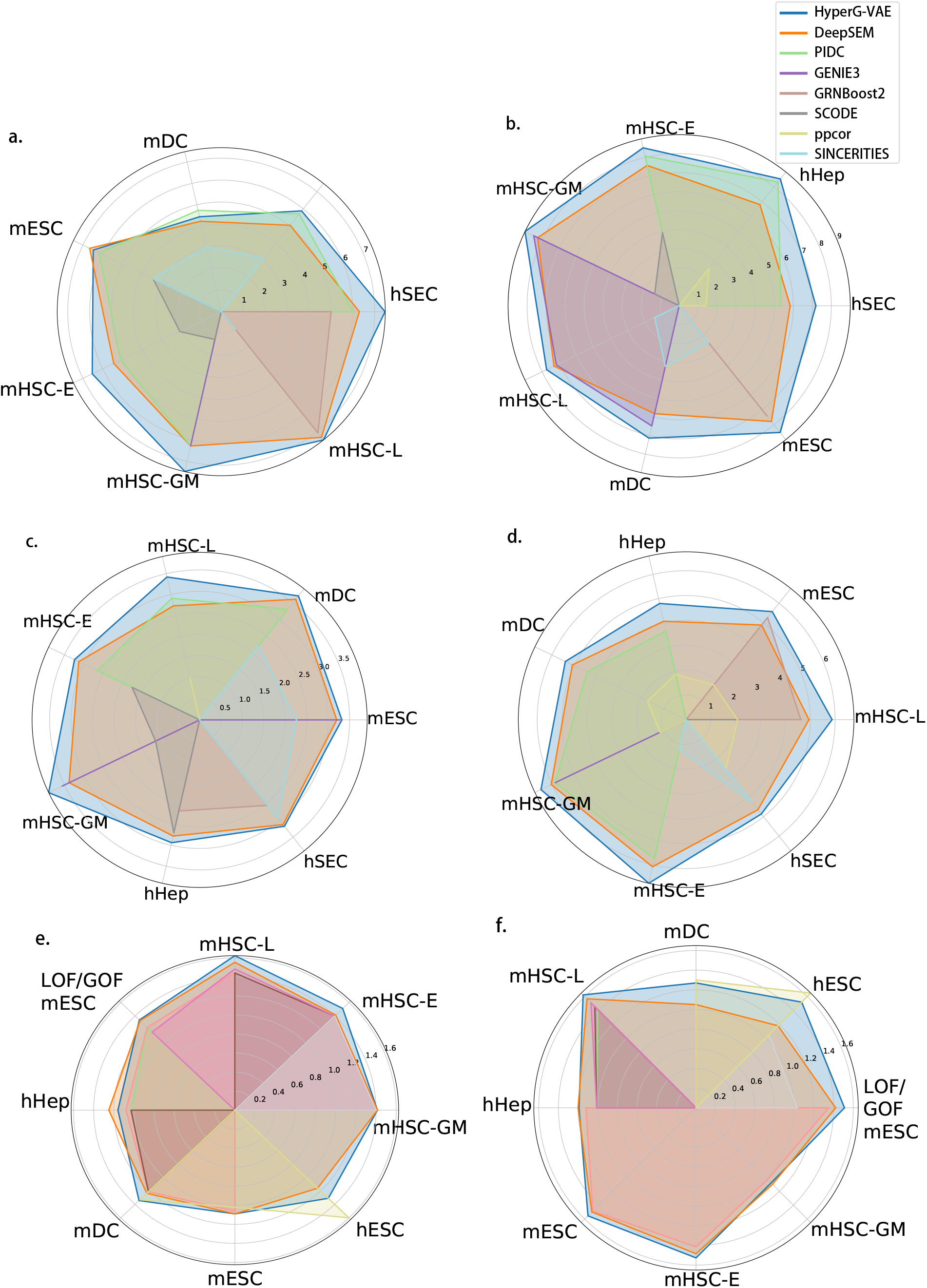
Summary of EPR and AUPRC results for experimental single-cell RNA-seq datasets. The overall figure shows results for datasets composed of all significantly varying TFs and the 1000 most-varying genes. Within each row of illustrations, we contrast the scaled performance of HyperG-VAE against seven alternative algorithms. These evaluations span seven datasets, delineated by four unique ground-truth benchmarks: a-b) STRING, c-d) Non-specific ChIP-seq, e-f) Cell-type-specific ChIP-seq, and LOF/GOF. For every figure pair, the left denotes the median AUPRC results, and the right represents the median EPR outcomes. Notably, results inferior to random predictions are omitted from these visualizations. EPR is defined as the odds ratio of the true positives among the top K predicted edges between the model and the random predictions where K denotes the number of edges in ground-truth GRN. AUPRC ratio is defined as the odds ratio of the area under the precision-recall curve (AUPRC) between the model and the random predictions.

**Extended Data Fig. 2.**
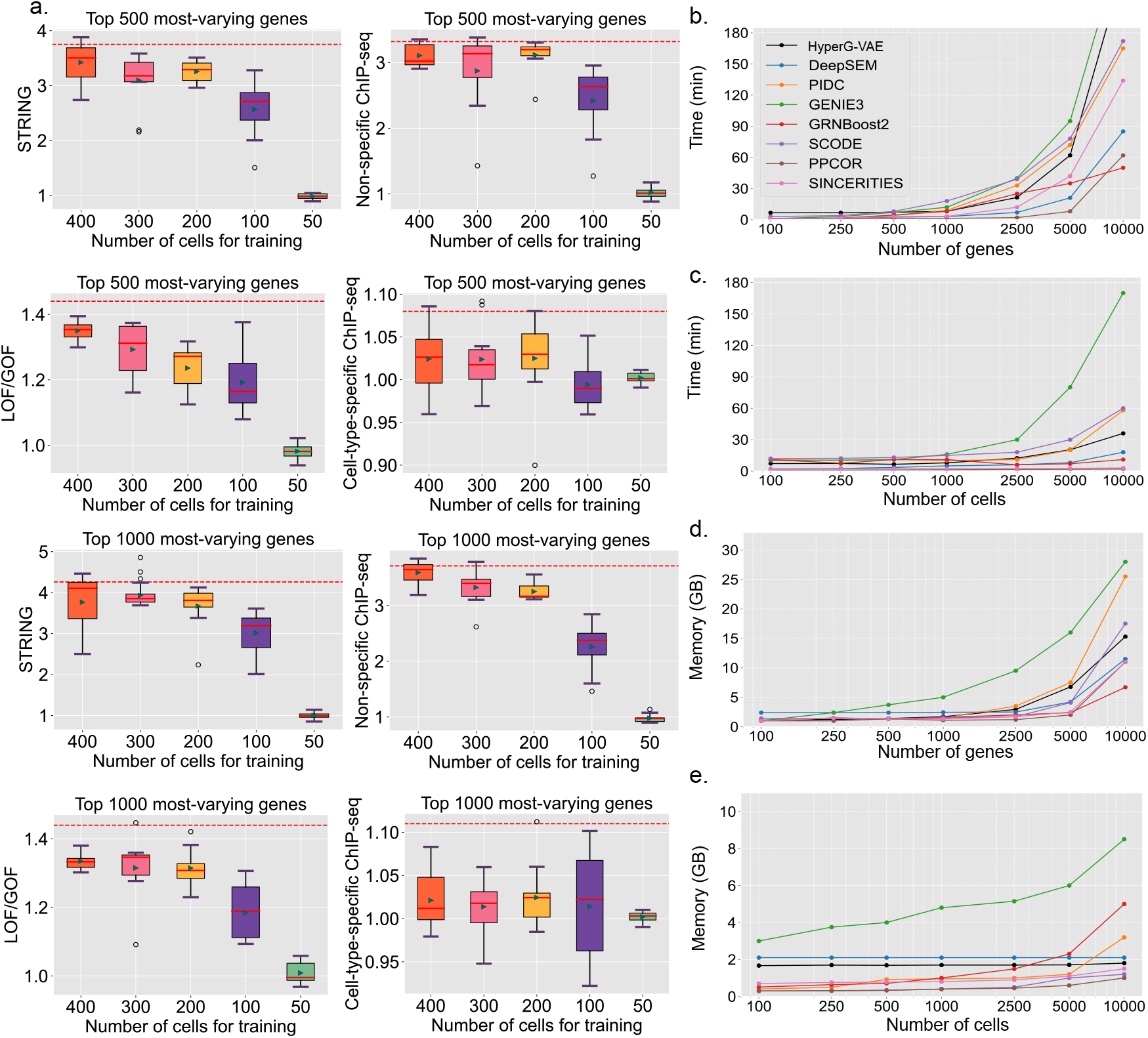
The EPR performance of HyperG-VAE with the limited number of training cells. And, running time and memory cost of different methods on the simulated datasets. **a**, mESC datasets composed of all significantly varying TFs and the 500/1000 most-varying genes are evaluated based on four unique groundtruth benchmarks: STRING, Non-specific ChIP-seq, Cell-type-specific ChIP-seq, and LOF/GOF. The visualization captures the median (represented by the internal line), the interquartile range (shown by the box), and the whiskers (which stretch to 1.5 times the interquartile range). Different colored boxes correspond to distinct training cell numbers, while the green markers within the boxes signify the mean values. Notably, the red dashed line represents the median EPR value across all cell counts. **b**, Running time of training HyperG-VAE and other GRN inference methods on a simulated dataset with 1000 cells when the number of genes for each cell increased. **c**, Running time for training HyperG-VAE and other GRN inference methods on a simulated dataset with 1000 genes for each cell when the number of cells increased. **d**, Memory cost of training HyperG-VAE and other embedding methods on a simulation dataset with 1000 cells when the number of genes for each cell increased. **e**, Memory cost of training HyperG-VAE and other embedding methods on a simulation dataset with 1000 genes for each cell when the number of cells increased.

**Extended Data Fig. 3.**
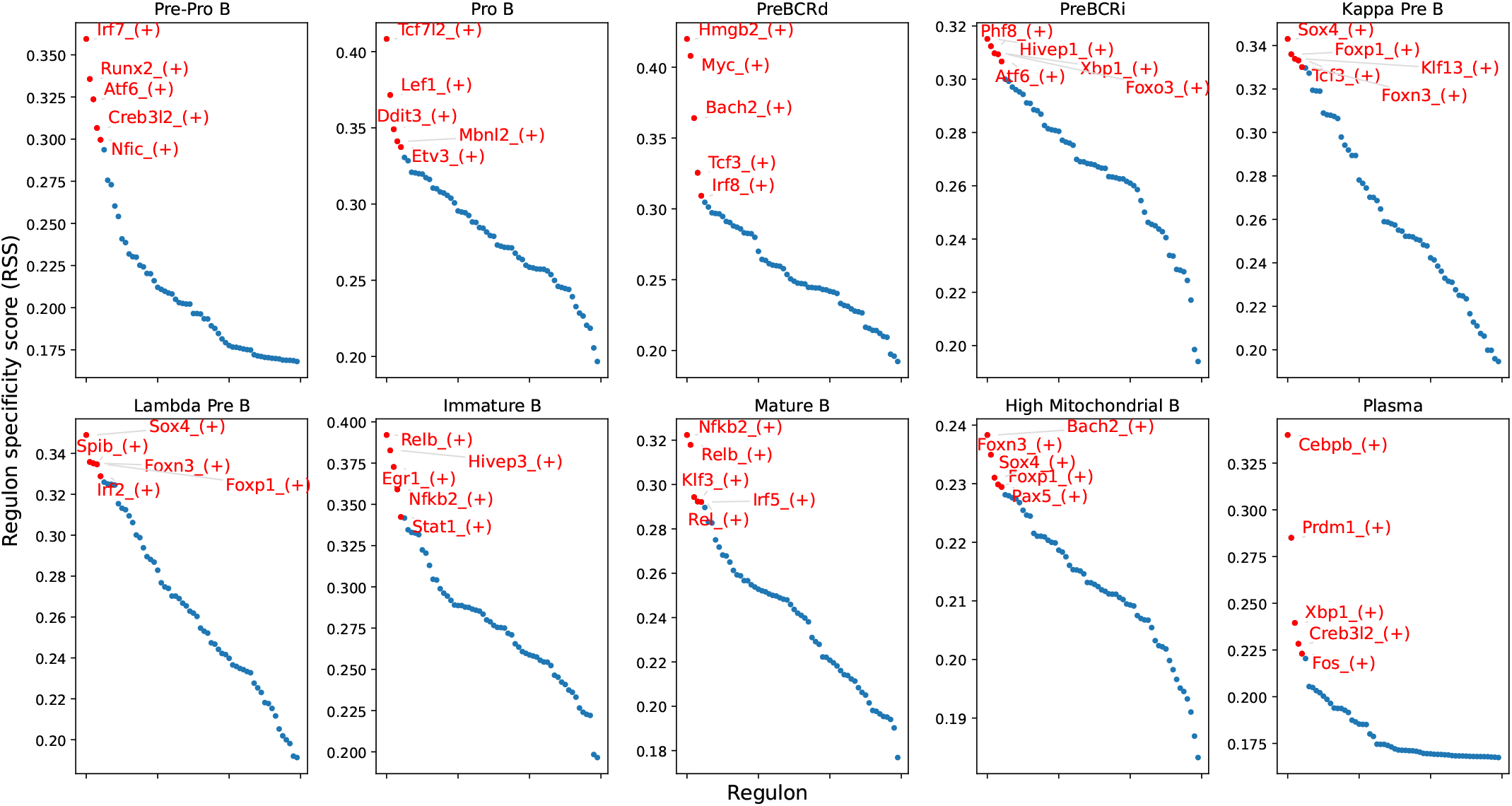
Regulon specificity score for each bone marrow B cell state. The top five regulons in each cell type are highlighted in red and labeled on the plot. The specificity score is shown on the y-axis.

## (Supplementary)

### A Hypergraph Variational Evidence Lower Bound

In the process of HyperG-VAE, latent node embeddings ***Z***^***𝒱***^ and high-order relation embeddings ***Z***^***ε***^ are first generated independently from a parameter-free prior distribution, typically a Gaussian. The observed data points ***H***^***𝒱***^ are then generated conditionally, based on these latent embeddings, with each data point being conditioned on its corresponding latent node embedding ***Z***^***𝒱***^ and high-order relation embeddings ***Z***^***ε***^, parameterized by ***λ***. The objective of HyperG-VAE is to optimize these parameters ***λ*** to maximize the log-likelihood of the observed data. To derive a lower bound for the log-likelihood, known as the Evidence Lower Bound (ELBO). HyperG-VAE leverages Jensen’s Inequality as follows:

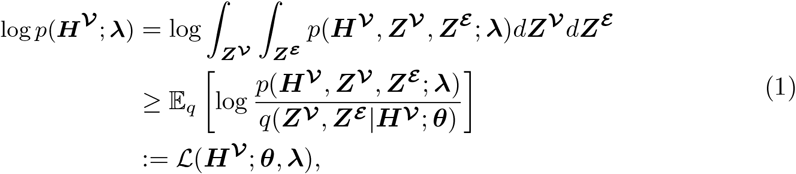

where *q*(***Z***^***𝒱***^, ***Z***^***ε‖***^ ***H***^***𝒱***^ ; ***θ***) is the variational posterior used to approximate the true posterior *p*(***Z***^***𝒱***^, ***Z***^***ε‖***^ ***H***^***𝒱***^ ), and ***θ*** is the parameter that we need to estimate in the learning phase. The Evidence Lower Bound (ELBO) on the marginal likelihood of ***H***^***𝒱***^, denoted as ℒ (***H***^***𝒱***^ ; ***θ, λ***), is derived by applying the logarithmic product rule to the joint probability distribution, facilitating a tractable lower bound for model optimization:

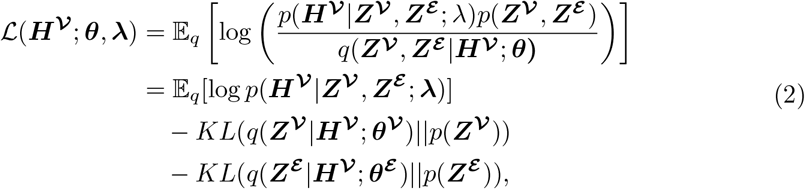

In the variational autoencoder framework, specifically within the HyperG-VAE, the Kullback-Leibler (KL) divergence acts as a regularization factor. It aligns the variational distribution *q*(·|·; ***θ***) with the prior distribution *p*(·), reinforcing the model’s adherence to initial assumptions. Concurrently, the expected log-likelihood of reconstruction, expressed as ℰ[log *p*(·| ***λ***)], dictates the fidelity of data reconstruction from latent embeddings, which are shaped by the learned distribution. The parameter ***λ***, crucial to this reconstruction, is optimized during the learning phase. This dual mechanism ensures that while the model is incentivized to replicate observed data accurately, it remains regularized by the prior, establishing a balance pivotal to the ELBO’s effectiveness in training variational models like HyperG-VAE.

***H***^***𝒱***^ and ***H***^***ε***^ are transposed relations. To better tailor the learning process to specific objectives, weighting components within a loss function, as in Beta-VAE [1], offers nuanced control over regularization, fostering more interpretable and generalizable models. And we will get the ELBO used in HyperG-VAE as:

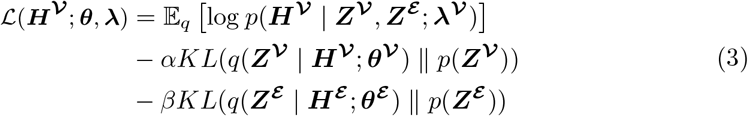

### B Hyperparameter

Hyperparameters tuned in this paper can be found in Supplementary Table 1.

**Supplementary Table 1:** Summary of tuned hyperparameters

| Hyperparamters | Values |
| --- | --- |
| $\alpha$ | 1, 10 |
| $\beta$ | 0.1, 0.2 |
| $\omega$ | 0.1 |
| Learning rate | 0.001, 0.01, 0.0001 |
| Weight decay | 0.1 |
| Batchsize | 64, 128, 256 |
| Dropout rate | 0, 0.5 |
| # of heads | 6, 8 |
| # of training epochs | 70 |

### C Gene expression module learning enhances HyperG-VAE in GRN inference

We further input gene lists of corresponding gene clusters of bone marrow B cell learned by gene cluster within Metascape [2], protein-protein interaction enrichment analysis has been carried out with the following databases: STRING [3], BioGrid [4], OmniPath [5], and InWeb IM [6]. Only physical interactions in STRING (physical score larger than 0.132) and BioGrid are used (details). The resultant network contains the subset of proteins that form physical interactions with at least one other member in the list. If the network contains between 3 and 500 proteins, the Molecular Complex Detection (MCODE) algorithm [7] has been applied to identify densely connected network components. The MCODE networks identified for individual gene lists have been gathered and are shown in Supplementary Figure 1-4.

For each given gene list that input in the Metascape for enrichment analysis, pathway and process enrichment analysis have been carried out with the following ontology sources: KEGG Pathway, GO Biological Processes, Reactome Gene Sets, Canonical Pathways, CORUM, WikiPathways, and PANTHER Pathway. All genes in the genome have been used as the enrichment background. Terms with a p-value *<* 0.01, a minimum count of 3, and an enrichment factor *>* 1.5 (the enrichment factor is the ratio between the observed counts and the counts expected by chance) are collected and grouped into clusters based on their membership similarities. More specifically, p-values are calculated based on the cumulative hypergeometric distribution [8], and q-values are calculated using the Benjamini-Hochberg procedure to account for multiple testings [9]. Kappa scores [10] are used as the similarity metric when performing hierarchical clustering on the enriched terms, and sub-trees with a similarity of *>* 0.3 are considered a cluster. The most statistically significant term within a cluster is chosen to represent the cluster. More details could be found in Supplementary Table 2-5.

**Supplementary Table 2:**
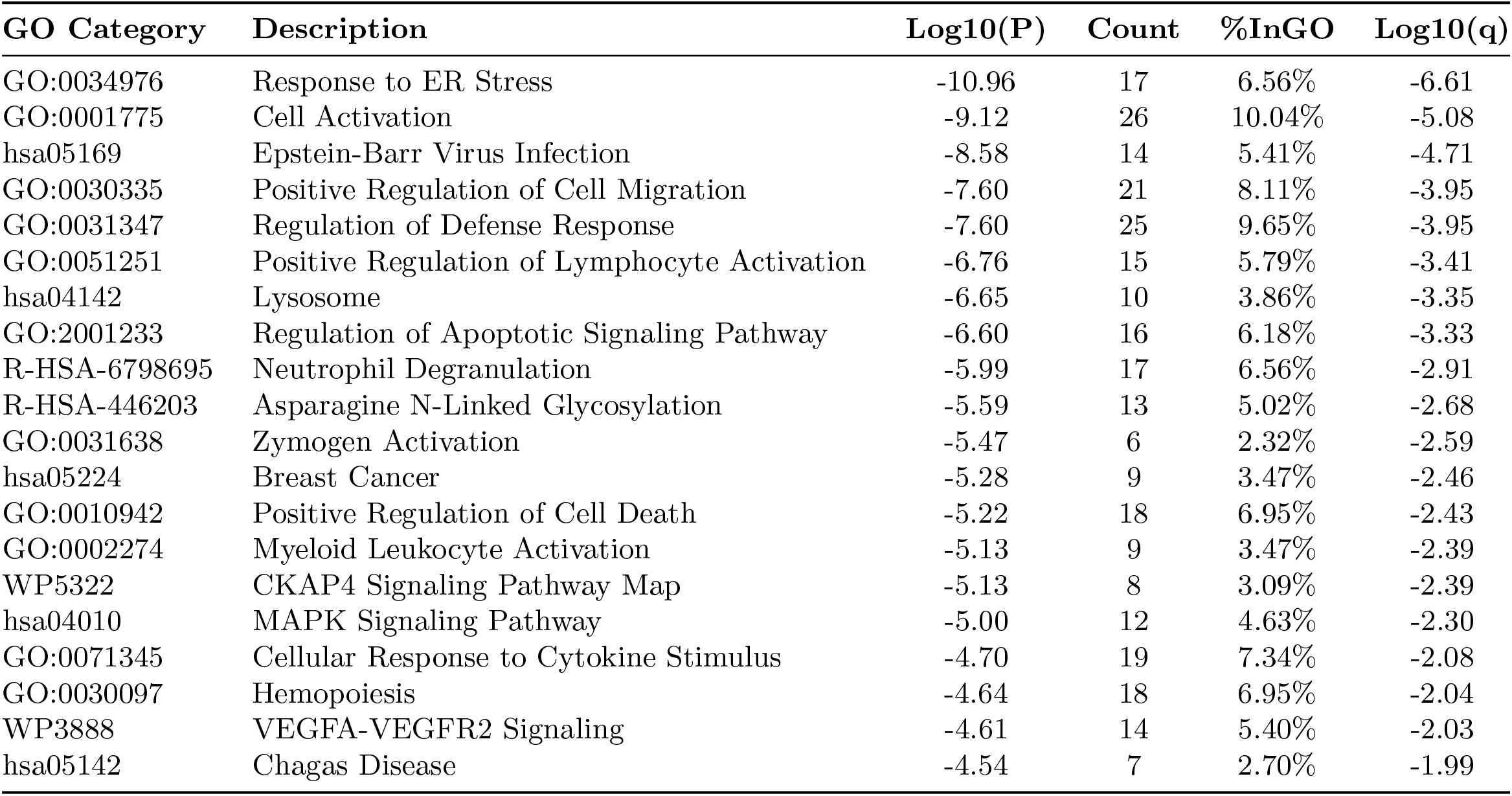
**Pathway and Process Enrichment Analysis of Plasma**. Top 20 clusters with their representative enriched terms (one per cluster). “Count” is the number of genes in the user-provided lists with membership in the given ontology term. “%InGO” is the percentage of all of the user-provided genes that are found in the given ontology term (only input genes with at least one ontology term annotation are included in the calculation). “Log10(P)” is the p-value in log base 10. “Log10(q)” is the multi-test adjusted p-value in log base 10.

| GO Category | Description | Log10(P) | Count | %InGO | Log10(q) |
| --- | --- | --- | --- | --- | --- |
| GO:0034976 | Response to ER Stress | -10.96 | 17 | 6.56% | -6.61 |
| GO:0001775 | Cell Activation | -9.12 | 26 | 10.04% | -5.08 |
| hsa05169 | Epstein-Barr Virus Infection | -8.58 | 14 | 5.41% | -4.71 |
| GO:0030335 | Positive Regulation of Cell Migration | -7.60 | 21 | 8.11% | -3.95 |
| GO:0031347 | Regulation of Defense Response | -7.60 | 25 | 9.65% | -3.95 |
| GO:0051251 | Positive Regulation of Lymphocyte Activation | -6.76 | 15 | 5.79% | -3.41 |
| hsa04142 | Lysosome | -6.65 | 10 | 3.86% | -3.35 |
| GO:2001233 | Regulation of Apoptotic Signaling Pathway | -6.60 | 16 | 6.18% | -3.33 |
| R-HSA-6798695 | Neutrophil Degranulation | -5.99 | 17 | 6.56% | -2.91 |
| R-HSA-446203 | Asparagine N-Linked Glycosylation | -5.59 | 13 | 5.02% | -2.68 |
| GO:0031638 | Zymogen Activation | -5.47 | 6 | 2.32% | -2.59 |
| hsa05224 | Breast Cancer | -5.28 | 9 | 3.47% | -2.46 |
| GO:0010942 | Positive Regulation of Cell Death | -5.22 | 18 | 6.95% | -2.43 |
| GO:0002274 | Myeloid Leukocyte Activation | -5.13 | 9 | 3.47% | -2.39 |
| WP5322 | CKAP4 Signaling Pathway Map | -5.13 | 8 | 3.09% | -2.39 |
| hsa04010 | MAPK Signaling Pathway | -5.00 | 12 | 4.63% | -2.30 |
| GO:0071345 | Cellular Response to Cytokine Stimulus | -4.70 | 19 | 7.34% | -2.08 |
| GO:0030097 | Hemopoiesis | -4.64 | 18 | 6.95% | -2.04 |
| WP3888 | VEGFA-VEGFR2 Signaling | -4.61 | 14 | 5.40% | -2.03 |
| hsa05142 | Chagas Disease | -4.54 | 7 | 2.70% | -1.99 |

**Supplementary Table 3:** Pathway and Process Enrichment Analysis of Kappa Pre B. Top 20 clusters with their representative enriched terms (one per cluster). “Count” is the number of genes in the user-provided lists with membership in the given ontology term. “%InGO” is the percentage of all of the user-provided genes that are found in the given ontology term (only input genes with at least one ontology term annotation are included in the calculation). “Log10(P)” is the p-value in log base 10. “Log10(q)” is the multi-test adjusted p-value in log base 10.

| GO Category | Description | Log10(P) | Count | %InGO | Log10(q) |
| --- | --- | --- | --- | --- | --- |
| GO:0071345 | "Cellular Response to Cytokine Stimulus" | -7.90 | 21 | 10.77% | -3.55 |
| GO:0045321 | "Leukocyte Activation" | -7.17 | 18 | 9.23% | -3.22 |
| R-HSA-6798695 | "Neutrophil Degranulation" | -6.97 | 16 | 8.21% | -3.22 |
| GO:0050778 | "Positive Regulation of Immune Response" | -6.87 | 18 | 9.23% | -3.22 |
| WP5115 | "Network Map of SARS-CoV-2 Signaling Pathway" | -6.71 | 11 | 5.64% | -3.19 |
| GO:0002449 | "Lymphocyte Mediated Immunity" | -6.67 | 10 | 5.13% | -3.19 |
| GO:0002831 | "Regulation of Response to Biotic Stimulus" | -4.83 | 13 | 6.67% | -1.91 |
| GO:0050730 | "Regulation of Peptidyl-Tyrosine Phosphorylation" | -4.40 | 9 | 4.62% | -1.66 |
| R-HSA-1280218 | "Adaptive Immune System" | -4.37 | 16 | 8.21% | -1.63 |
| GO:0007264 | "Small GTPase Mediated Signal Transduction" | -4.34 | 9 | 4.62% | -1.63 |
| GO:0009617 | "Response to Bacterium" | -4.25 | 15 | 7.69% | -1.60 |
| WP4313 | "Ferroptosis" | -4.19 | 5 | 2.56% | -1.57 |
| R-HSA-1280215 | "Cytokine Signaling in Immune System" | -4.06 | 15 | 7.69% | -1.47 |
| M5885 | "NABA MATRISOME ASSOCIATED" | -3.93 | 15 | 7.69% | -1.36 |
| hsa04612 | "Antigen Processing and Presentation" | -3.82 | 5 | 2.56% | -1.28 |
| hsa05200 | "Pathways in Cancer" | -3.73 | 12 | 6.15% | -1.22 |
| GO:0044242 | "Cellular Lipid Catabolic Process" | -3.57 | 7 | 3.59% | -1.08 |
| GO:0045653 | "Negative Regulation of Megakaryocyte Differentiation" | -3.56 | 3 | 1.54% | -1.07 |
| WP23 | "B Cell Receptor Signaling Pathway" | -3.37 | 5 | 2.56% | -0.93 |
| R-HSA-112040 | "G-Protein Mediated Events" | -3.32 | 4 | 2.05% | -0.89 |

**Supplementary Table 4:** Pathway and Process Enrichment Analysis of PreBCRi B. Top 20 clusters with their representative enriched terms (one per cluster). “Count” is the number of genes in the user-provided lists with membership in the given ontology term. “%InGO” is the percentage of all of the user-provided genes that are found in the given ontology term (only input genes with at least one ontology term annotation are included in the calculation). “Log10(P)” is the p-value in log base 10. “Log10(q)” is the multi-test adjusted p-value in log base 10.

| GO Category | Description | Log10(P) | Count | %InGO | Log10(q) |
| --- | --- | --- | --- | --- | --- |
| GO:0046651 | "Lymphocyte Proliferation" | -9.28 | 11 | 5.61% | -5.08 |
| GO:0001775 | "Cell Activation" | -8.72 | 22 | 11.22% | -4.90 |
| WP23 | "B Cell Receptor Signaling Pathway" | -7.83 | 9 | 4.59% | -4.44 |
| GO:0006954 | "Inflammatory Response" | -5.37 | 15 | 7.65% | -2.36 |
| GO:0071345 | "Cellular Response to Cytokine Stimulus" | -5.21 | 17 | 8.67% | -2.25 |
| GO:1904064 | "Positive Regulation of Cation Transmembrane Transport" | -5.15 | 8 | 4.08% | -2.23 |
| GO:0019725 | "Cellular Homeostasis" | -5.10 | 16 | 8.16% | -2.23 |
| GO:0030163 | "Protein Catabolic Process" | -4.93 | 17 | 8.67% | -2.07 |
| WP2203 | "Thymic Stromal Lymphopoietin (TSLP) Signaling Pathway" | -4.87 | 5 | 2.55% | -2.04 |
| R-HSA-6798695 | "Neutrophil Degranulation" | -4.77 | 13 | 6.63% | -1.99 |
| GO:0031347 | "Regulation of Defense Response" | -4.75 | 17 | 8.67% | -1.99 |
| M195 | "PID CMYB PATHWAY" | -4.74 | 6 | 3.06% | -1.98 |
| GO:0033673 | "Negative Regulation of Kinase Activity" | -4.67 | 9 | 4.59% | -1.95 |
| GO:1902532 | "Negative Regulation of Intracellular Signal Transduction" | -4.67 | 14 | 7.14% | -1.95 |
| GO:0042100 | "B Cell Proliferation" | -4.62 | 5 | 2.55% | -1.93 |
| M145 | "PID P53 DOWNSTREAM PATHWAY" | -4.49 | 7 | 3.57% | -1.87 |
| GO:0046631 | "Alpha-Beta T Cell Activation" | -4.23 | 6 | 3.06% | -1.71 |
| GO:0006909 | "Phagocytosis" | -4.09 | 7 | 3.57% | -1.63 |
| GO:0051345 | "Positive Regulation of Hydrolase Activity" | -4.01 | 13 | 6.63% | -1.57 |
| GO:0043462 | "Regulation of ATP-Dependent Activity" | -3.94 | 5 | 2.55% | -1.52 |

**Supplementary Table 5:** Pathway and Process Enrichment Analysis of Immature B. Top 20 clusters with their representative enriched terms (one per cluster). “Count” is the number of genes in the user-provided lists with membership in the given ontology term. “%InGO” is the percentage of all of the user-provided genes that are found in the given ontology term (only input genes with at least one ontology term annotation are included in the calculation). “Log10(P)” is the p-value in log base 10. “Log10(q)” is the multi-test adjusted p-value in log base 10.

| GO Category | Description | LogP | #GeneInGOAndHitList | %InGO | Log(q-value) |
| --- | --- | --- | --- | --- | --- |
| GO:0019886 | "Antigen Processing and Presentation of Exogenous Peptide Antigen via MHC Class II" | -6.25 | 5 | 3.16% | -2.20 |
| M7997 | "SA Caspase Cascade" | -5.58 | 4 | 2.53% | -2.15 |
| GO:0030036 | "Actin Cytoskeleton Organization" | -4.57 | 12 | 7.59% | -1.52 |
| WP707 | "DNA Damage Response" | -4.50 | 5 | 3.16% | -1.49 |
| GO:0002831 | "Regulation of Response to Biotic Stimulus" | -4.35 | 11 | 6.96% | -1.38 |
| WP5218 | "Extrafollicular and Follicular B Cell Activation by SARS-CoV-2" | -4.33 | 5 | 3.16% | -1.38 |
| hsa04144 | "Endocytosis" | -4.26 | 8 | 5.06% | -1.37 |
| R-HSA-877300 | "Interferon Gamma Signaling" | -3.90 | 5 | 3.16% | -1.16 |
| GO:0071900 | "Regulation of Protein Serine/Threonine Kinase Activity" | -3.87 | 9 | 5.70% | -1.15 |
| WP5115 | "Network Map of SARS-CoV-2 Signaling Pathway" | -3.81 | 7 | 4.43% | -1.12 |
| WP3646 | "Hepatitis C and Hepatocellular Carcinoma" | -3.67 | 4 | 2.53% | -1.09 |
| hsa04141 | "Protein Processing in Endoplasmic Reticulum" | -3.54 | 6 | 3.80% | -0.99 |
| GO:0031341 | "Regulation of Cell Killing" | -3.52 | 5 | 3.16% | -0.98 |
| M234 | "PID IL2 STAT5 Pathway" | -3.29 | 3 | 1.90% | -0.82 |
| GO:0043087 | "Regulation of GTPase Activity" | -3.16 | 8 | 5.06% | -0.75 |
| GO:0046649 | "Lymphocyte Activation" | -3.06 | 9 | 5.70% | -0.68 |
| M195 | "PID CMYB Pathway" | -3.00 | 4 | 2.53% | -0.63 |
| R-HSA-6798695 | "Neutrophil Degranulation" | -3.00 | 9 | 5.70% | -0.63 |
| GO:0009617 | "Response to Bacterium" | -2.91 | 11 | 6.96% | -0.56 |
| GO:0097190 | "Apoptotic Signaling Pathway" | -2.90 | 7 | 4.43% | -0.55 |

**Supplementary Figure 1:**
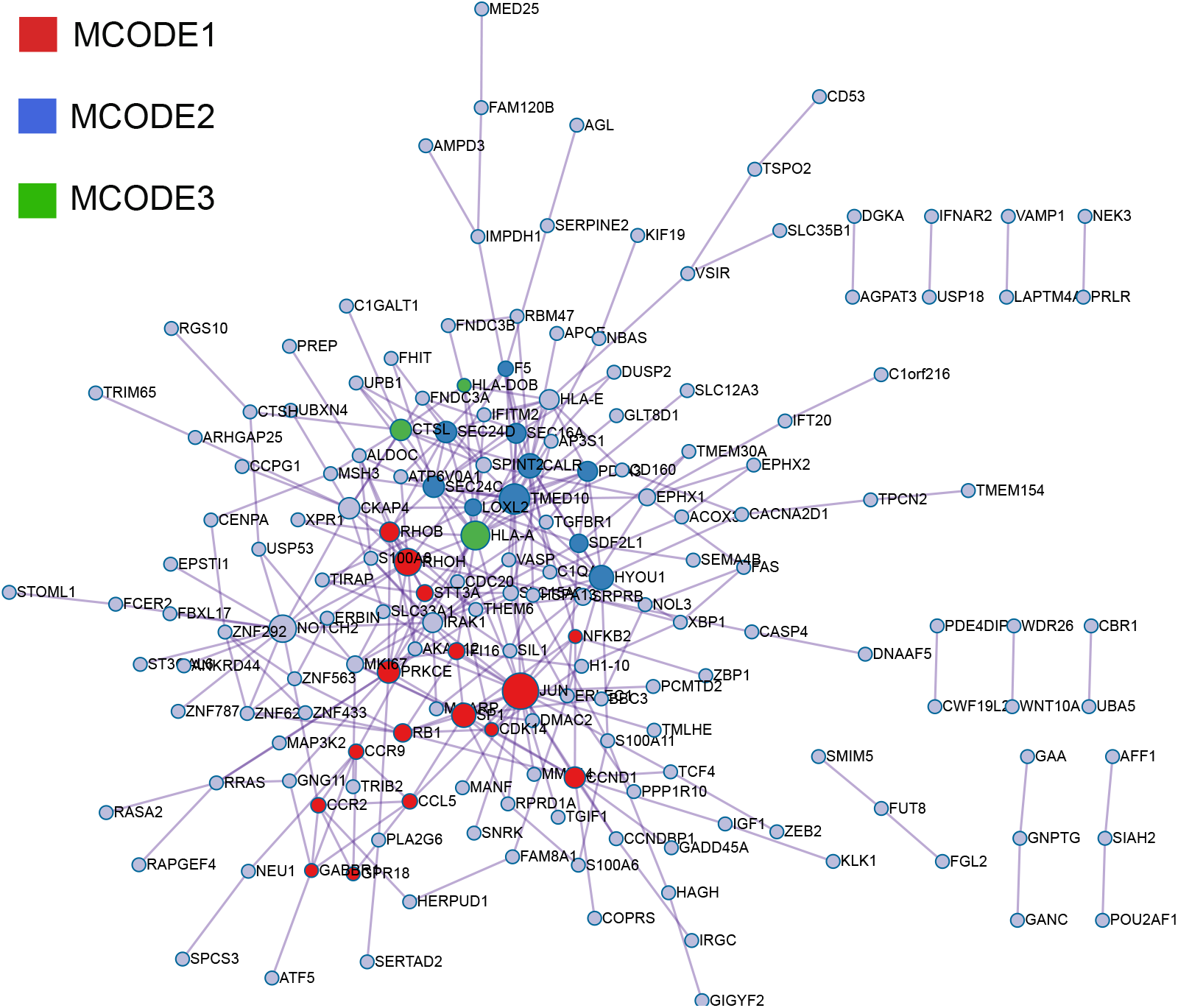
The MCODE network identified for Plasma B gene lists.

### D Details of the single-cell datasets used in the paper

Here, we summarize an overview of the single-cell datasets employed in our analyses. Details about the datasets utilized for gene regulatory network (GRN) benchmark predictions are presented in Supplementary Table 6-7. Furthermore, the datasets encompassing B cell data from bone marrow, as well as those applied in cell clustering and visualization, are delineated in Supplementary Table 8.

### E Overview and Implementation Details of GRN Inference Algorithms in the Current Study

In this research, we provide a concise overview and operational specifics of the GRN inference methodologies employed. The algorithms include DEEPSEM [29], PIDC [30], GENIE3 [31], GRNBoost2 [32], PPCOR [33], SCODE [34], and SINCERITIES [35].

#### DEEPSEM

DeepSEM is a deep generative model designed to simultaneously infer Gene Regulatory Networks (GRNs) and interpret single-cell RNA sequencing (scRNA-seq) data meaningfully. Utilizing a neural network adaptation of the structural equation model (SEM), DeepSEM explicitly captures the regulatory interactions between genes. In benchmarking, DeepSEM outperforms or matches leading methods in GRN inference, scRNA-seq data visualization, clustering, and simulation.

#### PIDC

Using multivariate information theory, the study introduces PIDC, an efficient algorithm that identifies gene regulatory relationships in single-cell gene expression datasets. By leveraging partial information decomposition (PID), PIDC captures higher-order information, making it superior to algorithms based solely on pairwise mutual information. The algorithm’s performance is demonstrated using simulated and experimental data. PIDC also provides insights into network inference variables and factors.

**Supplementary Figure 2:**
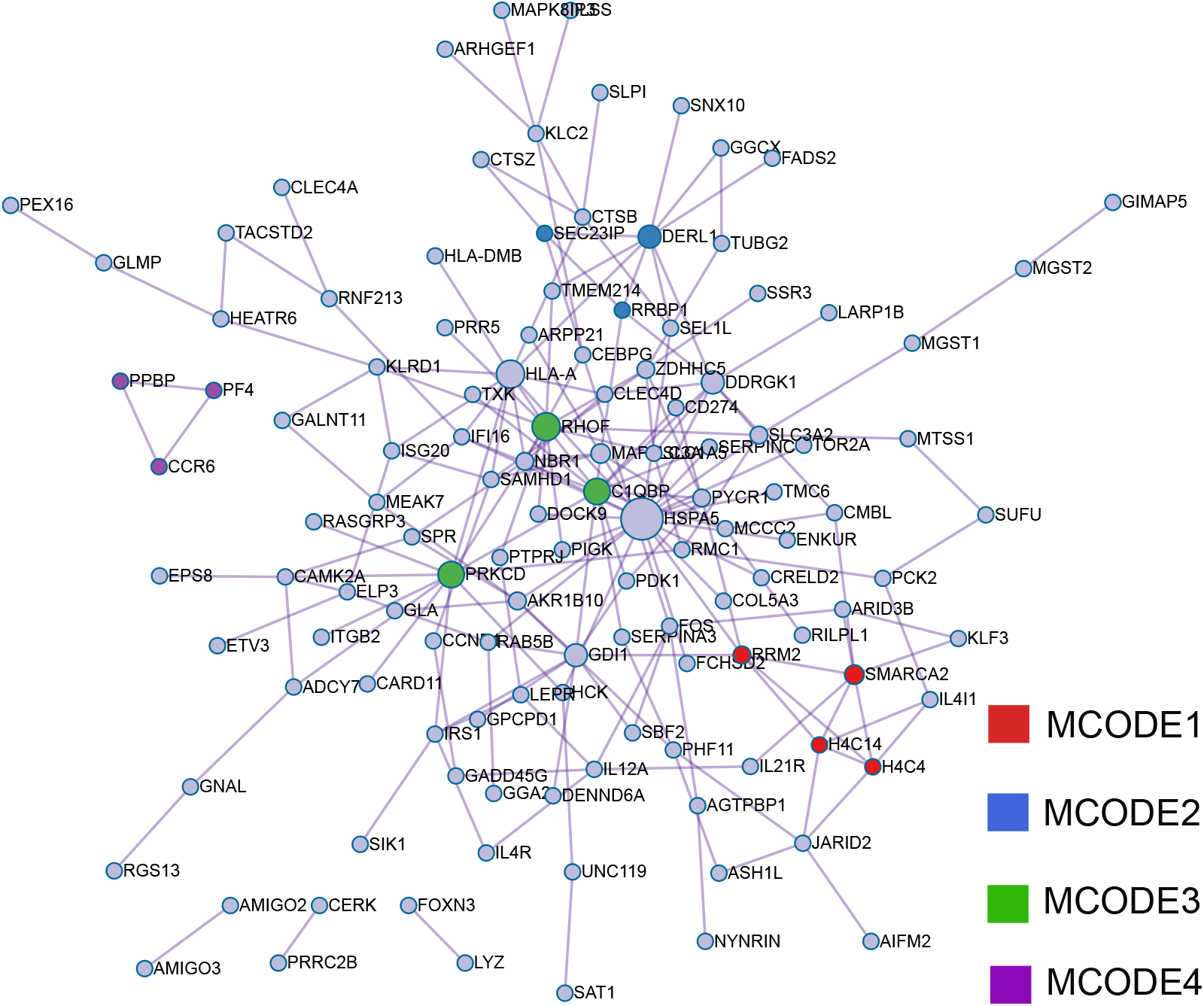
The MCODE network identified for Kappapre B gene lists.

#### GENIE3

GENIE3 is a top-performing algorithm designed to infer genetic regulatory networks (GRNs) from genomic data. By treating the GRN prediction as individual regression challenges, it predicts gene interactions using tree-based methods. Through this approach, GENIE3 can identify potential regulatory links between genes, creating a comprehensive network. Notably, it efficiently deciphers the GRN of Escherichia coli, manages complex interactions, and delivers directed network outcomes, making it a vital tool in GRN analysis.

#### GRNBoost2

GRNBoost2, built on the GENIE3 architecture, is an algorithm designed to infer Gene Regulatory Networks (GRNs) from large gene expression datasets using gradient boosting. To handle the computational challenges posed by voluminous data from technologies like single-cell RNA-seq, the Arboreto framework is introduced.

#### PPCOR

In the context of gene regulatory network predictions, PPCOR utilizes both partial and semi-partial correlations between genes. Adhering to the BEELINE [36] standards, we derived the gene interaction score from the absolute value of the semi-partial correlation between gene pairs.

#### SCODE

SCODE is an innovative algorithm designed to infer Gene Regulatory Networks (GRN) from single-cell RNA-Seq data during differentiation. Utilizing ordinary differential equations, SCODE effectively reconstructs expression dynamics and has demonstrated superior or competitive performance against existing benchmarks. Notably, compared to alternative methods, SCODE operates with significantly reduced runtimes, making it a promising tool for advanced single-cell GRN analyses.

**Supplementary Figure 3:**
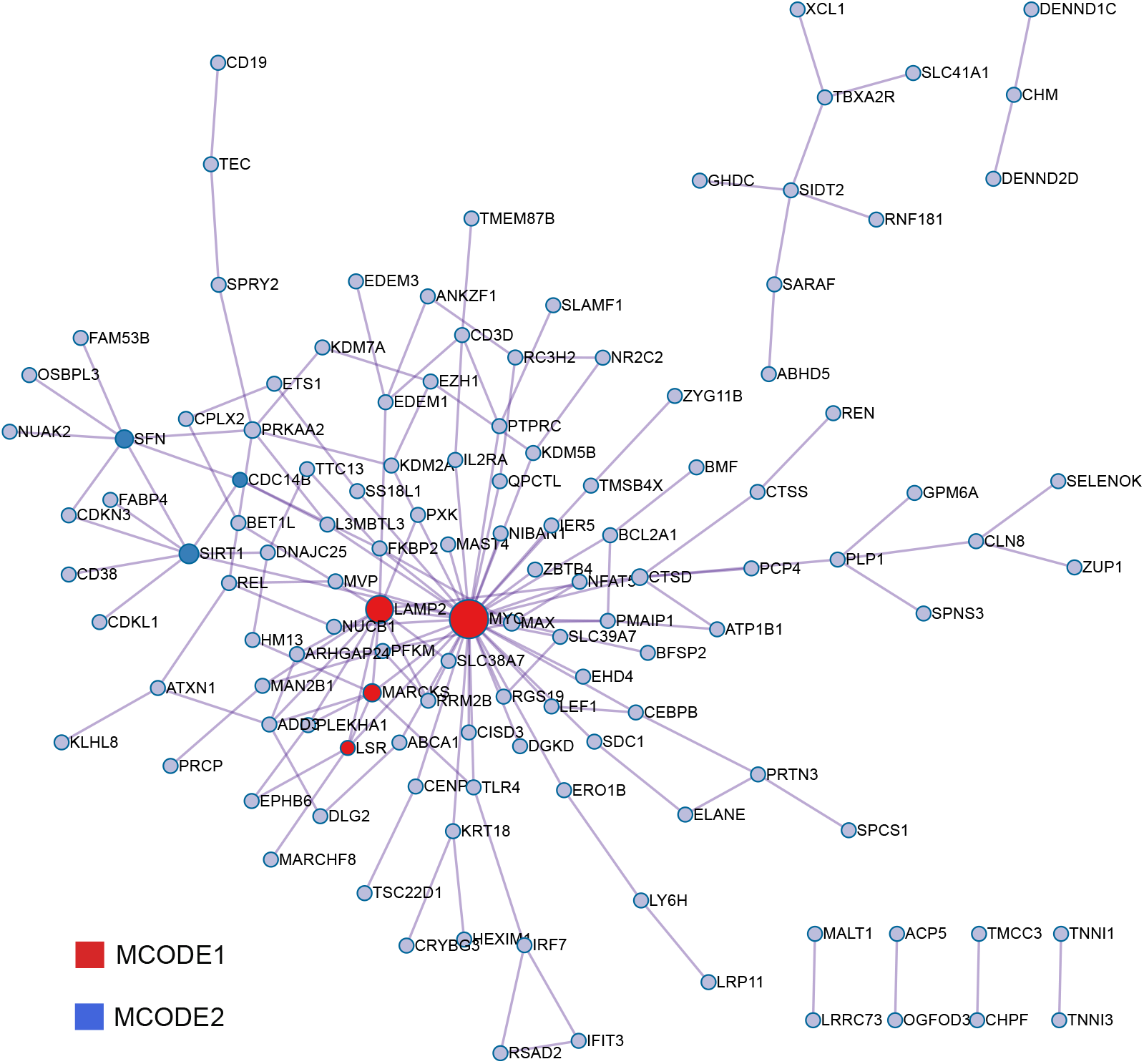
The MCODE network identified for PreBCRi B gene lists.

#### SINCERITIES

SINCERITIES is a novel method designed for the reconstruction of Gene Regulatory Networks (GRN) from single cell transcriptional profiles, specifically focusing on time-stamped cross-sectional data. It utilizes regularized linear regression to determine directed gene-gene interactions based on temporal changes in gene expression distributions. Furthermore, it discerns the nature of gene regulations (activation or repression) using partial correlation analyses. When tested, SINCERITIES outperformed several other GRN inference tools like TSNI, GENIE3, and JUMP3 in accuracy, efficiency, and computational complexity. Its application to real-world data identified BATF as a potential regulator of erythroid development.

### F An Introduction to the Cell Embedding Algorithm and its Implementation in This Investigation

In this research, we provide a concise overview and operational specifics of the cell clustering methodologies employed. The algorithms include autoCell [37], DCA [38], scVI [39], DESC [40], SAUCIE [41], scVAE [42]

#### autoCell

Utilizing a variational autoencoding network, it adeptly handles the sparse nature of scRNA-seq data, facilitating dropout imputation and crucial feature extraction. The tool not only enhances cell trajectory identification but also aids in pinpointing disease-specific gene networks, making it a comprehensive solution for scRNA-seq data analysis.

#### DCA

DCA, or Deep Count Autoencoder, is a denoising method tailored specifically for scRNA-seq datasets, addressing challenges like noise due to amplification and dropout. Employing a negative binomial noise model, DCA effectively captures the intricate gene-gene dependencies and considers the overdispersion and sparsity inherent to the data. Demonstrated to outpace other imputation methods in both quality and speed, DCA is scalable, suited for extensive datasets with millions of cells, and significantly augments biological discovery.

**Supplementary Figure 4:**
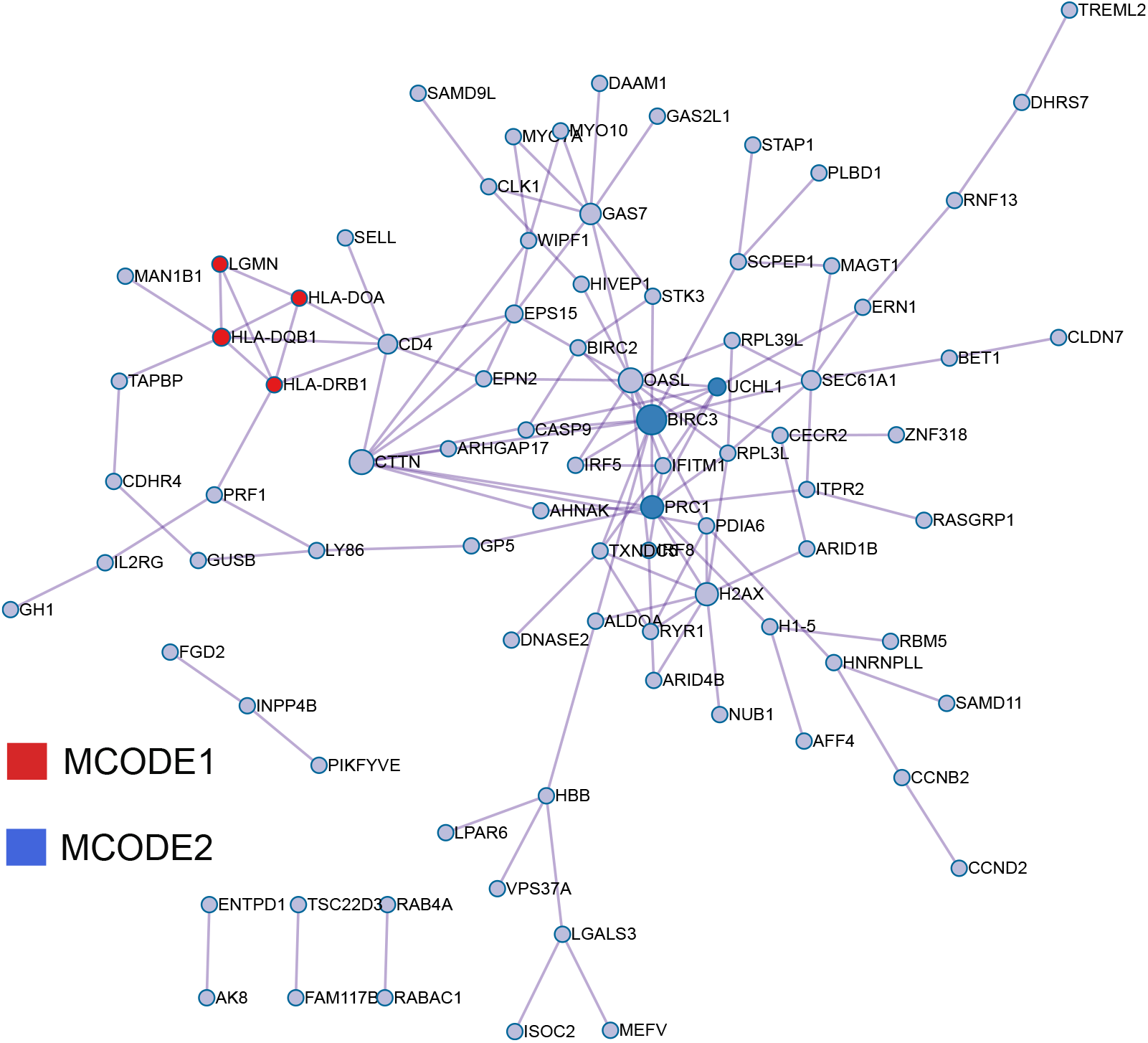
The MCODE network identified for Immature B gene lists.

**Supplementary Table 6:**
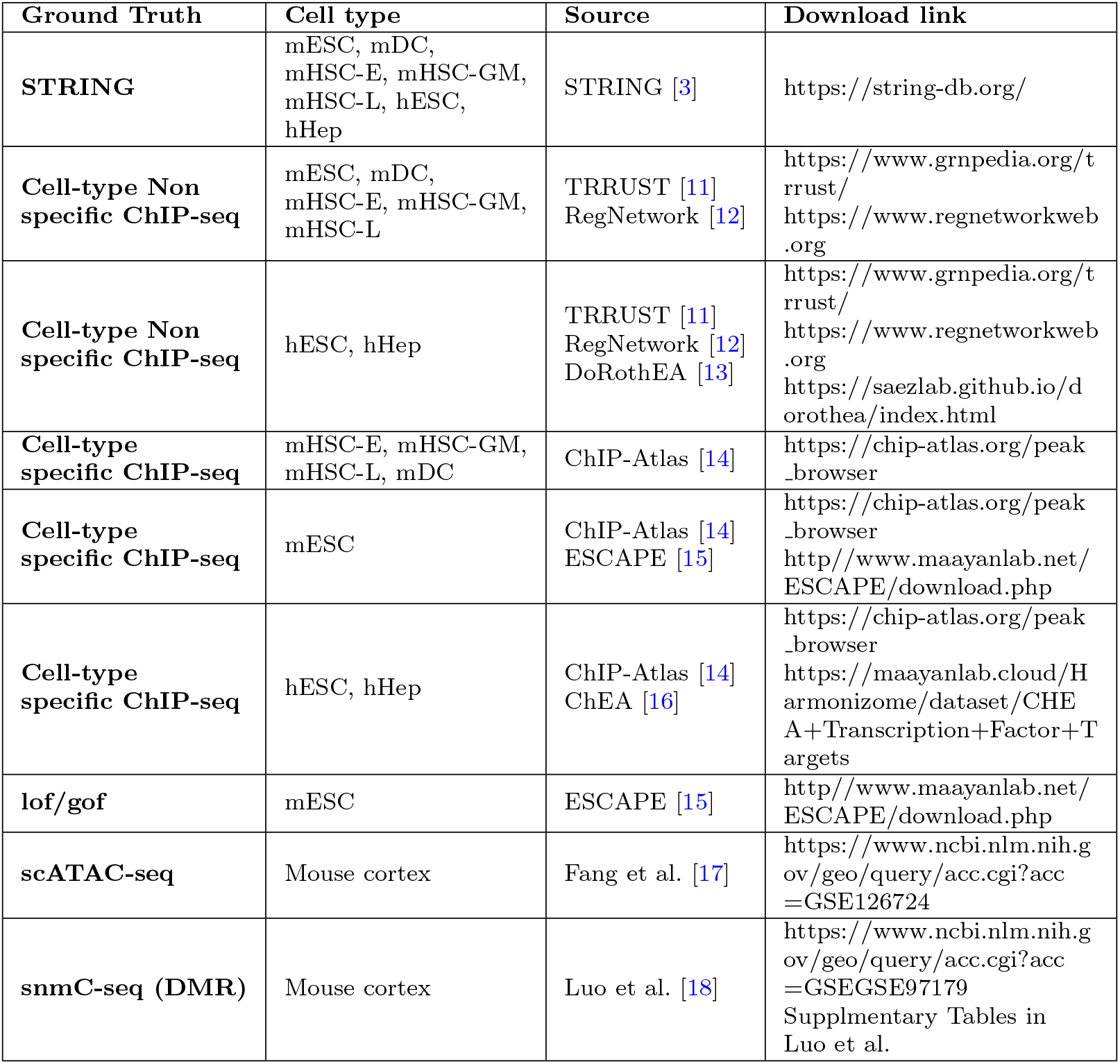
Summary of ground truth GRN networks used in the GRN predictions.

| Ground Truth | Cell type | Source | Download link |
| --- | --- | --- | --- |
| **STRING** | mESC, mDC, mHSC-E, mHSC-GM, mHSC-L, hESC, hHep | STRING [3] | https://string-db.org/ |
| **Cell-type Non specific ChIP-seq** | mESC, mDC, mHSC-E, mHSC-GM, mHSC-L | TRRUST [11]<br>RegNetwork [12] | https://www.grnpedia.org/trrust/<br>https://www.regnetworkweb.org |
| **Cell-type Non specific ChIP-seq** | hESC, hHep | TRRUST [11]<br>RegNetwork [12]<br>DoRothEA [13] | https://www.grnpedia.org/trrust/<br>https://www.regnetworkweb.org<br>https://saezlab.github.io/dorothea/index.html |
| **Cell-type specific ChIP-seq** | mHSC-E, mHSC-GM, mHSC-L, mDC | ChIP-Atlas [14] | https://chip-atlas.org/peak_browser |
| **Cell-type specific ChIP-seq** | mESC | ChIP-Atlas [14]<br>ESCAPE [15] | https://chip-atlas.org/peak_browser<br>http://www.maayanlab.net/ESCAPE/download.php |
| **Cell-type specific ChIP-seq** | hESC, hHep | ChIP-Atlas [14]<br>ChEA [16] | https://chip-atlas.org/peak_browser<br>https://maayanlab.cloud/Harmonizome/dataset/CHEA+Transcription+Factor+Targets |
| **lof/gof** | mESC | ESCAPE [15] | http://www.maayanlab.net/ESCAPE/download.php |
| **scATAC-seq** | Mouse cortex | Fang et al. [17] | https://www.ncbi.nlm.nih.gov/geo/query/acc.cgi?acc=GSE126724 |
| **snmC-seq (DMR)** | Mouse cortex | Luo et al. [18] | https://www.ncbi.nlm.nih.gov/geo/query/acc.cgi?acc=GSEGSE97179<br>Supplmentary Tables in Luo et al. |

**Supplementary Table 7:**
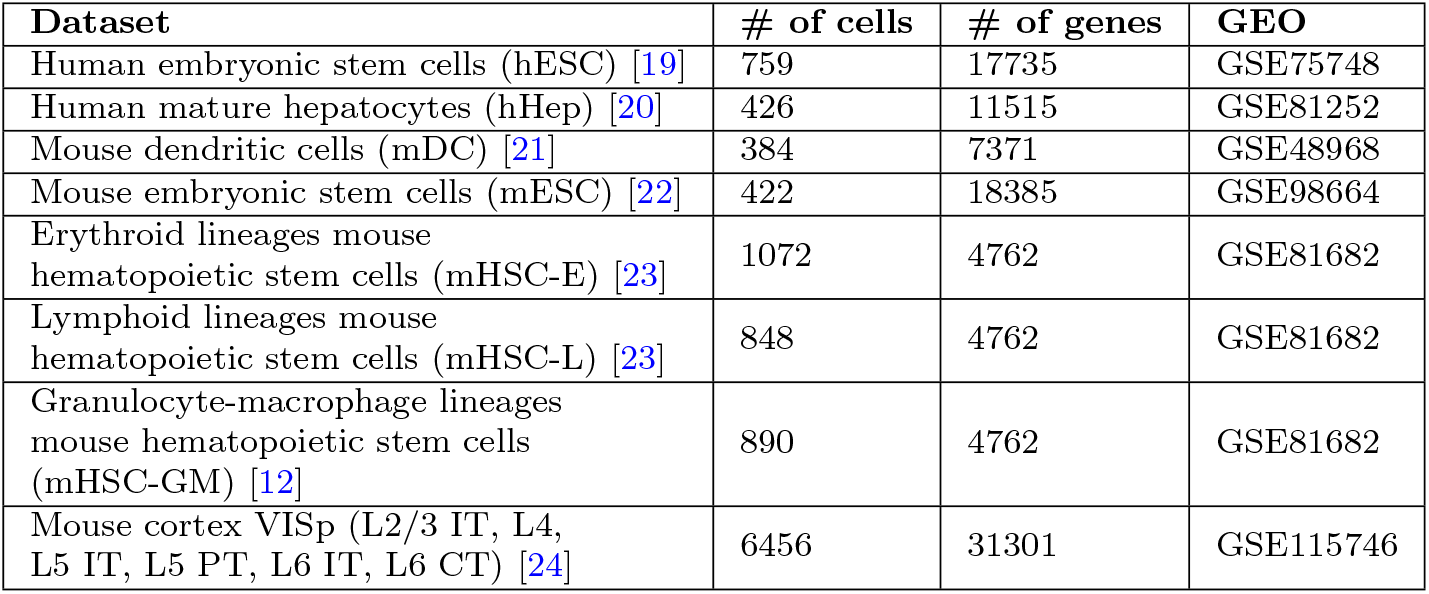
Summary of scRNA-seq datasets used in GRN prediction.

#### scVI

Single-cell variational inference (scVI) is a scalable framework designed to probabilistically represent and analyze gene expression in individual cells, addressing the challenges of technical noise and bias inherent in single-cell transcriptome measurements. Utilizing stochastic optimization and deep neural networks, scVI efficiently aggregates data across similar cells and genes, approximating the fundamental distributions of observed expression values while factoring in batch effects and limited sensitivity. The framework excels in various single-cell analysis tasks such as batch correction, visualization, clustering, and differential expression.

**Supplementary Table 8:**
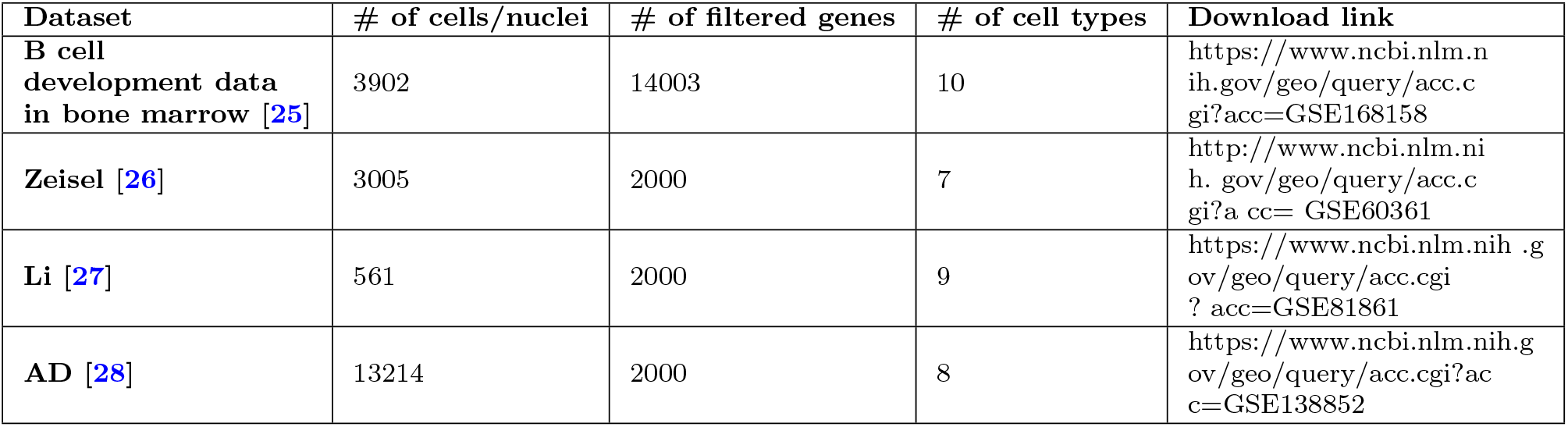
Summary of datasets used in embedding visualization and clustering.

| Dataset | # of cells/nuclei | # of filtered genes | # of cell types | Download link |
| --- | --- | --- | --- | --- |
| **B cell development data in bone marrow [25]** | 3902 | 14003 | 10 | https://www.ncbi.nlm.nih.gov/geo/query/acc.cgi?acc=GSE168158 |
| **Zeisel [26]** | 3005 | 2000 | 7 | http://www.ncbi.nlm.nih.gov/geo/query/acc.cgi?acc=GSE60361 |
| **Li [27]** | 561 | 2000 | 9 | https://www.ncbi.nlm.nih.gov/geo/query/acc.cgi?acc=GSE81861 |
| **AD [28]** | 13214 | 2000 | 8 | https://www.ncbi.nlm.nih.gov/geo/query/acc.cgi?acc=GSE138852 |

#### DESC

DESC is an unsupervised deep embedding algorithm designed to cluster scRNA-seq data, addressing the challenges posed by the increasing number of cells and batch effects. Through iterative self-learning, DESC effectively mitigates batch effects, provided the technical variations across batches are overshadowed by genuine biological differences. With its capability to provide biologically interpretable soft clustering, DESC reveals both discrete and pseudotemporal cellular structures, offering a balanced blend of clustering accuracy, stability, and scalability.

#### SAUCIE

SAUCIE is a deep neural network designed for the analysis of large single-cell datasets, effectively addressing challenges related to batch effects and diverse sample preparations. By utilizing specialized regularizations, SAUCIE ensures interpretability in its learned features, allowing for denoised, batch-corrected data representation, unsupervised clustering, and insightful exploration of complex datasets, such as the immune response of dengue patients.

#### scVAE

Utilizing raw count data, scVAE offers a direct approach to analyzing single-cell RNA sequencing (scRNA-seq), negating the necessity for preprocessing. This method facilitates likelihood-based model comparisons, learns latent cellular representations, and adeptly captures variability across diverse cell populations.

### G Details of predicted regulons in bone marrow B cell states revealed through simultaneous GRN analysis

#### The target genes of transcription factor Irf7 are

Pltp, Tiparp, Gba, Sec24d, Ifi44, Scarb2, Oasl2, Oasl1, Rilpl1, Usp18, Klrd1, Isoc2b, Stard10, Irf7, Bcl2a1b, Rsad2, Mpeg1.

#### The target genes of transcription factor Atf6 are

Itm2c, Xpr1, Atf6, Fcer1g, Sec16a, Cobll1, Fndc3b, Tiparp, Pde4dip, Sec24d, Col27a1, Tmem214, Fam69a, Golga3, Hip1r, Gadd45a, Tacstd2, Cd9, Klrd1, Ceacam1, Rgs10, Ccnd1, Erich1, Vps37a, Ttc13, Anxa2, Ccr9, Sfi1, Trim7, Sec24a, Pik3r5, Ern1, Trim65, Slc38a10, Dusp22, Fyb, Zfp945, Nlrc4, Ahnak, Mpeg1.

#### The target genes of transcription factor Tcf7l2 are

Pcp4l1, Bmf, Akr1b10, Usp47.

#### The target genes of transcription factor Lef1 are

Lrp4, Rag1, Sdc4, Tifa, Lef1, Id3, Il16, Myb, Gse1, Rcbtb2, Cwf19l2, Ebf1, Msi2, Cyth1, Hes1, St3gal6, Tcf4.

#### The target genes of transcription factor Ddit3 are

Tram2, Dnajb2, Ier5, Hsd11b1, Bmf, Cebpb, Lamp2, Tmem164, Abca1, Eps15, Heyl, Zfp324, Tmem91, Atf5, Smpdl3a, Ddit3, Cog2, Zmym5, Birc2, Atp6v0a1, Dnajb9, Parp14, Nsun3, Hspa13, Rpl3l, Akap8l, Slc22a12, 1700018L02Rik.

#### The target genes of transcription factor Phf8 are

Edem3, Phf8, Etv3, Rab25, Steap4, Aff1, Nr2c2, Hcst, Atg16l2, Usp47, Ttc13, Tsc22d1, Icam1, Myo1e, Zbtb38, Pik3r5, Atp6v0a1, Zfyve16, Pcnx, Myo10, Prm1, Ergic1, Dhx57, Slc41a1.

#### The target genes of transcription factor Xbp1 are

Tram2, Gpr55, Nek7, Dnm3, Mia3, 1110008P14Rik, Ubr3, Map1lc3a, Edem2, Slpi, Zbp1, Ssr4, Tmem154, Fam46c, Erp44, Ddost, Cpeb2, Srp72, Tesc, Bhlha15, Rpn1, Edem1, Gpr19, Kcnn4, Pld3, Fxyd5, Isg20, Cited2, Enpp1, Tspan15, Glipr1, Tmbim4, Mt3, Spcs1, Fndc3a, Dnajc3, Srpr, Rexo2, Ppib, Uba5, Manf, Xbp1, Cdc42se2, Gm2a, Evi2a, Slc35b1, Atp6v0a1, Ccr10, Wipi1, Pycr1, Gzma, Sdc1, Laptm4a, Pqlc3, Dhrs7, Fut8, Sel1l, Derl1, Tg, Grina, Trabd, Fkbp11, Itgb7, Prr13, Tnfrsf17, Eaf2, Tapbp, H2-Ke6, Epcam, Ndfip1, Cst6, Oosp1, Nfkb2.

#### The target genes of transcription factor Klf3 are

Cr2, Etv3, Usp53, Abca1, Heyl, Vamp1, Utrn, Tmcc3, Rasgrp3, 3110002H16Rik, Slc14a1.

#### The target genes of transcription factor Ebf1 are

Atp1b1, H3f3a, Ogt, Tmsb4x, Klf3, Igf1, Ebf1, H3f3b, Sox4, Gadd45g, Cplx2, Tspan13, H2-D1.

#### The target genes of transcription factor Myc are

Ephx1, Thnsl1, Car1, Hivep3, 5730409E04Rik, Trim62, Srm, Klhl8, Tpst1, Snx10, Aqp1, Snca, Mgll, Tnni3, Kcnn4, Nkg7, Trim3, Fut10, Nfix, Ccl17, Rab4a, Nrgn, Hyal2, Ccl5, Ccl4, Odc1, Tnfaip2, Myc, Cacna1i, Amigo2, Cela1, Ergic1, Tspo2.

#### The target genes of transcription factor Bach2 are

Rgs2, 4930523C07Rik, Atp1b1, Zeb2, Gpcpd1, Tifa, Bach2, Bcl7a, C1galt1, Cecr2, Prc1, Iqgap1, Arhgap17, Fam53b, Irf2, Cdkn3, Dnajc7, Myl4, Bptf, Nfkbia, Cerk, and Rsph1.

#### The target genes of transcription factor Tcf3 are

4930523C07Rik, Fcrla, Zeb2, Cytip, Tifa, Lmo4, Cecr2, Fam129c, Clint1, Myl4, Cplx2, Nfkbia, Foxn3, Cerk, and Snn.

#### The target genes of transcription factor Pax5 are

4930523C07Rik, Rcsd1, S1pr1, Bank1, Bach2, Pax5 itself, Zcchc11, Foxp1, Cotl1, Cyth1, Tnrc6b, Klhl24, and Arid1b.

#### The target genes of transcription factor Klf13 are

Stk17b, St8sia4, Ralgps2, 4930523C07Rik, Atp1b1, Fcrla, Gpcpd1, Samhd1, Ctsz, Tsc22d3, S100a11, Ppp3ca, Lmo4, Cd72, Smap2, Hvcn1, C1galt1, Fam3c, Cecr2, Pou2f2, Blvrb, Klf13 itself, Irf8, Cbfa2t3, Cdkn3, Dbnl, Peli1, Cdc42se2, Tnfrsf13b, Spns3, Msi2, Arl5c, Dnajc7, Sox4, Ly86, Dok3, Map3k1, H2-DMa, H2-Ab1, and H2-Aa.

#### The target genes of transcription factor Sox4 are

Atp1b1, Rcsd1, Fcrla, Zeb2, Rag1, Dstn, Ogt, Txnip, Tifa, Bach2, Pnrc1, Bcl7a, Foxp1, Mgst1, Ypel3, Fam53b, Myb, Marcks, Sec63, Vpreb3, Phip, Ebf1, Clint1, Gas7, Arl5c, Dnajc7, Sox4 itself, Cplx2, Tspan13, Nipbl, and Tcf4.

#### The target genes of transcription factor Foxp1 are

Clk1, Sp100, St8sia4, Ralgps2, 4930523C07Rik, Atp1b1, Fcrla, Arhgap30, Zeb2, Slc12a6, Dusp2, Ctsz, Ogt, Tmsb4x, Ash1l, S100a11, Ctss, Tifa, Lmo4, Lyn, Bach2, Pax5, Zcchc11, Smap2, Rhoh, Actb, Gimap1, Foxp1 itself, Blvrb, Akap13, Iqgap1, Ypel3, Pycard, Fam53b, Lsp1, Marcks, Sec63, Psap, Btg1, Irf2, Jund, Bnip3l, Cwf19l2, Pou2af1, Spg21, Rel, Gas7, Arl5c, Ikzf3, Dnajc7, Limd2, H3f3b, Tspan13, Nfkbia, Nipbl, Xrcc6, Arid1b, H2-Aa, H2-Eb1, Ltb, H2-Q7, and Sec11c.

#### The target genes of transcription factor Spib are

Clk1, Btg2, Ptprc, Ralgps2, 4930523C07Rik, Atp1b1, Fcrla, H3f3a, Zeb2, Cytip, Dstn, Ctsz, Ogt, Tsc22d3, Tmsb4x, Ash1l, S100a11, Lmo4, Lyn, Pax5, Krit1, Hvcn1, Bcl7a, Actb, C1galt1, Foxp1, Itpr2, Slc1a5, Pou2f2, Pafah1b3, Iqgap1, Sbk1, Marcks, Tcf3, Irf2, Jund, Klf2, Cbfa2t3, Prkcd, Bnip3l, Pou2af1, Rel, Clint1, Gm2a, Limd2, Bptf, Arid4b, Sox4, Jarid2, Dok3, Zfp36l1, Sec11c, and Ehd1.

#### The target genes of transcription factor Myb are

Zeb2, Lef1, Lmo4, Cplx2, and Pim1.

#### The target genes of transcription factor Klf2 are

Ralgps2, Sat1, Tmsb4x, Ctss, Txnip, Lmo4, Foxp1, Pou2f2, Blvrb, Arhgap17, Psap, Irf2, Jund, Ly86, and H2-T23.

#### The target genes of transcription factor Relb are

Rgs13, Dnajc1, Lmo2, Ganc, Cd40, Arhgap4, S1pr1, Srp72, Rpia, Relb, Nfkbid, Il4i1, Foxo3, Erich1, Junb, Pxk, Col5a3, Icam1, Sorl1, Myo1g, Lgals8, Cd83, Cd180, Gpr65, Btla, H2-DMb2, H2-T24, Klhl14, Gramd3, Ms4a4c, and Nfkb2.

#### The target genes of transcription factor Hivep3 are

Wnt10a, Chpf, Srpk3, Hivep3, Fosb, Nfkbid, Il4i1, Ccnd1, Pkib, Mfhas1, Smpd3, Egr3, Icam1, Cd83, Prr7, Cdc14b, Myc, St3gal1, Slc39a7, Klhl14, Setbp1, and Nfkb2.

#### The target genes of transcription factor Egr1 are

Dnajb2, Serpine2, Rgs13, Dusp10, Etl4, Cd40, Mllt3, Prdm2, Hspb1, Relb, Il4i1, Agbl1, Prmt2, Junb, Rab4a, Atxn7, Egr3, Tgfbr2, B230217C12Rik, Hdac9, Gxylt1, Pou6f1, and Egr1 itself.

#### The target genes of transcription factor Nfkb2 are

A530032D15Rik, Xpr1, Dusp10, Sec16a, Rasgrp1, Ganc, Hck, Pim2, Fam46c, Agl, Ddx58, Hivep3, Sfn, Gimap3, Exoc6b, Iqsec1, Relb, Apoe, Zfp36, Il4i1, Trim30b, Olfr482, Il4ra, Arid5b, Derl3, Stat6, Fcer2a, Extl3, Casp4, Myo1g, Ccr10, Hivep1, Cd83, Pcnx, Pla2g6, Btla, Fam120b, Fgd2, Nfkbie, Tmem63b, Crb3, Rasgrp3, Cd274, and Calhm2.

#### The target genes of transcription factor Hivep1 are

Tmem163, Cacna1e, Rag1, Cd40, Zmynd8, Il12a, Zfp296, Il4i1, Il16, Atg16l2, Swap70, Sbf2, Usp47, Slc12a3, Tk2, Col5a3, Spg21, Bcl2a1b, Gas7, Rnf167, Grn, Kdm1b, Trib2, Daam1, Calcoco1, Socs1, Gtpbp2, Camk2a, and Ehd1.

#### The target genes of transcription factor Cebpb are

Nacc2, Reln, Prdm1, Nol3, Pla2g6, and Mgat3.

#### The target genes of transcription factor Prdm1 are

Ctla4, Nacc2, Cebpb, Zbp1, Lmna, Dennd2d, Clec2g, and Cib2.

#### The target genes of transcription factor Runx2 include

Raph1, Pikfyve, Gmppa, Rgl1, Fcer1g, Mia3, Bmyc, Surf4, Pltp, Cybb, Irak1, Fndc3b, Notch2, Sec24d, Pink1, Isg15, Wfs1, Scarb2, Fam69a, Tpst2, Card11, Alox5ap, Tes, Zfp467, Rassf4, Kctd14, Nucb2, Plekha1, Ccdc6, Slc41a2, Dtx3, Sec24c, Rcbtb1, Tsc22d1, Hyou1, Anxa2, Scap, Snrk, Ccr9, Rnf215, Sertad2, Slfn5, Slfn8, Scpep1, Slc38a10, Fyb, Basp1, Cd200r1, Alcam, Crybg3, Gnptg, Sik1, Haao, Kcnk12, Map3k8, Sil1, Cep120, Slc22a12, Ms4a6c, Osbp, Mpeg1, Entpd1, and Fam45a.

#### The transcription factor Mef2c regulates the following genes

Stat1, Ier5, Rrbp1, Sdc4, Ctsz, Ctss, Smap2, Gimap1, Capg, Ptpn6, Ccnd2, Cd22, Il21r, Aldoa, Irf2, Cotl1, Irf8, Tsc22d1, Ets1, Rel, Bcl11a, Hexb, Cyth4, Tapbp, H2-DMb2, H2-Ab1, H2-Eb1, Ltb, Cd74, Ehd1, and Pdcd4.

#### The transcription factor Hmgb2 regulates the following genes

Atp1b1, H3f3a, Cdca8, Actb, Prc1, Cdk1, Btg1, Hmgb2, Cdkn3, Nrgn, Tmem108, Ebf1, Hist3h2a, Top2a, Hist1h1b, Hist1h1c, Cks2, and H2-D1.

### H Details of predicted regulons in bone marrow B cell states revealed through cell type-specific GRN analysis

#### The transcription factor Stat5a regulates the following genes

Tcf4, Cplx2, Cdkn1a, Prdx4, Tkt, Tmed2, Commd7, Eef1e1, Wnk1, Pds5a, and Txnl1.

#### The transcription factor Mef2c regulates the following genes

Hist1h2ae, Top2a, Cenpf, Ube2c, Cenpa, Nusap1, Hmgb2, Cdk1, Ucp2, Foxp1, and Smc4.

#### The transcription factor Ebf1 regulates the following genes

Vpreb2, Vpreb1, Igll1, 4930523C07Rik, Bfsp2, Slc25a1, Cd24a, Rhoh, Cd79b, Hhex, Cd69, Blnk, BC028528, Rbms1, Sox4, Pitpnc1, Ankfy1, Arntl, Tra2a, Cd79a, Tifa, Yif1b, Mxd4, Lrmp, Foxp1, Agpat3, Jun, Cd37, Smap2, Srp14, Fam53b, H2-Ob, Jmjd1c.

#### The transcription factor Pax5 regulates the following genes

Vpreb2, Vpreb1, Hoxa7, Igll1, Cd74, H2-Eb1, H2-Aa, Rgs18, Tcf4, Tpd52, 4930523C07Rik, H2-Ab1, Bfsp2, Med13l, Spib, Slc25a1, Cd24a, Rhoh, Mllt3, Luzp1, Cd69, Zfp36l2, Tbca, Blnk, BC028528, Pitpnc1, Serp1, Ankfy1, Rdx, Arntl, Tra2a, Cd79a, Tifa, Yif1b, Cotl1, Zeb1, Creg1, Mxd4, Lrmp, Arl5c, Foxp1, Dtnbp1, Agpat3, Cecr2, Jun, B2m, 1110008P14Rik, Myg1, Ogt, Smap2, Srp14, Fam53b, Arf6, Jmjd1c.

#### The transcription factor Myb regulates the following genes

Cenpf, Top2a, Hist1h2ae, Hmgb2, Mki67, Hist1h3c, Neil3, Hist1h3i, Ube2c, Hist2h2ac, Cenpa, Cdca7, Hist1h4h, Btg1, H2afv, Cd79a, Ctcf, Snrpf, Snrpc, Rdx, Acsl5, Lrmp, Ebf1, Hibadh, Pcbp2, Top1, Clk1, Zeb2, Gnb1, Ucp2, Hnrnpm, Foxp1, Myb, Ndufv3, Bcl7a, Fam53b, Ubap2l, Paip2.

#### The transcription factor Myc regulates the following genes

Tmem97, Psat1, Myc, Gcsh, Timm8a1, Srfbp1, Nop58, Gart, Naa10, Cdv3, Eif4e, Mrto4, Apex1, Ndufaf4, H2-K1, Fam162a, Nop56, Agpat5, Smn1, Polr2l, Eef1e1, Tomm5, Ebna1bp2, Mrps5, Rps19bp1, Prdx6, Nudt19, Lsm12, Txnrd3, Aprt, Aen, Nip7, Rpn1, Rsl24d1, Fxyd5, Prmt3, Glrx3, Eno1, Glrx5, Bola2, Pdcd5, Ptpn18, Txndc9, Ccnd2, Eif5b, Adh5, Gtf3c6, Phgdh, Tmed2, Psme1, Polr1d.

#### The transcription factor Bach2 regulates the following genes

Dnajc7, Atp1b1, Lmo4, Cdk6, Nusap1, Fcrla, Bach2, Rrm2, Cdk1, Sec63, Hist1h4d, Spns3, Vpreb3, Rgs2, Ebf1, Zeb2, Fam53b, Smc4, Sox4, Nfkbia, Tpd52, Dcaf7, Srpk2, Btg1, Cmc1, Actn4, Grcc10, Cnn3, Jund, Dcps, Etfb, Spcs3, Fkbp3, Mrps34, Myb, Cd79b, Tcf3, Pole4, Prcp, Atg3, Rtn3, Klhl24, Sae1, Snrpd1, Stag1, Hmgb2, Rab10, Hspa14, Calr, St13, Trim35, Pgk1, Prkar1a, Mtf2, Arid1a, Med30, Marcks, Pkn2, Capns1, Cd69, Eef2k, Nrd1, Foxp1, Selplg, Drap1, Rapgef6, Magoh, Ola1, Lsm3, Zmynd8, Ddah2, Srsf7, Cfdp1, Hp1bp3, Hspa5, Hsp90aa1, Ndufa9, Cnpy2, Usp15, Sdha, Calm1, Ndufb6, Stambpl1, Ctcf, Jun, Atf1, Dck, Slfn2, Dnaja2, H2-T23, Sbno1, Arntl, Chd4, Tpm4, Tmsb4x, Cdc5l, Eif3i, Brd2, Rae1, Gabpb1, Psmd12, Srp14, Actr3, Pdap1, Matr3, Rpl36a, Unc93b1, Hsp90b1, Diablo, Rplp1, Ddx39b, Akirin2, Napa, Cct8, Raly, Mocs2, Rab14.

#### The transcription factor Tcf3 regulates the following genes

Dnajc7, Ebf1, Btg1, Lgals1, Fkbp3, Spg21, Tcf3, Gon4l, Fkbp2, Clasp1, Ppp3cb, Zcchc7, Cisd2, Foxp1, Rapgef6, Hint1, Gapdh, Serf2, Ddx42, Sssca1, Slc25a11, Rbm39, Chd2.

#### The transcription factor Klf2 regulates the following genes

Cd52, Hmgb1, Klf3, Ralgps2, Cd2, Cd79b, Tmsb4x, Tsc22d3, Smap2, Zfp706, Wasf2, Tcp11l2, Jund, Farsb, Sec61g, H2-K1, H2-D1, Sat1, and Arid1a.

#### The transcription factor Ctcf regulates the following genes

Fcrl1, Ctss, Nasp, H2afv, Gimap6, Vpreb3, Rnaseh2a, Dbi, Exosc3, Ppp1cb, Mrpl34, Cox7a2, Fcrla, Coro1a, Tcp1, Ndufc2, Hnrnpm, Smap2, Ubn1, Pdcd4, Tomm7, Nup50, Dmxl1, Cox6a1, Aco2, Ssr3, Zc3h13, Ufm1, Fam103a1, Mapkap1, Mrps12, Prkar1a, Cmpk1, H2-D1, Mapk1, Ptp4a2, Cuedc2, Stk4, Tomm22, Arhgap17, Trappc10, Dctn6, and Cct5.

**Supplementary Table 9:** EPR of Non-specific Chip-seq with top 500 most varying genes. Values below the performance of random predictors have been excluded from the table.

|  | hHep | mDC | mHSC-E | mHSC-L | hESC | mESC | mHSC-GM |
| --- | --- | --- | --- | --- | --- | --- | --- |
| HyperVAE | 3.49 | 3.57 | 6.61 | 3.75 | 2.30 | 3.32 | 7.89 |
| DeepSEM | 2.50 | 2.92 | 6.59 | 3.12 | 2.20 | 2.79 | 7.22 |
| PIDC | 2.48 | 2.73 | 5.41 | 2.85 | - | - | - |
| GENIE3 | - | - | - | - | - | 3.25 | 6.68 |
| GRNBoost2 | - | - | - | 2.85 | 1.52 | - | - |
| SCODE | - | - | - | - | - | 1.66 | 1.25 |
| ppcor | 1.10 | 1.01 | 1.90 | 1.05 | - | - | - |
| SINCERITIES | - | - | - | - | 1.98 | - | - |

**Supplementary Table 10:**
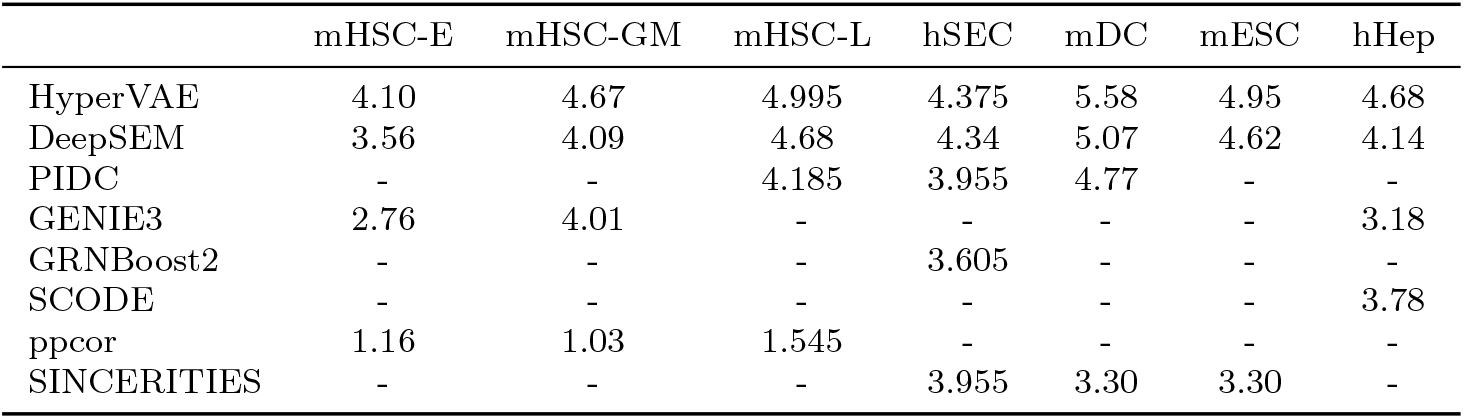
AUPRC of Non-specific Chip-seq with top 500 most varying genes. Values below the performance of random predictors have been excluded from the table.

|  | mHSC-E | mHSC-GM | mHSC-L | hSEC | mDC | mESC | hHep |
| --- | --- | --- | --- | --- | --- | --- | --- |
| HyperVAE | 4.10 | 4.67 | 4.995 | 4.375 | 5.58 | 4.95 | 4.68 |
| DeepSEM | 3.56 | 4.09 | 4.68 | 4.34 | 5.07 | 4.62 | 4.14 |
| PIDC | - | - | 4.185 | 3.955 | 4.77 | - | - |
| GENIE3 | 2.76 | 4.01 | - | - | - | - | 3.18 |
| GRNBoost2 | - | - | - | 3.605 | - | - | - |
| SCODE | - | - | - | - | - | - | 3.78 |
| ppcor | 1.16 | 1.03 | 1.545 | - | - | - | - |
| SINCERITIES | - | - | - | 3.955 | 3.30 | 3.30 | - |

**Supplementary Table 11:** EPR of STRING with top 1000 most varying genes. Values below the performance of random predictors have been excluded from the table.

|  | mHSC-E | mHSC-GM | mHSC-L | hSEC | mDC | mESC | hHep |
| --- | --- | --- | --- | --- | --- | --- | --- |
| HyperVAE | 5.53 | 3.71 | 8.52 | 9 | 7.74 | 2.38 | 4.26 |
| DeepSEM | 4.49 | 2.96 | 7.57 | 8.26 | 7.31 | 1.94 | 3.89 |
| PIDC | 4.12 | 3.62 | 8.08 | - | - | - | - |
| GENIE3 | - | - | - | 8.49 | 7.17 | 2.16 | - |
| GRNBoost2 | - | - | - | - | - | - | 3.71 |
| SCODE | - | - | 3.94 | 1.43 | - | - | - |
| ppcor | 1.09 | 1.09 | - | - | - | - | - |
| SINCERITIES | - | - | - | - | 1.43 | 1.09 | 1.24 |

**Supplementary Table 12:** AUPRC of STRING with top 1000 most varying genes. Values below the performance of random predictors have been excluded from the table.

|  | mHSC-E | mHSC-GM | mHSC-L | hSEC | mDC | mESC | hHep |
| --- | --- | --- | --- | --- | --- | --- | --- |
| HyperVAE | 2.49 | 1.96 | 1.48 | 2.16 | 6.53 | 7.47 | 7.48 |
| DeepSEM | 2.10 | 1.68 | 1.41 | 2.22 | 5.44 | 6.28 | 7.33 |
| PIDC | 2.01 | 1.90 | 1.58 | 2.06 | 5.09 | 6.27 | - |
| GENIE3 | - | - | - | - | - | 6.27 | - |
| GRNBoost2 | 1.67 | - | - | - | - | - | 7.07 |
| SCODE | - | - | - | 1.15 | 2.09 | 1.3 | - |
| ppcor | - | - | - | - | - | - | - |
| SINCERITIES | - | 1.04 | 1.01 | 1.15 | - | - | 1.04 |

### I The list of ten gene clusters predicted by the gene encoder of HyperG-VAE based on bone marrow B cells

**Supplementary Table 13:** AUPRC of STRING with top 500 most varying genes. Values below the performance of random predictors have been excluded from the table.

|  | hESC | hHep | mDC | mESC | mHSC-E | mHSC-GM | mHSC-L |
| --- | --- | --- | --- | --- | --- | --- | --- |
| HyperVAE | 2.3 | 2.0 | 1.64 | 2.17 | 6.35 | 7.35 | 7.14 |
| DeepSEM | 2.04 | 1.74 | 1.58 | 2.02 | 5.77 | 6.98 | 7.22 |
| PIDC | 1.95 | 1.92 | 1.57 | 1.89 | 4.8 | - | - |
| GENIE3 | - | - | - | - | - | 6.83 | 6.88 |
| GRNBoost2 | - | - | - | - | - | - | - |
| SCODE | - | - | - | - | - | - | - |
| ppcor | 1.02 | - | - | - | - | - | - |
| SINCERITIES | - | 1.14 | 1.02 | 1.09 | - | - | - |

**Supplementary Table 14:** EPR of Non-specific Chip-seq with top 1000 most varying genes. Values below the performance of random predictors have been excluded from the table.

|  | mHSC-L | mESC | hHep | mDC | mHSC-GM | mHSC-E | hESC |
| --- | --- | --- | --- | --- | --- | --- | --- |
| HyperVAE | 3.92 | 3.71 | 3.2 | 3.6 | 6.49 | 6.76 | 2.43 |
| DeepSEM | 3.3 | 3.25 | 2.71 | 3.39 | 6.03 | 6.08 | 2.32 |
| PIDC | - | - | 2.44 | 2.95 | 5.85 | 5.74 | - |
| GENIE3 | - | - | - | - | 5.85 | - | - |
| GRNBoost2 | 3.08 | 3.51 | - | - | - | - | 0 |
| SCODE | 1.38 | - | - | - | - | - | - |
| ppcor | 1.38 | 1.19 | 1.25 | 1.15 | 1.17 | - | 1.28 |
| SINCERITIES | - | - | - | - | - | 1.22 | 2.13 |

**Supplementary Table 15:** EPR of STRING with top 500 most varying genes. Values below the performance of random predictors have been excluded from the table.

|  | mDC | hSEC | hHep | mHSC-E | mHSC-GM | mHSC-L | mESC |
| --- | --- | --- | --- | --- | --- | --- | --- |
| HyperVAE | 2.22 | 4.8 | 3.75 | 8.13 | 9.4 | 7.58 | 3.75 |
| DeepSEM | 1.96 | 4.13 | 2.92 | 7.43 | 9.03 | 6.98 | 3.38 |
| PIDC | - | 3.75 | 3.51 | 7.49 | - | - | - |
| GENIE3 | 2.03 | - | - | - | 8.65 | 6.83 | - |
| GRNBoost2 | - | - | - | - | - | - | 3.35 |
| SCODE | - | - | - | - | - | 1.97 | 1.54 |
| ppcor | - | - | 1.04 | 1.96 | 1.58 | - | - |
| SINCERITIES | 1.03 | 1.11 | - | - | - | - | - |

**Supplementary Table 16:** AUPRC of Non-specific Chip-seq with top 1000 most varying genes. Values below the performance of random predictors have been excluded from the table.

|  | mESC | mDC | mHSC-L | mHSC-E | mHSC-GM | hHep | hSEC |
| --- | --- | --- | --- | --- | --- | --- | --- |
| HyperVAE | 1.63 | 1.8 | 3.42 | 3.24 | 3.91 | 1.47 | 1.27 |
| DeepSEM | 1.62 | 1.8 | 2.73 | 3.13 | 3.38 | 1.39 | 1.25 |
| PIDC | - | 1.65 | 2.91 | 2.65 | - | - | - |
| GENIE3 | 1.66 | - | - | - | 3.56 | - | - |
| GRNBoost2 | - | - | - | - | - | 1.09 | 1.02 |
| SCODE | - | - | - | 1.75 | 1.14 | 1.35 | - |
| ppcor | - | - | 1.01 | - | - | - | - |
| SINCERITIES | 1.13 | 1.1 | - | - | - | - | 1.17 |

**Supplementary Table 17:** AUPRC of Cell-type-specific Chip-seq with top 500 most varying genes. Values below the performance of random predictors have been excluded from the table.

| Method | hESC | mDC | hHep | mESC | LOF/GOF mESC | mHSC-GM | mHSC-E | mHSC-L |
| --- | --- | --- | --- | --- | --- | --- | --- | --- |
| HyperVAE | 1.37 | 1.46 | 1.25 | 1.05 | 1.33 | 1.06 | 1.06 | 1.08 |
| DeepSEM | 1.13 | 1.10 | 1.24 | 1.05 | 1.27 | 1.04 | 1.02 | 1.07 |
| PIDC | - | - | 1.05 | 1.02 | 1.24 | - | - | - |
| GENIE3 | 1.0 | 1.01 | 1.10 | 1.06 | - | - | 1.01 | - |
| GRNBoost2 | - | 1.01 | 1.05 | - | - | - | - | 1.02 |
| SCODE | - | - | - | - | 1.14 | 1.01 | 1.06 | - |
| ppcor | 1. | - | - | - | - | - | - | - |
| SINCERITIES | - | 1.14 | - | 1.02 | - | 1.03 | - | 1.06 |

**Supplementary Table 18:** EPR of Cell-type-specific Chip-seq with top 500 most varying genes. Values below the performance of random predictors have been excluded from the table.

| Method | hHep | mESC | hESC | mDC | mHSC-GM | mHSC-E | Lof/gof mESC | mHSC-L |
| --- | --- | --- | --- | --- | --- | --- | --- | --- |
| HyperVAE | 1.19 | 1.08 | 1.68 | 1.85 | 1.09 | 1.06 | 1.42 | 1.13 |
| DeepSEM | 1.19 | 1.04 | 1.19 | 1.17 | 1.12 | 1.05 | 1.37 | 1.06 |
| PIDC | 1.03 | - | - | - | - | - | - | - |
| GENIE3 | 1.12 | 1.06 | - | 1.04 | 1.06 | 1.03 | 1.36 | - |
| GRNBoost2 | - | - | - | - | - | 1.03 | - | 1.07 |
| SCODE | - | - | - | - | - | 1.03 | - | 1.07 |
| ppcor | - | 1.01 | 1.02 | - | - | - | - | - |
| SINCERITIES | - | - | - | 1.31 | 1.01 | 1.02 | - | - |

### In gene cluster 0, the following genes are included

Tram2, Ccdc150, Ankrd44, Wnt10a, Serpine2, Sp140, Gigyf2, Gm7967, Ubxn4, Xpr1, 4930523C07Rik, F5, Sh2d1b1, Fcgr3, Arhgap30, B930036N10Rik, Ifi209, Ephx1, Wdr26, Degs1, Gm15867, Thnsl1, Nacc2, Sec16a, Zeb2, Cytip, Rapgef4, C1qtnf4, Rag1, Ganc, Ccndbp1, Pdia3, Dusp2, Zc3h6, Smox, Lrrn4, Gm14167, Slc9a8, Zbp1, Zfp968, Ppdpf, Pcmtd2, B630019K06Rik, Xlr4c, Srpk3, Irak1, Gm5127, Gm15261, Fndc3b, Mgarp, Siah2, Slc33a1, Tmem154, S100a6, S100a8, S100a11, Cd160, Pde4dip, Notch2, Pifo, Cd53, Agl, Usp53, Sec24d, Mcub, Zfp292, Tgfbr1, Col27a1, Jun, Cdc20, Hivep3, 5730409E04Rik, Fuca1, C1qa, Tnfrsf18, Cdk14, Cacna2d1, Fgl2, Kmt2e, Cenpa, Gm9903, Acox3, Gm42726, Rhoh, Rbm47, Aff1, Lrrc8c, Ficd, Bcl7a, Tmem248, Dnaaf5, Gng11, C1galt1, Impdh1, E330009J07Rik, Trbc1, Zfp467, Gimap6, Gadd45a, Igkv2-116, Igkv2-109, Igkv12-46, Igkv6-32, Dqx1, Nagk, A430078I02Rik, Arhgap25, H1fx, Gm20696, Vgll4, Rasgef1a, Usp18, Vamp1, Bhlhe41, Zfp787, Bbc3, Vasp, Apoe, Dmac2, B3gnt8, Spint2, Klk1, Atf5, Med25, Rras, Agbl1, Sema4b, Cib1, Usp35, Trim3, Ampd3, Coq7, Rgs10, Mki67, Ifitm2, Ccnd1, Tpcn2, Akap12, Gm10827, 4933404K13Rik, Prep, Vsir, Arid5b, Upb1, Gstt2, Agpat3, Zfr2, Zfp433, Slc41a2, Ckap4, Igf1, Dgka, Fcer2a, Coprs, Nek3, Ankrd37, Spcs3, Calr, Slc12a3, Herpud1, Tk2, Nol3, Map10, Fhit, Sec24c, Glt8d1, Mmp14, Rcbtb1, Ephx2, Loxl2, Fndc3a, Rb1, Rubcnl, Epsti1, Gpr18, Cwf19l2, Casp4, Tirap, Stt3a, Hyou1, Sik3, Rexo2, Pou2af1, 2010007H06Rik, Stoml1, Dennd4a, Usp3, Ccpg1, Tmem30a, Ctsh, Rasa2, Srprb, Uba5, Manf, Nradd, Snrk, Slc6a20b, Ccr9, Ccr2, Xbp1, Sertad2, Acyp2, Erlec1, Grap, Aldoc, Ift20, Evi2a, Gm11205, Ccl5, Slc35b1, Atp6v0a1, Kcnh6, Abca5, Kif19a, Smim5, Trim65, Gaa, Tcrg-C4, Ripor2, Fam8a1, Ctsl, Msh3, Erbin, Gm15326, Fam228b, Rhob, Laptm4a, Nbas, Trib2, Odc1, Grhl1, Gdap10, Klhdc1, Fut8, Tmed10, Samd15, Ighg2c, Ighg2b, Ighv7-3, Ighv1-36, Ighv1-64, Gm5441, Capsl, Prlr, Gm34590, Them6, Ly6k, Zfp623, Grina, Pla2g6, AL591952.3, Gm26822, Sdf2l1, Tmem191c, Gramd1c, Cd200, St3gal6, Speer2, Hspa13, Ifnar2, Cbr1, Fam120b, Hagh, Gnptg, Zfp563, H2-Ob, Neu1, H2-Q6, Atat1, Ppp1r10, H2-T23, Gabbr1, AY036118, Crisp3, Tspo2, Tgif1, Gm26637, Prkce, Rprd1a, Ammecr1l, Map3k2, Sil1, Ap3s1, Cep120, Gm26742, Gm4951, Tcf4, Ctsw, Rom1, Slc15a3, Fas, Nfkb2, Ccdc186, Fam45a, Tmlhe, AC125149.3.

#### In gene cluster 1, the following genes are included

Mcmdc2, Creg2, Cd28, Ctla4, Sgpp2, Gpr55, Tmem37, C4bp, F730311O21Rik, Cacna1s, Dnm3, Tbx19, Fcrla, Ifi211, Ifi203, Ifi205, H3f3a, Etl4, Gm13610, Gm35202, Gpr21, Nr6a1, Gm13561, Hoxd3os1, Prg2, Lrp4, Mdk, Chac1, Cst3, a, 9230111E07Rik, Atp9a, Srms, Fndc11, Uckl1os, Gm10489, Xlr, Slc6a8, Ssr4, Tmsb4x, Serp1, Cd5l, Rab25, Krtcap2, Gm15417, Rps27, Gm15265, Fam46c, Cd101, Gm5547, Tifa, Lmo4, Cd72, AI427809, Trim62, Cd52, Cnksr1, Extl1, C1qb, Camk2n1, Tmem82, Fbxo2, Morn1, AW011738, Gm8879, Kdr, Cxcl10, Plac8, Cryba4, Tesc, Ccdc92, Cldn4, Hspb1, C130050O18Rik, Actb, Bhlha15, Ica1, Klrg2, Gm32479, Igkv2-137, Igkv9-129, Igkv17-121, Igkv13-84, Igkv4-59, Igkv4-57, Igkv4-55, Igkv5-43, Igkv5-39, Igkv8-30, Igkv6-29, Igkv6-25, Igkv8-24, Igkv6-23, Igkv8-21, Igkv6-17, Igkv6-15, Igkv3-12, Igkv3-10, Igkv3-5, Igkv3-1, Igkc, Thnsl2, Tmem150a, Tmsb10, Lrig1, Foxp1, Zfp9, Ninj2, Clec2g, Gm47861, Mansc1, Gm15510, Meis3, A930016O22Rik, Gm16174, Cd79a, Pafah1b3, E130208F15Rik, Upk1a, Siglecg, Napsa, Gm45552, Cd37, Ftl1, Kctd21, Spcs2, Dnajb13, Nupr1, Ypel3, Gm15533, Gm44623, Lsp1, Cd24a, Prdm1, Tspan15, Gstt1, Mif, Derl3, Chchd10, Vpreb3, Gm49322, Hsp90b1, Btg1, Kcnmb4os2, Eef1akmt3, Il23a, Hmgb2, Crlf1, Jund, Klf2, 2210011C24Rik, Mt3, Gm31805, Tsnaxip1, Cyba, Ear2, AC160336.1, Carmil3, Nuggc, Gm29642, Kcnj1, H2afx, Pclaf, Ppib, Filip1, Gm39383, Pls1, Gm19325, Ryk, Gm47328, P4htm, Tmppe, Eml6, Ebf1, Gm16033, Rasgef1c, Gpx3, Pld2, Wscd1, Dusp14, Tbx21, Arl5c, Top2a, Cnp, Dnajc7, Ccr10, Cyb561, Limd2, Gm10840, Cd79b, Hid1, H3f3b, Gm11754, Metrnl, Hist1h2ae, Gm31834, Wnk2, Gm47918, Gm48899, Ell2, Mef2c, F2rl1, Marveld2, Gm48684, Ankrd55, Gzma, Gm10734, 5430401H09Rik, Gm36756, Ifi27l2a, Igha, Adam6b, Ighv2-2, Ighv5-4, Ighv5-6, Ighv2-9-1, Ighv5-17, Ighv2-9, Ighv11-2, Ighv6-3, Ighv6-6, Ighv10-1, Ighv1-5, Ighv1-12, Ighv1-15, Ighv1-22, Ighv1-53, Ighv1-55, Ighv8-8, Ighv1-59, Ighv1-63, Ighv1-69, Ighv8-12, Snhg18, Gm35167, Tg, Ly6d, Ly6e, Apol9a, Lgals1, Kdelr3, Mgat3, A4galt, Abcd2, Fkbp11, Gm21917, AC191865.2, Tnfrsf17, Igll1, Vpreb1, Iglc1, Iglc3, Iglc2, Iglv3, Iglv2, Itgb5, Sidt1, Gcsam, Ripply3, Gm26753, Cacna1h, Tead3, H2-K1, BC051142, H2-D1, Gm42418, Gm19585, Prr22, Crb3, Xdh, Epcam, Mzb1, Nrg2, Spinkl, Cd74, Sec11c, Tubb6, Gal, Cst6, Malat1, Frmd8os, Syt7, Oosp2, Oosp1, Dmrt3, Cpeb3, mt-Co1, mt-Atp6, mt-Co3, mt-Nd4, mt-Cytb, AC168977.1.

**Supplementary Table 19:** EPR of Cell-type-specific Chip-seq with top 1000 most varying genes. Values below the performance of random predictors have been excluded from the table.

| Algorithm | LOF/GOF mESC | hESC | mDC | mHSC-L | hHep | mESC | mHSC-E | mHSC-GM |
| --- | --- | --- | --- | --- | --- | --- | --- | --- |
| HyperVAE | 1.44 | 1.52 | 1.27 | 1.16 | 1.19 | 1.11 | 1.09 | 1.08 |
| DeepSEM | 1.42 | 1.18 | 1.05 | 1.12 | 1.22 | 1.07 | 1.06 | 1.1 |
| PIDC | - | - | - | 1.01 | 1.01 | - | - | - |
| GENIE3 | 1.35 | - | 1.01 | - | 1.12 | 1.06 | 1.01 | 1.03 |
| GRNBoost2 | - | - | - | 1.04 | 1.01 | - | - | - |
| SCODE | - | - | - | 1.08 | 1.01 | - | - | - |
| ppcor | 1.03 | 1.05 | - | - | - | - | - | - |
| SINCERITIES | - | 1.65 | 1.3 | - | - | - | - | - |

**Supplementary Table 20:** AUPRC of Cell-type-specific Chip-seq with top 1000 most varying genes. Values below the performance of random predictors have been excluded from the table.

| Algorithm | mHSC-GM | mHSC-E | mHSC-L | LOF/GOF mESC | hHep | mDC | mESC | hESC |
| --- | --- | --- | --- | --- | --- | --- | --- | --- |
| HyperVAE | 1.01 | 1.08 | 1.16 | 1.34 | 1.16 | 1.12 | 1.09 | 1.31 |
| DeepSEM | 1.01 | 1.01 | 1.11 | 1.33 | 1.25 | 1.03 | 1.09 | 1.16 |
| PIDC | - | - | - | 1.23 | 1.03 | - | 1.02 | - |
| GENIE3 | - | - | 1.03 | 1.23 | 1.08 | 1.01 | 1.06 | - |
| GRNBoost2 | - | 1. | 1.03 | - | 1.03 | 1.01 | - | - |
| SCODE | 1. | 1. | 1.06 | 1.16 | - | - | - | - |
| ppcor | 1. | 1. | - | - | - | - | - | - |
| SINCERITIES | - | - | - | - | - | 1.11 | 1.02 | 1.6 |

#### In gene cluster 2, the following genes are included

4732440D04Rik, Neurl3, Tsga10, Inpp1, Raph1, Kansl1l, Sp100, Itm2c, Hdlbp, St8sia4, Rgs2, Cop1, Rcsd1, Gm15853, Pcp4l1, Fcer1g, Psen2, Dusp10, Kctd3, G0s2, Camk1g, Cr2, Dnajc1, Qrfp, Lcn2, Gad1, Gpr155, A330069E16Rik, Lmo2, Cd59a, Nusap1, Gm14005, 2900093K20Rik, Sdc4, Araf, Cfp, Akap17b, Ogt, Taf9b, Tceal1, Tiparp, Ctso, Lmna, Gm43714, Trim46, Pbxip1, Bank1, Ppp3ca, Gbp5, 2210414B05Rik, Lyn, Bach2, Erp44, Mknk1, Macf1, Ddost, Prkcz, Krit1, Reln, Insig1, Cpeb2, Bst1, Hopx, Jchain, Scarb2, Hsd17b11, Tpst2, Gm42903, Tmem120b, Hip1r, Ncf1, Muc3a, Gpc2, Alox5ap, D730045B01Rik, Creb3l2, Gm28053, Gimap7, Gimap1, Snca, Igkv15-103, Igkv10-94, Igkv19-93, Igkv8-28, Gm30211, Eif2ak3, Capg, Aup1, Pcyox1, Rpn1, Mgll, Necap1, Cd27, Prkcg, Zfp296, Relb, Kcnn4, Phldb3, Ceacam1, Tmem91, Blvrb, Zfp59, Tyrobp, Hcst, Nfkbid, Scn1b, Spib, Chd2, AU020206, Pde8a, Il16, Rnf169, Trim5, Trim12c, Gvin1, Smg1, Tmc7, Gm45184, Cox6a2, Ifitm3, Utrn, 1700027J07Rik, Adamts14, Reep3, Rhobtb1, Ccdc6, Gadd45b, Nfic, Gm17745, Os9, Ddit3, Stat6, Zbtb39, Cnpy2, Fut10, Mfhas1, 1700029J07Rik, March1, Gmip, Bst2, Colgalt1, Cib3, Rrad, Fuk, Gse1, Gm45890, Cog2, Atxn7, Zswim8, Arhgef3, Txndc16, Bnip3l, Egr3, Rgcc, Dnajc3, Fbxl12, Gm47079, Yipf2, Bbs9, Gm26787, Sorl1, Gm47232, Jaml, 1700017B05Rik, Gm26609, 4930429F24Rik, Fam46a, Uba7, Shisa5, Camp, Gm9856, Myo1g, Bcl11a, Hba-a1, Wwc1, Tgtp1, Mgat1, Rapgef6, Pik3r5, Inca1, Slfn5, Pgap3, Ccr7, Vat1, Pecam1, Mif4gd, Olfr1369-ps1, Zscan26, Hist1h1c, Gm47730, Tmem170b, Susd3, Syk, Zfp759, Clptm1l, Serinc5, Hdac9, Strn3, Zfyve26, Pcnx, Arel1, Ift43, Tex22, Ighm, Ighv14-2, Ighv9-3, Gm30948, Ighv1-82, Basp1, Otulin, Ankrd33b, Ncf4, AL590144.2, Cyth4, H1f0, Cbx7, Grap2, Sgsm3, Zc3h7b, Bik, Trabd, AC158554.1, Lmbr1l, Tuba1c, Prpf40b, Smagp, Ciita, 2510002D24Rik, St6gal1, Rubcn, Alcam, Nxpe3, Ermard, Zfp945, Ift140, Pim1, Pde9a, Gm19412, H2-DMa, Ltb, Slc25a27, B230354K17Rik, Tmem63b, Haao, Rhoq, Fbxo11, Ddx3y, Cables1, Klhl14, Egr1, Pcdhgb4, Iigp1, Gm9949, Setbp1, Ap5b1, Gm14964, Ifit3, Zfyve27, Ablim1, CAAA01118383.1.

#### In gene cluster 3, the following genes are included

Gm16152, Gm20342, Wnt6, Irs1, Mroh2a, Ramp1, 5033417F24Rik, Farp2, Lax1, Rgs13, Serpinc1, Gm37065, Ifi214, Ifi213, Phyh, Sfmbt2, Enkur, Lcn4, Prrc2b, Tor2a, Hspa5, Mettl5os, Pdk1, Zdhhc5, Ptprj, Pex16, B230118H07Rik, Slc12a6, Cpxm1, Ddrgk1, Slc23a2, Gpcpd1, Rrbp1, Hck, Map1lc3a, Samhd1, Slpi, Ctsz, Gm14403, Gm14325, Gm14327, Gdi1, Med12, Gla, Sat1, 1700125G22Rik, Larp1b, Mgst2, Ssr3, Il12a, Etv3, Glmp, Ash1l, Hist2h4, Pigk, Tmem245, Gm26566, Lepr, Bend5, Slc5a9, Gm13031, Gm13075, Galnt11, Tmem214, Uvssa, Gm45495, Klf3, Txk, Ppbp, Pf4, Fam109a, Rhof, Rilpl1, Card11, Akr1b10, Gimap5, 5430402O13Rik, Snx10, Tacstd2, Igkv14-126-1, Igkv12-44, Gm45051, Ggcx, Spr, Kbtbd12, Clec4a3, Clec4d, Gpr162, Gm15987, Cd69, Klrd1, Klra7, Eps8, Mgst1, Slc1a5, Arhgef1, Zfp260, Cd22, Cebpg, Gm26526, Il4i1, Fam169b, Isg20, Fchsd2, Olfr655, Sbf2, Gga2, Il4ra, Il21r, Sbk1, Sec23ip, Nt5dc1, Aifm2, Lss, Itgb2, Zbtb7a, Lyz2, Gm32235, Cpm, Rab5b, A430078G23Rik, Aga, Fam129c, Tmem38a, Ier2, Adcy7, Adgrg1, Tldc1, Dennd6a, Prkcd, Sh3bp5, Pck2, Nynrin, Zmym5, Phf11b, Ctsb, Elp3, Klf12, Dock9, Col5a3, Gm47230, Arid3b, Lipc, 4933433G15Rik, Slc17a5, Hyal2, Amigo3, Ngp, Arpp21, Fbxl2, Zkscan7, Lztfl1, Adam19, Hs3st3b1, Sox15, C1qbp, Unc119, Heatr6, Fbxo47, Tubg2, Rundc1, Nbr1, Acbd4, Myl4, Wipi1, Tmc6, Rnf213, Pycr1, Gm26601, Hist1h4d, Cmah, Pxdc1, Gm29458, Jarid2, Mylip, Gadd45g, Prr7, Fam193b, Agtpbp1, Gm36445, Mccc2, Ccnb1, Gm21762, Rrm2, Gm9887, Fos, Sel1l, Foxn3, Serpina3f, Ighv5-16, Ighv3-6, Ighv1-52, Ighv1-77, Cmbl, Derl1, Mtss1, Tnfrsf13c, Prr5, Cerk, Creld2, Amigo2, Pou6f1, Mgrn1, Vpreb2, AC140186.1, Parp14, Eaf2, Ccr6, Mapk8ip3, AI413582, Gm15420, Rsph1, Sik1, H2-DMb2, H2-DMb1, H2-Q7, Rasgrp3, 4833418N02Rik, Ston1, Gm26734, 3110002H16Rik, 4930426D05Rik, Camk2a, Gnal, Stard6, Clcf1, Klc2, Slc3a2, Fads2, Ms4a6c, Smarca2, 4430402I18Rik, Cd274, Sufu, Mirt1, AC149090.1.

#### In gene cluster 4, the following genes are included

Gm16152, Gm20342, Wnt6, Irs1, Mroh2a, Ramp1, 5033417F24Rik, Farp2, Lax1, Rgs13, Serpinc1, Gm37065, Ifi214, Ifi213, Phyh, Sfmbt2, Enkur, Lcn4, Prrc2b, Tor2a, Hspa5, Mettl5os, Pdk1, Zdhhc5, Ptprj, Pex16, B230118H07Rik, Slc12a6, Cpxm1, Ddrgk1, Slc23a2, Gpcpd1, Rrbp1, Hck, Map1lc3a, Samhd1, Slpi, Ctsz, Gm14403, Gm14325, Gm14327, Gdi1, Med12, Gla, Sat1, 1700125G22Rik, Larp1b, Mgst2, Ssr3, Il12a, Etv3, Glmp, Ash1l, Hist2h4, Pigk, Tmem245, Gm26566, Lepr, Bend5, Slc5a9, Gm13031, Gm13075, Galnt11, Tmem214, Uvssa, Gm45495, Klf3, Txk, Ppbp, Pf4, Fam109a, Rhof, Rilpl1, Card11, Akr1b10, Gimap5, 5430402O13Rik, Snx10, Tacstd2, Igkv14-126-1, Igkv12-44, Gm45051, Ggcx, Spr, Kbtbd12, Clec4a3, Clec4d, Gpr162, Gm15987, Cd69, Klrd1, Klra7, Eps8, Mgst1, Slc1a5, Arhgef1, Zfp260, Cd22, Cebpg, Gm26526, Il4i1, Fam169b, Isg20, Fchsd2, Olfr655, Sbf2, Gga2, Il4ra, Il21r, Sbk1, Sec23ip, Nt5dc1, Aifm2, Lss, Itgb2, Zbtb7a, Lyz2, Gm32235, Cpm, Rab5b, A430078G23Rik, Aga, Fam129c, Tmem38a, Ier2, Adcy7, Adgrg1, Tldc1, Dennd6a, Prkcd, Sh3bp5, Pck2, Nynrin, Zmym5, Phf11b, Ctsb, Elp3, Klf12, Dock9, Col5a3, Gm47230, Arid3b, Lipc, 4933433G15Rik, Slc17a5, Hyal2, Amigo3, Ngp, Arpp21, Fbxl2, Zkscan7, Lztfl1, Adam19, Hs3st3b1, Sox15, C1qbp, Unc119, Heatr6, Fbxo47, Tubg2, Rundc1, Nbr1, Acbd4, Myl4, Wipi1, Tmc6, Rnf213, Pycr1, Gm26601, Hist1h4d, Cmah, Pxdc1, Gm29458, Jarid2, Mylip, Gadd45g, Prr7, Fam193b, Agtpbp1, Gm36445, Mccc2, Ccnb1, Gm21762, Rrm2, Gm9887, Fos, Sel1l, Foxn3, Serpina3f, Ighv5-16, Ighv3-6, Ighv1-52, Ighv1-77, Cmbl, Derl1, Mtss1, Tnfrsf13c, Prr5, Cerk, Creld2, Amigo2, Pou6f1, Mgrn1, Vpreb2, AC140186.1, Parp14, Eaf2, Ccr6, Mapk8ip3, AI413582, Gm15420, Rsph1, Sik1, H2-DMb2, H2-DMb1, H2-Q7, Rasgrp3, 4833418N02Rik, Ston1, Gm26734, 3110002H16Rik, 4930426D05Rik, Camk2a, Gnal, Stard6, Clcf1, Klc2, Slc3a2, Fads2, Ms4a6c, Smarca2, 4430402I18Rik, Cd274, Sufu, Mirt1, AC149090.1.

#### In gene cluster 5, the following genes are included

Gm16152, Gm20342, Wnt6, Irs1, Mroh2a, Ramp1, 5033417F24Rik, Farp2, Lax1, Rgs13, Serpinc1, Gm37065, Ifi214, Ifi213, Phyh, Sfmbt2, Enkur, Lcn4, Prrc2b, Tor2a, Hspa5, Mettl5os, Pdk1, Zdhhc5, Ptprj, Pex16, B230118H07Rik, Slc12a6, Cpxm1, Ddrgk1, Slc23a2, Gpcpd1, Rrbp1, Hck, Map1lc3a, Samhd1, Slpi, Ctsz, Gm14403, Gm14325, Gm14327, Gdi1, Med12, Gla, Sat1, 1700125G22Rik, Larp1b, Mgst2, Ssr3, Il12a, Etv3, Glmp, Ash1l, Hist2h4, Pigk, Tmem245, Gm26566, Lepr, Bend5, Slc5a9, Gm13031, Gm13075, Galnt11, Tmem214, Uvssa, Gm45495, Klf3, Txk, Ppbp, Pf4, Fam109a, Rhof, Rilpl1, Card11, Akr1b10, Gimap5, 5430402O13Rik, Snx10, Tacstd2, Igkv14-126-1, Igkv12-44, Gm45051, Ggcx, Spr, Kbtbd12, Clec4a3, Clec4d, Gpr162, Gm15987, Cd69, Klrd1, Klra7, Eps8, Mgst1, Slc1a5, Arhgef1, Zfp260, Cd22, Cebpg, Gm26526, Il4i1, Fam169b, Isg20, Fchsd2, Olfr655, Sbf2, Gga2, Il4ra, Il21r, Sbk1, Sec23ip, Nt5dc1, Aifm2, Lss, Itgb2, Zbtb7a, Lyz2, Gm32235, Cpm, Rab5b, A430078G23Rik, Aga, Fam129c, Tmem38a, Ier2, Adcy7, Adgrg1, Tldc1, Dennd6a, Prkcd, Sh3bp5, Pck2, Nynrin, Zmym5, Phf11b, Ctsb, Elp3, Klf12, Dock9, Col5a3, Gm47230, Arid3b, Lipc, 4933433G15Rik, Slc17a5, Hyal2, Amigo3, Ngp, Arpp21, Fbxl2, Zkscan7, Lztfl1, Adam19, Hs3st3b1, Sox15, C1qbp, Unc119, Heatr6, Fbxo47, Tubg2, Rundc1, Nbr1, Acbd4, Myl4, Wipi1, Tmc6, Rnf213, Pycr1, Gm26601, Hist1h4d, Cmah, Pxdc1, Gm29458, Jarid2, Mylip, Gadd45g, Prr7, Fam193b, Agtpbp1, Gm36445, Mccc2, Ccnb1, Gm21762, Rrm2, Gm9887, Fos, Sel1l, Foxn3, Serpina3f, Ighv5-16, Ighv3-6, Ighv1-52, Ighv1-77, Cmbl, Derl1, Mtss1, Tnfrsf13c, Prr5, Cerk, Creld2, Amigo2, Pou6f1, Mgrn1, Vpreb2, AC140186.1, Parp14, Eaf2, Ccr6, Mapk8ip3, AI413582, Gm15420, Rsph1, Sik1, H2-DMb2, H2-DMb1, H2-Q7, Rasgrp3, 4833418N02Rik, Ston1, Gm26734, 3110002H16Rik, 4930426D05Rik, Camk2a, Gnal, Stard6, Clcf1, Klc2, Slc3a2, Fads2, Ms4a6c, Smarca2, 4430402I18Rik, Cd274, Sufu, Mirt1, AC149090.1.

#### In gene cluster 6, the following genes are included

Clk1, Fam117b, Pikfyve, Tmem163, Rgl1, 4930439D14Rik, Sell, Gm13383, Man1b1, Ak8, Ak1, Wipf1, Lpcat4, Rasgrp1, Gm10762, H2al1m, Rhox8, Yipf6, Il2rg, Magt1, Tsc22d3, Car1, Rnf13, Fcrl1, 4933434E20Rik, Gm43573, Pax5, Akap2, Tm2d1, Eps15, Zdhhc18, Casp9, Per3, 1500002C15Rik, Samd11, Nub1, Mxd4, Grk4, Uchl1, Stap1, Gm32051, Oasl2, Hvcn1, Gusb, Samd9l, Bet1, Irf5, Igkv1-135, Igkv1-133, C87436, Sec61a1, Cecr2, Cd4, Ccnd2, Klrb1c, Klre1, Plbd1, Itpr2, Isoc2b, Fosb, Rabac1, Ryr1, Fxyd5, Selenos, Prc1, Myo7a, Atg16l2, Arap1, Hbb-bt, Trim30b, Arhgap17, Aldoa, Ifitm1, Pkp3, Tnnt3, Cttn, Gm26740, Sesn1, Prf1, 4930507D05Rik, Gm867, Zfp280b, Pofut2, Fgd6, Kcnmb4, Gm16553, Erich1, 5830468F06Rik, Hook3, Vps37a, Inpp4b, Dnase2a, Mt1, Ctrl, Irf8, Rab4a, Ccser2, Gpr137c, Lgals3, Lpar6, Tmem123, Birc2, Birc3, Olfr889, Gm26737, Hykk, Ccnb2, Bcl2a1b, 4930524O07Rik, Rbm5, Cdhr4, Gas2l1, Aff4, Igtp, Epn2, Gas7, Cldn7, Slfn2, Ccl9, Scpep1, Ormdl3, Gh, Ern1, Abca6, Narf, Arid4b, Hist1h1b, Hist1h2ap, Sox4, Ly86, Txndc5, Hivep1, Rgs14, Txndc15, Zfp457, Pqlc3, Pdia6, Tspan13, Daam1, Dhrs7, Susd6, Lgmn, Ighv1-19, Ighv1-81, Selenop, Myo10, Stk3, Gm49085, Sla, Lynx1, C1qtnf6, Rpl39l, 3110001I22Rik, B3gnt5, Gp5, D930030I03Rik, Arid1b, Rpl3l, Fam234a, Crebrf, Fgd2, Tapbp, H2-Oa, H2-Ab1, H2-Eb2, Nfkbie, Zfp318, Treml2, Hnrnpll, Pcdhb16, Alpk2, Neat1, Ahnak, AW112010, Ms4a4c, Mpeg1, Entpd1, 2310034G01Rik, Calhm2.

#### In gene cluster 7, the following genes are included

Fam135a, Lmbrd1, 4930403P22Rik, Stat1, Stk17b, Mogat1, Bcl2, Slc35f5, Cd55, Nek7, Trove2, Gm10138, Trmt1l, Creg1, Cd84, Opn3, C8g, Cir1, Zfp120, Ninl, Tbc1d20, Dusp15, Pltp, Zmynd8, 3830403N18Rik, Smim10l2a, Rab39b, Anxa5, Foxo1, Kcnab1, Slc50a1, S100a3, Mcl1, Txnip, Cd2, S1pr1, Ifi44, Manea, Slc44a1, Bspry, Aknaos, Mllt3, Faah, Smap2, Cdca8, Pink1, Wfs1, Sel1l3, Lrrc8d, Golga3, Slc15a4, Zfp113, Igkv5-48, Gfpt1, Etfrf1, Gm15873, Zfp329, Pglyrp1, Pou2f2, Pld3, Zfp36, Rasip1, Nav2, Siglech, Ints4, Usp47, Lmntd2, Chst3, Cnn2, Arhgap45, Tcf3, Tle2, Lrp1, Letm2, Plpp5, Micu3, Cope, Il12rb1, Junb, Ccl17, Coq9, Slc12a4, Terf2ip, Cbfa2t3, Samd8, Gm26772, Ppp3cc, Mbnl2, Casp1, Srpr, Anxa2, Rnf111, Adam10, Zbtb38, Peli1, Psme4, Hba-a2, Clint1, 9930111J21Rik2, Hist3h2a, Chrnb1, Wfdc21, Synrg, Msi2, Plekhh3, Lgals8, Serpinb1a, Cks2, 1810034E14Rik, Tmem161b, Gm47551, Map3k1, Slc38a9, Noxred1, Ifi27, Ckb, Ighv1-26, Dennd3, Ly6a, Xrcc6, Gxylt1, Snn, Txndc11, Dnajb11, Filip1l, Ergic1, H2-Aa, Gm26917, Nrtn, Ankrd12, Dhx57, Arhgef33, Kcnk12, Pkd2l2, Ndfip1, Gramd3, Siglec15, 1700018L02Rik, Dntt, Scd1.

#### In gene cluster 8, the following genes are included

A530040E14Rik, Insig2, Tor3a, Gm26620, Capn2, Mia3, Nebl, Surf4, Tmem87a, Slc28a2, Edem2, Cd40, Zfp973, Pim2, Cybb, Cysltr1, Skil, Zbtb7b, Ube2j1, Ddx58, Reck, Heyl, Zmym6, Prdm2, 5031425E22Rik, Hpse, Fam69a, Gm14508, Ubc, Asns, Tmem106b, 1110019D14Rik, Tes, Fam3c, Bpgm, Braf, Igkv14-111, Exoc6b, Cpne9, Rassf4, Dyrk4, Dennd5b, Ncr1, Zfp865, Zfp324, Plekhf1, Pex11a, Kctd14, Trim34a, Trim30a, Nucb2, Sec63, Gucd1, Dusp6, Glipr1, B4galnt1, Arhgef18, Galnt7, Rab3a, B3gnt9, Carmil2, Ctsg, Ptk2b, Gm4285, Rab3d, Nrgn, Il10ra, Zc3h12c, Spg21, Myo1e, Trim7, Sec24a, Trim11, Xaf1, Mir142hg, 2610035D17Rik, Slc38a10, Myadml2, Nid1, Prss16, C530050E15Rik, Irf4, Cd83, Kdm1b, Nxnl2, Slc34a1, Zfyve16, Cd180, Dnajb9, Fam177a, Nfkbia, Gm10457, Gpr65, Ccdc88c, Serpina3g, Adssl1, Pld4, Fyb, Arfgap3, Calcoco1, 2010309G21Rik, Olfr166, Klhl24, Hes1, Iqcb1, Nlrc4, Map3k8, Gm17227, Ms4a6b, Frat1, Tcf7l2, Shtn1.

#### In gene cluster 9, the following genes are included

Cacna1e, Tmem164, Serpini1, Gba, Gm31243, Pnrc1, B4galt1, Gm12678, Zcchc11, Cited4, Id3, Ddi2, Mib2, Tbc1d1, Gbp9, Oasl1, Gm43409, Mlxip, Zfp12, Daglb, Igkv9-120, Iqsec1, Gpr19, Leng8, Zfp773, Zfp719, Akap13, Stard10, Pycard, Tspan32, Man1a, Ddx21, Rufy2, Cyp4f18, Nfix, Icam1, Gm27201, Hyal3, Lars2, Dok3, Gm34215, Ighv5-2, Shisa8, Nr4a1, Itgb7, Brwd1, Gtpbp2, Tnfsf9, Uty.

### J Original results of BEELINE benchmark comparison

Original performance on GRN inference of HyperG-VAE based on the setting of BEELINE framework [36] can be found in Supplementary Table 9-20.

## Footnotes

1 In gene regulatory networks (GRNs) analysis, gene modules refer to clusters of genes that are regulated together by the same set of transcription factors.

